# Lysosomal cystine export regulates mTORC1 signaling to guide kidney epithelial cell fate specialization

**DOI:** 10.1101/2022.08.28.505580

**Authors:** Marine Berquez, Zhiyong Chen, Beatrice Paola Festa, Patrick Krohn, Svenja Aline Keller, Silvia Parolo, Mikhail Korzinkin, Anna Gaponova, Endre Laczko, Enrico Domenici, Olivier Devuyst, Alessandro Luciani

**Affiliations:** Institute of Physiology, University of Zurich, 8057 Zurich, Switzerland; Fondazione The Microsoft Research University of Trento (Centre for Computational and Systems Biology (COSBI), 38068 Rovereto, Italy; Insilico Medicine Hong Kong Ltd., Hong Kong Science and Technology Park, Hong Kong, Hong Kong SAR, China; Functional Genomics Center Zurich, University of Zurich, Winterthurerstrasse 190, CH-8057, Zurich, Switzerland; Department of Cellular, Computational and Integrative Biology (CIBIO), University of Trento, 38123 Trento, Italy; Institute for Rare Diseases, UC Louvain Medical School, 1200 Brussels, Belgium

## Abstract

Differentiation is critical for cell fate decisions, but the signals involved remain unclear. The kidney proximal tubule (PT) cells reabsorb disulphide-rich proteins through endocytosis, generating cystine via lysosomal proteolysis. Here we report that defective cystine mobilization from lysosomes through cystinosin (CTNS), which is mutated in cystinosis, diverts PT cells towards growth and proliferation, disrupting their functions. Mechanistically, cystine storage stimulates Ragulator-Rag GTPase-dependent recruitment of mechanistic target of rapamycin complex 1 (mTORC1) and its constitutive activation. Re-introduction of CTNS restores nutrient-dependent regulation of mTORC1 in knockout cells, whereas cell-permeant analogues of L-cystine, accumulating within lysosomes, render wild-type cells resistant to nutrient withdrawal. Therapeutic mTORC1 inhibition corrects lysosome and differentiation downstream of cystine storage, and phenotypes in a zebrafish model of cystinosis. Thus, cystine serves as a lysosomal signal that tailors mTORC1 and metabolism to direct epithelial cell fate decisions. These results identify mechanisms and therapeutic targets for dysregulated homeostasis in cystinosis.

## Introduction

The epithelial cells lining the kidney proximal tubule (PT) play a critical role in homeostasis by efficiently reabsorbing small proteins from the primary urine. The process is mediated by receptor-mediated endocytosis and lysosomal pathways, preventing the loss of precious nutrients in the urine (1). The lysosomal degradation of disulphide-rich plasma proteins like albumin accounts for the supply of cystine that is exported to cytoplasm through the lysosomal cystine transporter cystinosin (2). In the cytoplasm, cystine is reduced to cysteine and utilized for the maintenance of cellular redox homeostasis (3). Additionally, the lysosome-mediated removal of dysfunctional mitochondria regulates the differentiating state of PT cells through junctional integrity and transcriptional programs, including the transcription factor Y box 3 (Ybx3) (4). Recessive mutations in *CTNS*, the gene that encodes cystinosin (CTNS), cause cystinosis, a lysosomal storage disease characterized by loss of endocytosis/reabsorptive properties of PT cells, progressing towards kidney failure (5). How, mechanistically, the increased levels of cystine within lysosomes, reflecting its defective export through CTNS, affect cell fate decisions remains unknown.

A crucial step in nutrient sensing is the recruitment of the evolutionarily conserved mechanistic target of rapamycin complex 1 (mTORC1) to the surface of the lysosome (6). In presence of nutrients, such as amino acids, glucose, and lipids, a multiprotein complex composed of heterodimeric Rag (Ras-related GTP binding) GTPases (consisting of RagA or RagB bound to RagC or RagD), Ragulator, and vacuolar H^+^-ATPase (V-ATPase) tethers mTORC1 to the lysosomal surface, enabling its activation by the (growth factor-induced) GTP-binding protein Rheb (Ras homolog enriched in brain; refs. 7,8). The mTORC1 signaling from lysosomes promotes growth and proliferation while repressing lysosome biogenesis and catabolic autophagy (9). The proper regulation of mTORC1 sustains polarized transport properties in epithelial cell types (10). This has been well illustrated in the kidney, where hypo and/or hyperactivation of mTORC1 pathway can drive pathological changes in podocytes and tubular cells, whereas blunting of hyperactive mTORC1 may prove beneficial in renal cell carcinoma and polycystic and/or diabetic kidney disease (11).

Like other lysosomal amino acid transporters, the CTNS protein physically interacts with the V-ATPase-Ragulator-Rag GTPase lysosomal scaffold that activates mTORC1 (12). During periods of starvation in *Drosophila*, mobilization of cystine from lysosomes fuels acetyl-CoA synthesis and limits TORC1 reactivation, altogether balancing catabolic autophagy and growth in developing organs (13). In the mammalian kidney, PT cells are particularly vulnerable to imbalances in metabolism, which can curb their differentiation and cause disease (14). These findings raise the possibility that lysosomal cystine export may shape the response of mTORC1 and metabolism to fate decisions during the differentiation of the kidney tubule epithelium.

Here, we combine genetic model organisms (mouse, rat, and zebrafish) and physiologically relevant cell culture systems to test the causal role of an evolutionarily conserved pathway that tunes mTORC1 signaling and metabolism in response to levels of lysosomal cystine to instruct fate specialization. Together the results provide insights into the mechanisms that regulate homeostasis and specialized function in kidney epithelial cells, and therapeutic targets for cystinosis.

## Results

### Cystine storage diverts differentiation towards proliferation and dyshomeostasis

The genetic deletion of *Ctns* in 24-week-old mouse kidneys induces cystine storage and lysosome abnormalities in PT segments (Fig. 1a,b), leading to defective receptor-mediated endocytosis of low molecular weight proteins (LMWPs; Fig. 1c) and their losses in the urine (Fig. 1d). Other markers of kidney tubular dysfunction, such as albuminuria and glucosuria, were significantly higher in *Ctns* knockout (KO) mice when compared with wild-type (WT) littermates (<u>Supplementary Fig. 1a,b</u>), confirming previous studies (5,16,17). The expression of genes involved in the tubular reabsorption of albumin (*Lrp2*), glucose (*Slc5a2*), and LMWP/CC16 (*Cubn*) inversely correlated with solute loss in *Ctns*^KO^ compared with *Ctns*^WT^ animals (<u>Supplementary Fig. 1c-e</u>), reinforcing the link between loss of differentiation markers (i.e., transport systems) and development of kidney tubular dysfunction. Analysis of the transcriptome of PT segments from 24-week-old littermates (Fig. 1e; <u>Supplementary Fig. 1f</u>) identified 61 genes differentially expressed between *Ctns*^KO^ and *Ctns*^WT^ mice, of which 43 were upregulated and 18 downregulated, respectively. Gene set enrichment analysis revealed an over-representation of pathways involved in the cell cycle regulation, whereas membrane trafficking, vesicle-mediated/small molecule transport, and hemostasis were significantly under-represented (Fig. 1f). The dysregulation of these pathways was verified (<u>Supplementary Fig. 1g</u>), with increased expression of proteins regulating cell cycle (Ccna2, Cyclin A2, and Cdc20, Cell cycle division 20) and proliferation (Pcna, Proliferating cell nuclear antigen), contrasting with decreased levels of Lrp2 (LDL receptor related protein 2)/megalin, the master coordinator of LMWP uptake (17), in the PT of *Ctns*^KO^ kidneys compared with *Ctns*^WT^ littermates (Fig. 1g,h; <u>Supplementary Fig. 1h</u>). In line, decreased expression of cell-differentiation related genes (*Slc5a2* and *Slc34a1*; ref. 14) and increased (de)differentiation markers (Foxm1, Sox9, and Vimentin; refs. 18,19) were observed in the PTs of 24-week-old *Ctns*^KO^ mouse kidneys compared with *Ctns*^WT^ littermates (Fig. 1g,i; <u>Supplementary Fig. 1i</u>). Consistent with altered tubular differentiation programs, we noted that the PT cells from *Ctns*^KO^ mice changed from a cuboidal shape to a flattened and spread-out epithelium (Fig. 1j). Thus, the cystine build-up secondary to the loss of CTNS triggers anabolic programs for growth and proliferation, disrupting the differentiating state of the PT cells and their reabsorptive properties.

**Figure 1.**
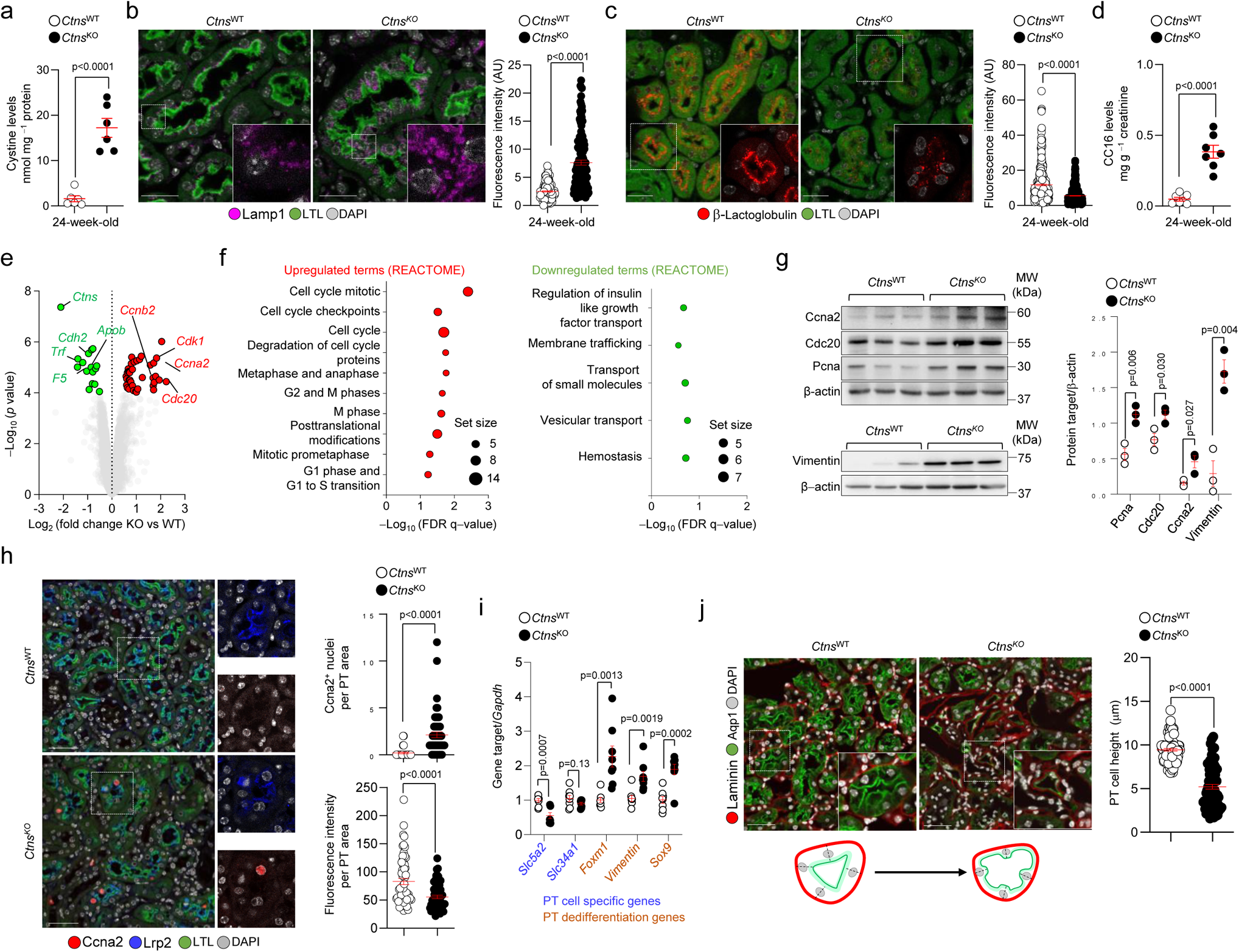
Cystine storage divert differentiation towards growth and proliferation. (**a**) Cystine levels in *Ctns* mouse kidneys; n=6 animals per group. (**b**) Immunofluorescence and quantification of the Lamp1 mean fluorescence intensity (MFI) in LTL^+^ PTs of *Ctns* kidneys, n>137 PTs pooled from three mice. (**c**) Confocal microscopy and quantification of the Cy5-tagged β-lactoglobulin MFI in LTL^+^ PTs of *Ctns* kidneys, n>330 PTs pooled from three mice. (**d**) CC16 levels in the urine samples from *Ctns* mice; n=7 animals per group. (**e**) Volcano plot of genome-wide changes in PT segments from *Ctns* kidneys. Red and green dots show the differentially expressed genes (DEGs; adjusted *p* value < 0.05; n=3 animals per group). (**f**) GSEA of over- and under-represented biological pathways in the PT segments of *Ctns*^KO^ mouse kidneys. (**g**) Immunoblotting and quantification of the indicated proteins in *Ctns* kidneys; n=3 animals per group. (**h**) Confocal microscopy of whole-kidney sections stained for Ccna2 (red), Lrp2 (blue), and PT marker (LTL, green); Quantification of Ccna2^+^ nuclei per PT: n>31 PTs and Lrp2 MFI: n>56 PTs pooled from three mice. (**i**) mRNA levels for the indicated genes; n=8 replicates pooled from 3 biologically independent experiments. (**j**) Whole-kidney sections stained for PT apical marker (Aqp1, green) and PT basolateral marker (Laminin, red). Confocal microscopy and quantification of PT cell height in *Ctns* kidneys; n=90 PTs pooled from 3 mice per each group. Plots represent mean ± SEM. Statistics calculated by unpaired two-tailed Student’s *t* test. Dotted white squares contain images at high magnification. Nuclei counterstained with DAPI (grey). Scale bars are 50μm in **b**, **c**, and **h**, and 20μm in **j**. Unprocessed scans of original blots shown in Supplementary Fig. 14. Source data are provided as a Source Data file.

### Defective lysosomes lead to tubular cell fate changes

In silico predictions of gene function and module associations (GeneBridge toolkit; www.systems-genetics.org; ref. 20) suggested that the lysosomally localized *Ctns* relates to lytic vacuole and organelle pathways beyond cystine transport (<u>Supplementary Fig. 2a-c</u>). This led us to investigate whether changes in the differentiation and tubular cell fate induced by CTNS loss and cystine storage reflect alterations in the lysosomal catabolic activities. We used primary cells derived from microdissected PT segments of mouse *Ctns* kidneys (referred to as mPTCs; Ref. 4) and transduced them with an adenovirus expressing the lysosomal transmembrane protein 192 (Tmem192) fused to 3×haemagglutinin (T192-3×HA; ref. 21; Fig. 2a) to enable the immunoisolation and functional probing of pure and intact lysosomes (<u>Supplementary Fig. 2d-h</u>). The *Ctns*^KO^ mPTCs, which accumulate cystine within lysosomes (Fig. 2b), showed abnormal growth/proliferation rates (Fig. 2c), defective endocytosis of fluorescent-tagged ligands (Fig. 2d), and changes in the expression of PT cell (de)differentiation markers (Fig. 2e), closely mimicking the changes observed *in vivo*. Moreover, these cells showed abnormal/elongated primary cilium − an antenna-like sensory organelle whose signaling regulates differentiation and cell fate (22) − when compared with WT cells (Fig. 2f). These changes were paralleled by a marked decrease in the numbers of functionally degradative lysosomes in *Ctns*^KO^ mPTCs, both under full and nutrient-depleted conditions, when compared to *Ctns*^WT^ cells (<u>Supplementary Fig. 3a-c</u>). In agreement, the *Ctns*^KO^ cells showed higher numbers of Map1lc3b-positive autophagic puncta and autolysosomes, insensitive to treatment with Bafilomycin A1 (Fig. 2g; <u>Supplementary Fig. 3d</u>). Comparative analysis of lysosome proteins from *Ctns*^KO^ versus *Ctns*^WT^ mPTCs revealed decreased proteolytic generation of the 32 kDa luminal mature cathepsin D (CtsD) and augmented levels of the lipidated, autophagosome-associated form Map1Lc3b-II (Fig. 2h). Similar alterations were also observed in *Ctns*^KO^ mouse kidneys (<u>Supplementary Fig. 3e</u>), as well as in the *ctns*-deficient zebrafish (4) and *Ctns*^KO^ rat model (23), both displaying significant cystine storage, metabolic defects, and PT cell dedifferentiation and tubular dysfunction (<u>Supplementary Fig. 3f-i</u>). The proteolytic defects were not linked to dysregulated lysosomal acidification as judged by the measurements in *Ctns* mPTCs transiently expressing a bona fide biosensor (R2pH-LAMP1-3×HA; ref. 24; <u>Supplementary Fig. 4a-d</u>) that readily measures the changes in luminal pH of the lysosome or stained with LysoTracker (<u>Supplementary Fig. 4a-d)</u> (25). Thus, the tubular cells lacking CTNS/ accumulating cystine reprogramme their metabolism from catabolism to anabolism to fuel growth and proliferation, leading to dysregulated differentiation and dyshomeostasis.

**Figure 2.**
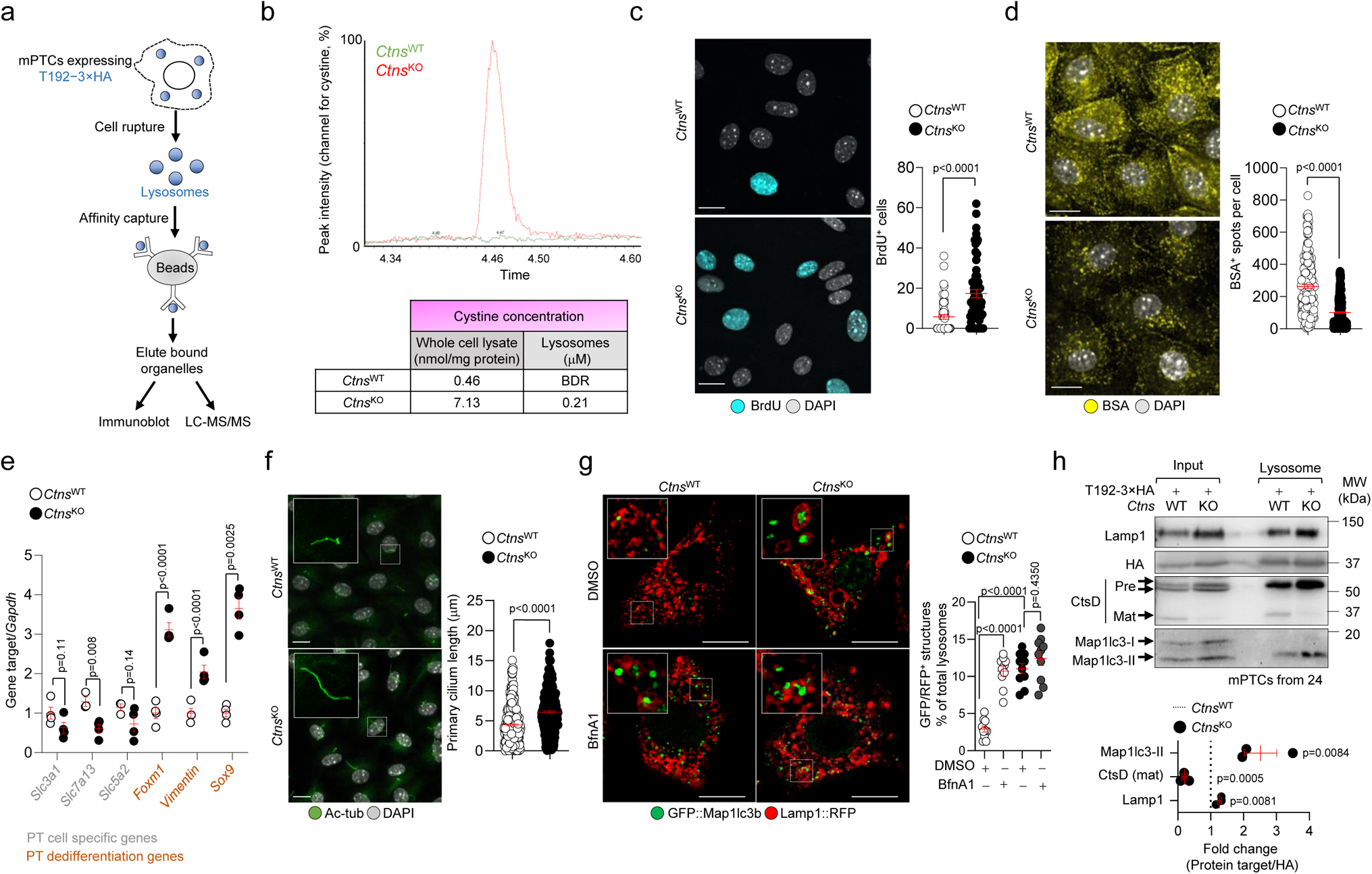
Defective lysosomes lead to loss of differentiation and dyshomeostasis. (**a**) Schematic showing lysosome purification using affinity-based capture from cells transiently expressing T192-3×HA. LC–MS/MS, liquid chromatography with tandem mass spectrometry. (**b**) Quantification of cystine levels in the lysosomes purified from *Ctns* mPTCs; n=2 biologically independent experiments per group. (**c**) Confocal microscopy and quantification of BrdU^+^ cells (expressed as percentage of total cells; n>55 randomly selected fields of views per each condition pooled from 3 biologically independent experiments). (**d**) Confocal microscopy and quantification of BSA^+^ structures per cell (n>175 cells per each condition pooled from 3 biologically independent experiments). (**e**) mRNA levels for the indicated genes; n>3 replicates pooled from 3 biologically independent experiments. (**f**) Maximum intensity projection of image stacks and quantification of the primary cilium length (Ac-tubulin, green) in *Ctns* mPTCs; n>285 cells pooled from 4 biologically independent experiments. (**g**) Quantification of GFP/RFP^+^ structures in *Ctns* cells transiently expressing GFP::Map1lc3b and Lamp1::RFP (expressed as percentage of total lysosomes; n=10 randomly selected fields of views per each condition). (**h**) Immunoblots and quantification of the indicated proteins in input samples and corresponding Lyso-IP fractions from *Ctns* mPTCs; n=3 biologically independent experiments. Plots represent mean ± SEM. Statistics calculated by unpaired two-tailed Student’s *t* test in **c**, **d**, **e**, **f** and **h** or by one-way ANOVA followed by Sidak’s multiple comparisons test in in **g**. Dotted white squares contain images at high magnification. Nuclei counterstained with DAPI (grey). Scale bars are 10μm. Unprocessed scans of original blots shown in Supplementary Fig. 14. Source data are provided as a Source Data file.

### Hyperactive mTORC1 signalosome drives the metabolic switch

To identify the factors that drive the metabolic switch induced by CTNS deficiency, we conducted proteomics- and metabolomics-based profiling of *Ctns* mPTCs (Fig. 3a). PCA and hierarchical clustering revealed that proteome and metabolome of *Ctns*^KO^ mPTCs substantially differ from the *Ctns*^WT^ control cells (Fig. 3b,c; <u>Supplementary Fig. 5a-b</u>). Protein-protein interaction analysis of differentially expressed proteins revealed significant enrichment in metabolism, cell cycle and ribosome biogenesis, growth and proliferation, apoptosis regulation, signal transduction, and differentiation pathways in *Ctns*^KO^ versus *Ctns*^WT^ mPTCs (<u>Supplementary Fig. 5c,d</u>). Metabolite set enrichment analysis (MSEA) identified in *Ctns*^KO^ cells chemical categories predominantly containing amino acids, organic acids, sugars, and nucleotides (<u>Supplementary Fig. 5e-g</u>). Integrated biological network analysis crossing differentially produced proteins with metabolites (Fig. 3b,c) identified pathways linked to regulation of the mTORC1 signalosome, such as glucose and amino acids (6), mTORC1 signaling itself, and downstream anabolic (protein, fatty acid, and nucleotide biosynthesis) or catabolic (autophagy) processes enriched in *Ctns*^KO^ mPTCs. An enrichment in pathways related to cancer cell metabolism and renal cell carcinoma, known to be driven by aberrant mTORC1 signaling (26), was also observed in these cells (Fig. 3d). These results were confirmed when applying an artificial intelligence (AI) engine (PandaOmics; ref. 27) that uses machine learning (ML) tools and statistical validation to rank disease-target associations and prioritize actionable drug targets. Accordingly, this platform identified mTOR on the top of the list of predicted drug targets (<u>Supplementary Fig. 6</u>) potentially reversing dyshomeostasis and tubular dysfunction downstream of cystine storage. Inspired by the biological evidence and in silico predictions, we next examined the mTORC1 signaling in the CTNS-deficient PT cells. The CTNS deficiency rendered mTORC1 less sensitive to nutrient starvation and constitutively active, as indicated by persistent phosphorylation of the ribosomal protein S6 (pS6^Ser235/236^) and eukaryotic translation initiation factor 4E (eIF4E)-binding protein 1 (p4EBP1^Ser65^) (Fig. 3e). Increased phosphorylation rates of the canonical mTORC1 substrate (pS6^Ser235/236^) were also observed in *Ctns*^KO^ mouse kidneys (<u>Supplementary Fig. 7a,b</u>). Prolonged starvation led to a partial and progressive reactivation of mTORC1 in wild-type cells, while this process was severely impaired in starved *Ctns*^KO^ mPTCs, as measured by phosphorylation of the mTORC1 substrate S6 (<u>Supplementary Fig. 7c</u>). Increasing the cytosolic cysteine levels through the supplementation of non-toxic concentrations of L-cysteine (1mM for 16h; <u>Supplementary Fig. 7d</u>) or modified molecules such as N-acetylcysteine (NAC; 1mM for 16h; <u>Supplementary Fig. 7e</u>) did not blunt the hyperactive mTORC1 signaling in *Ctns*^KO^ PT cells (<u>Supplementary Fig. 7f</u>), suggesting that the recycling of lysosomal cystine through CTNS regulates mTORC1 signaling in the tubular cells. In agreement, CTNS deficiency augmented the lysosomal levels of components (28) of the V-ATPase subunits Atp6v0d1 and Atp6v1b2, Ragulator complex subunit Lamtor2 and RagC GTPase that control mTORC1 activation (Fig. 3f). In *Ctns*^KO^ mPTCs, RagC GTPase was strongly clustered on Lysosomal-associated membrane protein 1 (Lamp1)-positive structures compared to *Ctns*^WT^ mPTCs (Fig. 3g), substantiating the regulatory effect of CTNS and cystine storage on mTORC1 activation. The impact of CTNS loss on mTORC1 signaling was further examined in *ctns*-deficient zebrafish (4) and *Ctns*^KO^ rat model (23), both displaying increased phosphorylation rates of pS6^Ser235/236^ compared to controls (Fig. 3h-j), demonstrating the evolutionary conservation of this connection.

**Figure 3.**
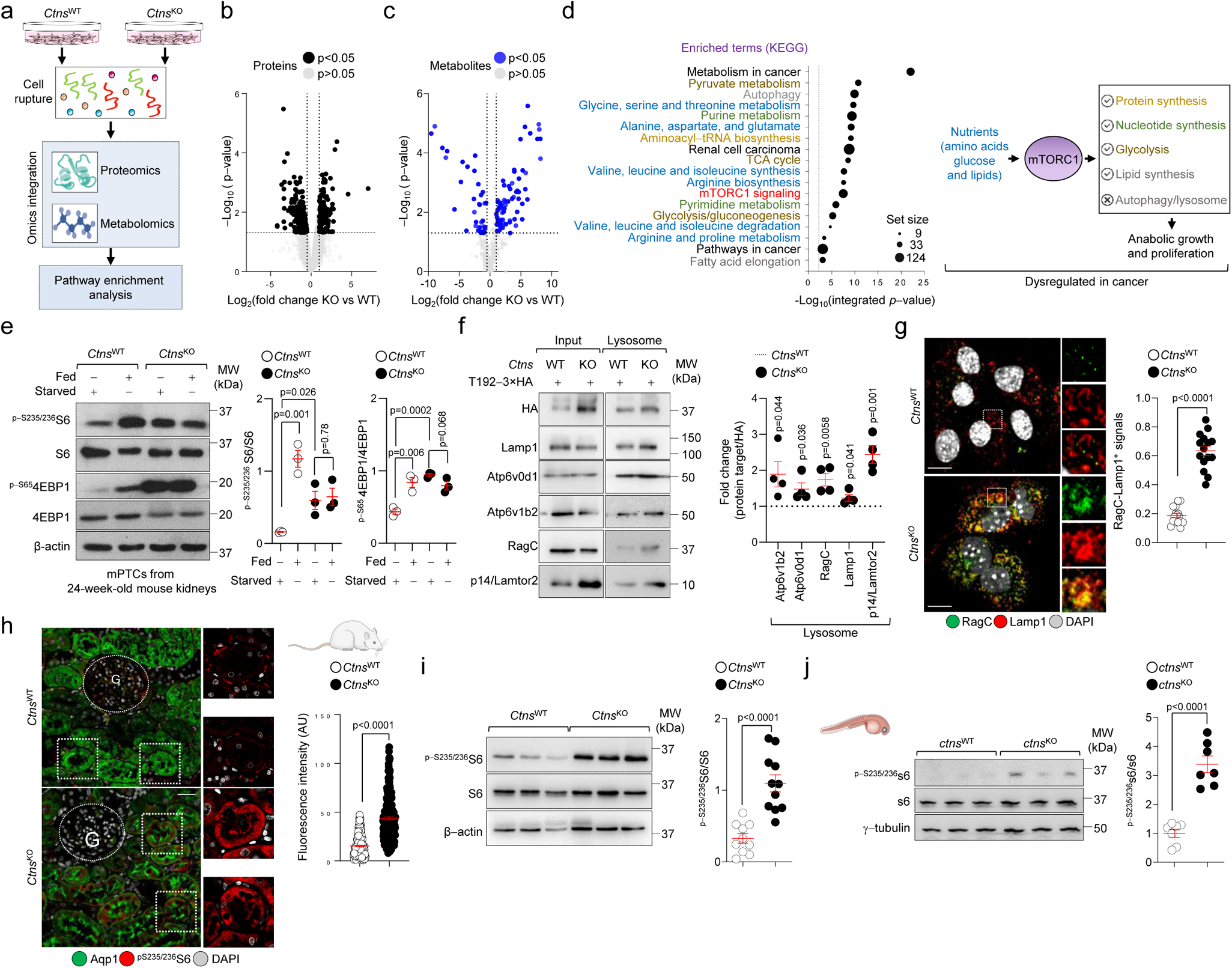
Hyperactive mTORC1 drives metabolic switch in CTNS-deleted/cystine accumulating PT cells. (**a**) Schematic showing the multi-omics integration. (**b**-**c**) Volcano plot of (**b**) proteome and (**c**) metabolome changes induced by CTNS loss in the PT cells. Black dots show the proteins and metabolites [Log_2_ (fold change) < −0.5 and Log_2_ (fold change) >1; and p−value < 0.05]. (**d,** left) Kyoto Encyclopaedia of Genes and Genomes (KEGG) pathway analysis; n=3 biologically independent samples. Black dashed line shows the threshold of significant enrichment. Black circles show the size of enriched proteins/metabolites. (**d,** right) Schematic showing the upstream regulators of mTORC1 and downstream regulated pathways. (**e**) Quantification of the indicated proteins; n=3 biologically independent replicates. (**f,** right) Immunoblots of the indicated proteins in input samples and corresponding Lyso-IP fractions from *Ctns*^WT^ and *Ctns*^KO^ mPTCs; n= 4 biologically independent experiments. (**g**) Confocal microscopy and quantification of RagC/Lamp1^+^ structures (expressed as percentage of total lysosomes; n=15 randomly selected fields of views per each condition). Dotted white squares contain images at high magnification. (**h**) Confocal microscopy and quantification of ^pS235/236^S6 MFI in Aqp1^+^ PTs (green) of *Ctns* rat kidneys; n>227 PTs pooled from 3 animals per each group. (**i**-**j**) Immunoblots and quantification of the indicated proteins in (**i**) 24-week-old *Ctns* rats or (**j**) 5-dpf-*ctns* zebrafish; n= 10 rats per group and n=7 samples per each group (with each sample representing a pool of 10 zebrafish larvae). Plots represent mean ± SEM. Statistics calculated by unpaired two-tailed Student’s *t* test. Nuclei counterstained with DAPI (grey). Scale bars are 10 μm in **g** and 50μm in **h**. Unprocessed scans of original blots shown in Supplementary Fig. 14. Source data is provided as a Source Data file.

### Temporal mTORC1 activation by cystine storage directs kidney disease development

The possibility to monitor progressive PT dysfunction (Fig. 4a) and cystine storage (Fig. 4b) over time allowed us to test whether mTORC1 activation may drive the onset and progression of kidney disease in CTNS deficient mice (Fig. 4c). Analyses in mPTCs from *Ctns*^KO^ kidneys revealed nutrient-dependent mTORC1 activation, similar to control *Ctns*^WT^ mPTCs, at 6 weeks of age (Fig. 4d). Conversely, in mPTCs from pre-symptomatic 12-week-old *Ctns*^KO^ mice, mTORC1 activation was stronger than in control cells and largely resistant to nutrient starvation (Fig. 4e). Like wild-type cells, mTOR translocated to the surface of Lamp1-flagged lysosomes in presence of nutrients in mPTCs from 6-week-old *Ctns*^KO^ kidneys, but it readily dissociated from lysosomes following nutrient starvation (Fig. 4f), as expected in response to changes in nutrient availability. Conversely, in mPTCs from 12-week-old *Ctns*^KO^ kidneys, mTOR remained clustered on Lamp1-flagged lysosomes following nutrient deprivation, contrasting with the dissociation observed in control cells (Fig. 4g). These changes were accompanied by reduced numbers of enlarged Lamp1^+^ structures in mPTCs from kidneys of 6- to 12-week-old *Ctns*^KO^ mice (<u>Supplementary Fig. 8a</u>). Despite these metabolic and signaling alterations, the *Ctns*^KO^ mouse kidneys displayed mild tubular damage (<u>Supplementary Fig. 8b,c</u>), with no changes in kidney function (blood urea nitrogen and creatinine clearance; <u>Supplementary Fig. 8d,e</u>) and no alterations in kidney growth (as indicated by the kidney-to-body weight ratio; <u>Supplementary Fig. 8f</u>) and − even in aged animals. The effect of CTNS loss on mTORC1 activity did not result from alterations in growth factor-directed or 5’-AMP activated protein (AMPK)-mediated signaling upstream of mTORC1, which were similar between wild-type and *Ctns*^KO^ mPTCs (<u>Supplementary Fig. 8g</u>). Thus, CTNS loss/ cystine storage activates mTORC1 through mechanisms that are, at least in part, distinct from growth factor-induced signals and/or energy availability.

**Figure 4.**
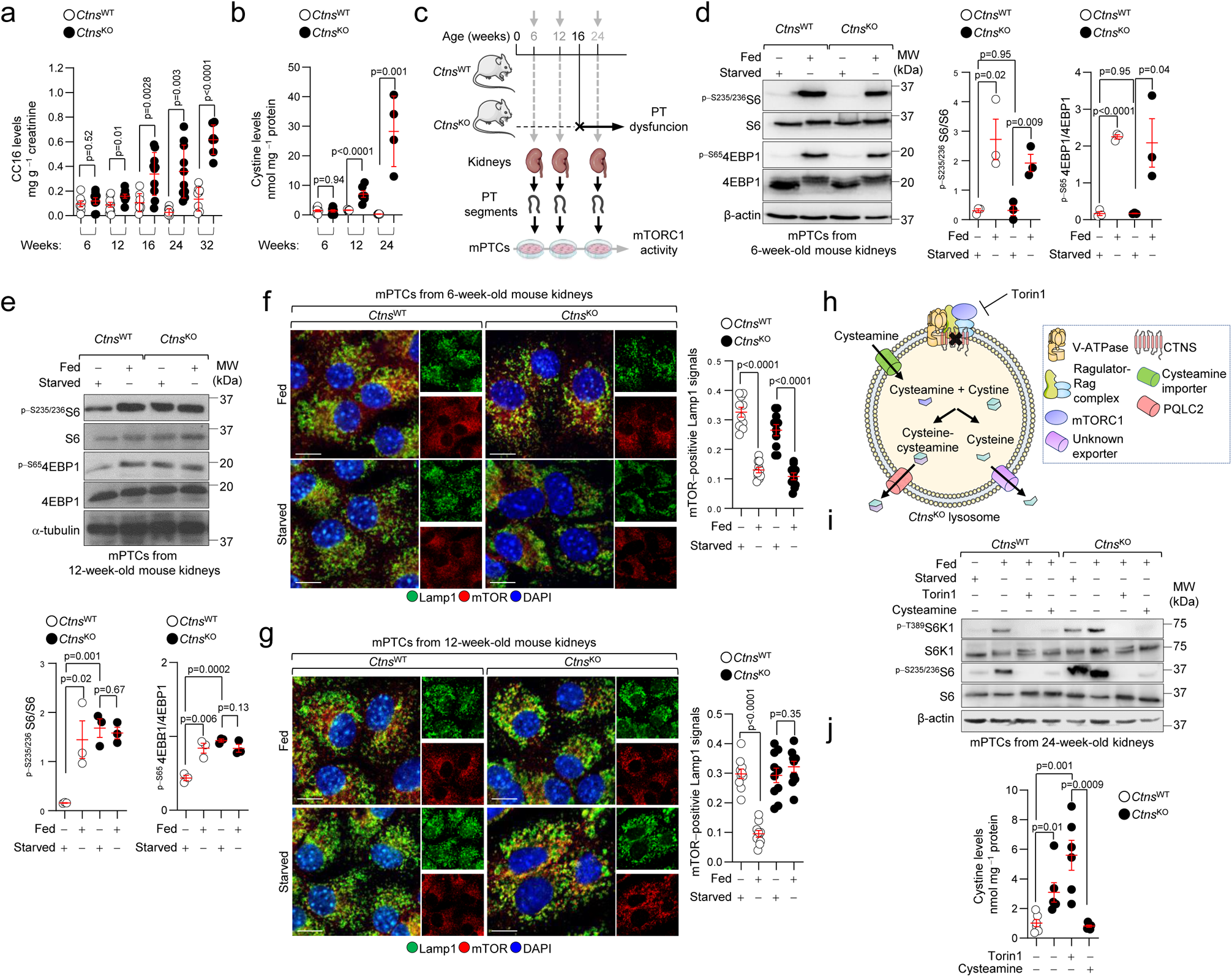
Temporal mTORC1 activation by cystine buildup leads to kidney disease development in CTNS deficient mice. (**a**) Quantification of LMW/Clara cell 16 (CC16) protein in the urine of *Ctns* mice at the indicated times; n>6 animals per each group/time point. (**b**) Cystine levels in the *Ctns* mouse kidneys; n>4 mice per each group. (**c**) Workflow for monitoring mTORC1 activity in *Ctns* mice and derived cells. mPTCs from (**d** and **f**) asymptomatic (6 weeks of age), (**e** and **g**) pre-symptomatic (12 weeks of age), and (**i**) symptomatic (24 weeks of age) *Ctns*^KO^ mice and their (age-matched) *Ctns*^WT^ littermates were cultured under fed and starved conditions. (**d** and **e**) Immunoblotting and quantification of the indicated proteins; n=3 mice per each group. (**f** and **g**) Confocal microscopy and quantification of mTOR^+^-Lamp1^+^ signals; n= 10 randomly selected and non-overlapping fields of views per each condition. Nuclei counterstained with DAPI (blue). (**h**) Fed mPTCs were treated with cysteamine (100 nM) or Torin1 (250 nM) for 16h. As illustrated in the cartoon, cysteamine penetrates lysosomes by a distinct importer and reacts with cystine to exchange its disulfide bridge into a mixed (cysteamine-cysteine) disulfide, which exits via the cationic amino-acid exporter PQLC2 (5), and generates free cysteine, which exits from lysosomes through an unknown transporter. (**i**) Immunoblotting of the indicated proteins; n=2 biologically independent experiments. (**j**) Cystine levels in *Ctns* mPTCs; n=6 biologically independent samples per each condition. Plots represent mean ± SEM. Statistics calculated by unpaired two-tailed Student’s *t* test. Scale bars, 10μm. Unprocessed scans of original blots shown in Supplementary Fig. 14. Source data are provided as a Source Data file.

### Lysosomal cystine regulates mTORC1 through Ragulator-Rag GTPase complex

We next tested whether cystine storage per se might force mTORC1 to remain on the lysosomal surface, using pharmacologic and genetic tools. Emptying the cystine stores with the drug cysteamine (29) blunted mTORC1 activation in mPTCs derived from 24-week-old *Ctns*^KO^ mice (Fig. 4h,i). Notably, the latter treatment corrected the mTORC1 activation to the same extent than the selective mTOR inhibitor Torin1 which, contrary to cysteamine, does not empty the cystine storage (Fig. 4h,i).

To substantiate the causal link between CTNS loss, lysosomal cystine storage, and hyperactive mTORC1 signaling, we transduced *Ctns*^KO^ PT cells with an adenovirus expressing HA-tagged wild-type (HA-CTNS^WT^) or Gly339Arg mutant (HA−CTNS^G339R^) CTNS (Fig. 5a; <u>Supplementary Fig. 9a</u>). This disease-causing CTNS^Gly339Arg^ mutant (30) preserves the delivery of mutant CTNS to the lysosomal surface while abrogating its transport function, thus causing cystine storage (Fig. 5b; <u>Supplementary Fig. 9b</u>). The re-introduction of CTNS^WT^ in *Ctns*^KO^ mPTCs rescued the cystine levels (Fig. 5b) and restored the nutrient-directed regulation of mTORC1 compared to *Ctns*^KO^ mPTCs transduced with an empty vector (Fig. 5c; +HA-CTNS^WT^). Conversely, the nutrient-dependent regulation of mTORC1 signaling activity was markedly abolished in *Ctns*^KO^ mPTCs expressing mutant CTNS^G339R^/ accumulating cystine (Fig. 5b), as evidenced by elevated phosphorylation rate of mTORC1 substrate S6 (Fig. 5c; +HA-CTNS^G339R^) in nutrient-depleted conditions. Immunoblotting analyses of anti-HA immunoprecipitates from *Ctns*^KO^ mPTCs transiently expressing diverse tagged CTNS forms revealed a decreased binding of CTNS^WT^ to endogenous members of Ragulator-Rag GTPases lysosomal scaffold (Lamtor2 and RagC GTPase) compared to mPTCs transducing with mutant CTNS^G339R^ (Fig. 5d). In line, the reconstitution with CTNS^WT^, but not the mutant CTNS^G339R^, rescued autophagy-lysosome degradation systems in *Ctns*^KO^ mPTCs (<u>Supplementary Fig. 9c,d</u>).

**Figure 5.**
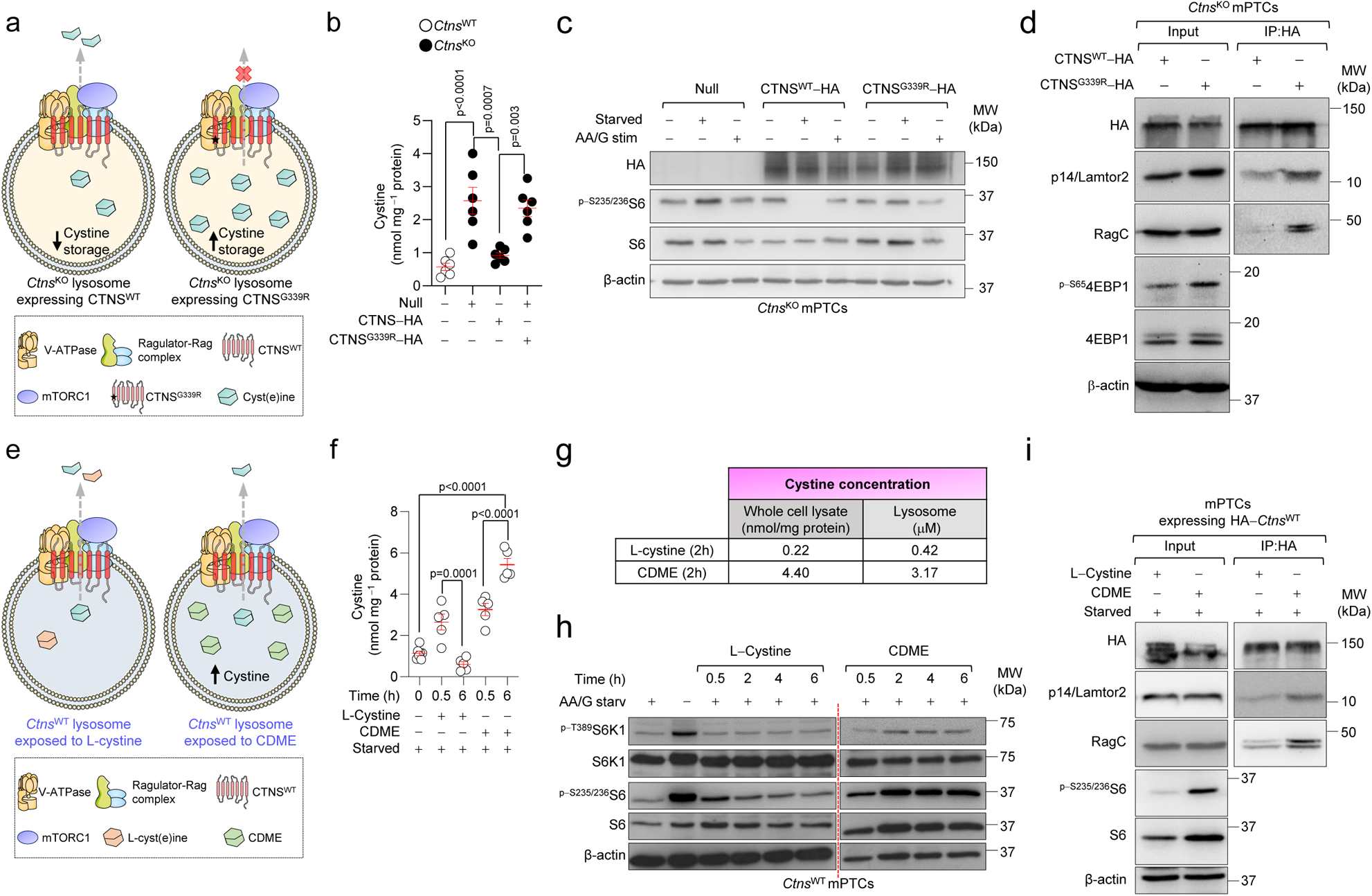
Lysosomal cystine regulates mTORC1 activity through Ragulator-Rag GTPase lysosomal scaffold. (**a**-**d**) mPTCs from *Ctns*^KO^ mice were transduced with (**a**) Null or hemagglutinin-tagged CTNS^WT^ (CTNS^WT^-HA) or HA-tagged mutant CTNS (CTNS^G339R^-HA) bearing adenoviral particles for 2 days. (**b**) Cystine levels in mPTCs; n=6 biologically independent experiments. (**c** and **h**) Cells were cultured under fed or starved conditions or restimulated by adding back amino acids and glucose (AA/G stim). (**c**) Immunoblots of the indicated proteins. (**d**) Fed *Ctns*^KO^ mPTCs transiently expressing tagged CTNS^WT^ or CTNS^G339R^ were lysed, and the samples were subjected to HA immunoprecipitation and immunoblotting for the indicated proteins. (**e**-**i**) Starved *Ctns*^WT^ mPTCs were exposed to (**e**) CDME (0.1 mM) or with unmodified L-cystine (0.1 mM) for the indicated times. (**f**) Cystine levels in *Ctns* mPTCs; n>4 biologically independent samples per each condition. (**g**) Cystine levels in whole cell lysates and purified lysosomes; n=2 biologically independent experiments. (**h**) Immunoblots of the indicated proteins; n=3 biologically independent experiments. Quantification of the indicated proteins is shown in Supplementary Fig. 9g. (**i**) The cells were lysed, and the samples were subjected to HA immunoprecipitation and immunoblotting for the indicated proteins; n=2 biologically independent experiments in **c**, **d** and **i**. Plots represent mean ± SEM. Statistics calculated by one-way ANOVA followed by Tukey’s or Sidak’s multiple comparisons test in **b** and **f**. Unprocessed scans of original blots shown in Supplementary Fig. 14. Source data are provided as a Source Data file.

To prove that lysosomal cystine is involved in this signaling loop, we bypassed the transport function of CTNS by culturing *Ctns*^WT^ mPTCs with cystine methyl ester derivates (CDME; 28,31). These amino acid derivates rapidly enter the cells and are hydrolysed by esterases into native amino acids, ultimately accumulating within mPTCs and lysosomes (Fig. 5e-g). Exposure of nutrient-depleted *Ctns*^WT^ mPTCs with low doses of CDME stimulated mTORC1 activity in a time-dependent fashion (Fig. 5h; <u>Supplementary Fig. 9e-g</u>). This lag correlated with the time required for CDME to accumulate within lysosomes and could not be recapitulated by treating starved cells with unmodified L-cystine analogues that do not accumulate (Fig. 5f,g). Immunoprecipitation followed by immunoblotting analysis revealed that the heightened levels of lysosomal cystine increase the binding of HA-tagged CTNS to endogenous components of Ragulator-Rag GTPases scaffold complex (Lamtor2 and RagC) in CDME-treated cells compared with those exposed to unmodified L-cystine analogues (Fig. 5i). Collectively, these data reveal that, through interactions with Ragulator-Rag GTPase lysosomal scaffold, CTNS regulates the activation of mTOR in response to variations in the levels of lysosomal cystine.

### Blocking mTORC1 rescues lysosome proteolysis and cell differentiation downstream of cystine storage

We next tested whether mTORC1 inhibition restores lysosomal catabolic activities and differentiating state of PT cells lacking CTNS/accumulating cystine. We treated *Ctns*^WT^ and *Ctns*^KO^ mPTCs with low, non-toxic doses of Torin1 (250nM for 16 h; <u>Supplementary Fig. 10a</u>). Treatment with Torin1, which fails to correct cystine storage (Fig. 4j) while effectively blocking mTORC1 activity (Fig. 6a, <u>Supplementary Fig. 10b</u>), enhanced the nuclear translocation of the transcription factor E3 (Tfe3), the master modulator of lysosome biogenesis and autophagy, in *Ctns*^KO^ mPTCs compared to vehicle (<u>Supplementary Fig. 10c</u>). In turn, the increased nuclear import heightened Tfe3-targeted autophagy and lysosomal genes, re-activating lysosome function and catabolic autophagy (Fig. 6b,c; <u>Supplementary Fig. 10d-f</u>). In agreement, treatment with Torin1 ameliorated the mitochondrial bioenergetics and lowered the overproduction of mitochondria-derived reactive oxygen species (ROS), preventing the disruption of tight junctions manifested in *Ctns*^KO^ mPTCs (Fig. 6d,e; <u>Supplementary Fig. 10g,h</u>). This markedly repressed the nuclear translocation of Ybx3 and its driven activation of growth/proliferation (Fig. 6f-h; <u>Supplementary Fig. 10i</u>), while restoring PT cell differentiation markers and the endocytic/reabsorptive activities of the tubular cells downstream of cystine storage (Fig. 6f,i,j,k). Similar functional rescues, including lysosomal catabolic activities, autophagy, proliferation, and differentiation were also observed in *Ctns*^KO^ cells treated with low concentrations of the allosteric mTOR inhibitor rapamycin (250nM for 16 h; <u>Supplementary Fig. 11</u>) or depleted of *Raptor* (<u>Supplementary Fig. 12</u>) − a gene encoding protein that docks the mTOR protein kinase on the surface of lysosomes (6). The modulation of mTORC1 signaling by either pharmacological or genetic means did not alter the tubular cell differentiation nor the Lrp2-mediated endocytosis in *Ctns*^WT^ mPTCs (Fig. 6f,I,j,k; <u>Supplementary Fig. 11h</u>; <u>Supplementary Fig. 12f</u>).

**Figure 6.**
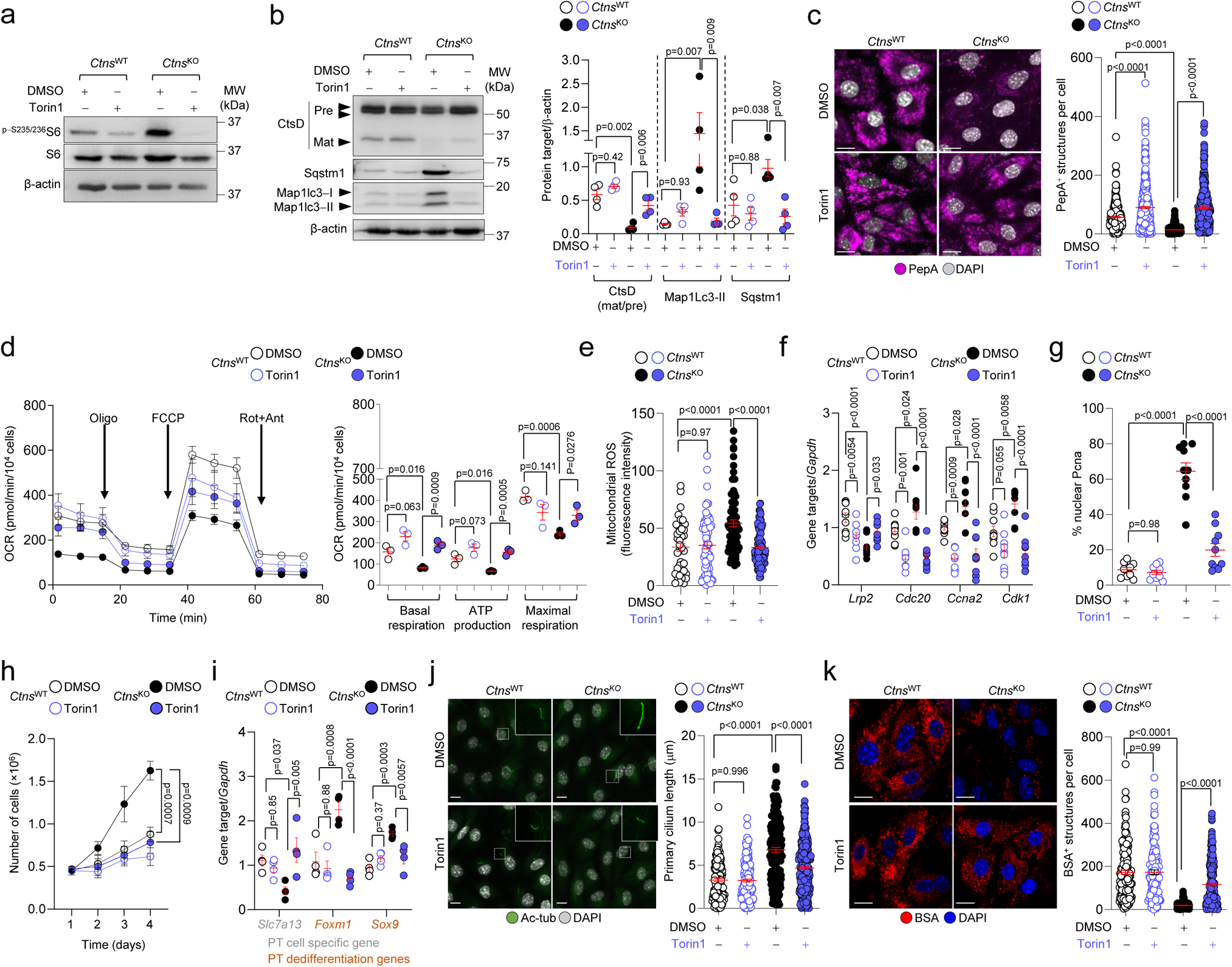
Suppressing mTORC1 rescues lysosome proteolysis and differentiation downstream of CTNS loss and cystine storage. (**a**-**j**) Cells were treated with vehicle (DMSO) or Torin1 (250nM for 16h). (**a**-**b**) Immunoblots and quantification of the indicated proteins; n= 3 (**a**) and 4 (**b**) biologically independent experiments. Quantification of the indicated proteins in (**a**) shown in Supplementary Fig. 10b. (**c**) Confocal microscopy and quantification of PepA^+^ puncta per cell; n>247 cells per each condition pooled from three biologically independent experiments. (**d**) Oxygen consumption rates (OCRs) were measured under basal level and after the sequential addition of oligomycin (Oligo, 1 μM), FCCP (0.5 μM), and Rotenone (ROT; 1μM) + Antimycin A (ANT; 1μM); n = 3 biologically independent experiments. (**e**) Confocal microscopy and quantification of MitoSOX fluorescence intensity per cell; n>42 cells per each condition. Representative images are shown in Supplementary Fig. 10g. (**f** and **i**) mRNA levels for the indicated genes; n>6 replicates (**f**) or n=4 replicates (**i**), both pooled from 3 biologically independent experiments. (**g**) Quantification of the number of Pcna^+^ nuclei; n=10 non-overlapping fields of views per each condition. Representative images are shown in Supplementary Fig. 10j. (**h**) Growth curves in *Ctns* mPTCs; n=3 biologically independent experiments. (**j**) Maximum intensity projection of image stacks and quantification of the primary cilium length (Ac-tubulin, green) in *Ctns* mPTCs; n>132 cells pooled from 3 biologically independent experiments. (**k**) Confocal microscopy and quantification of BSA^+^ structures per cell (n>175 cells per each condition pooled from 3 biologically independent experiments). Plots represent mean ± SEM. Statistics calculated by one-way ANOVA followed by Tukey’s or Sidak’s multiple comparisons test in **b**, **c**, **e**, **f**, **I**, **j**, and k; and by unpaired two-tailed Student’s *t* test in **d** and **h**. Nuclei counterstained with DAPI (blue or grey). Scale bars, 10μm. Unprocessed scans of original blots shown in Supplementary Fig. 14. Source data are provided as a Source Data file.

In a similar vein, the treatment of *Ctns* rats with low doses (32) of Rapamycin (1.5 mg/kg/day for 2 weeks; Fig. 7a) reduced/normalized mTORC1 activity and lysosomal abnormalities (Fig. 7b; <u>Supplementary Fig. 13a,b</u>), inhibited growth/proliferation (Fig. 7c) while restoring the expression of tubular differentiation markers and cuboidal tubule cell morphology (Fig. 7d-f; <u>Supplementary Fig. 13c</u>), and the Lrp2-mediated endocytosis in the PT segments of *Ctns*^KO^ rat kidneys (Fig. 7g) compared with those treating with vehicle. These functional rescues occurred in the absence of significant changes in rat kidney cystine levels (<u>Supplementary Fig. 13d</u>). Thus, genetic and pharmacological suppression of mTORC1 signaling corrects the catabolic activities of lysosomes and the differentiating states of PT cells downstream of CTNS loss/ cystine storage.

**Figure 7.**
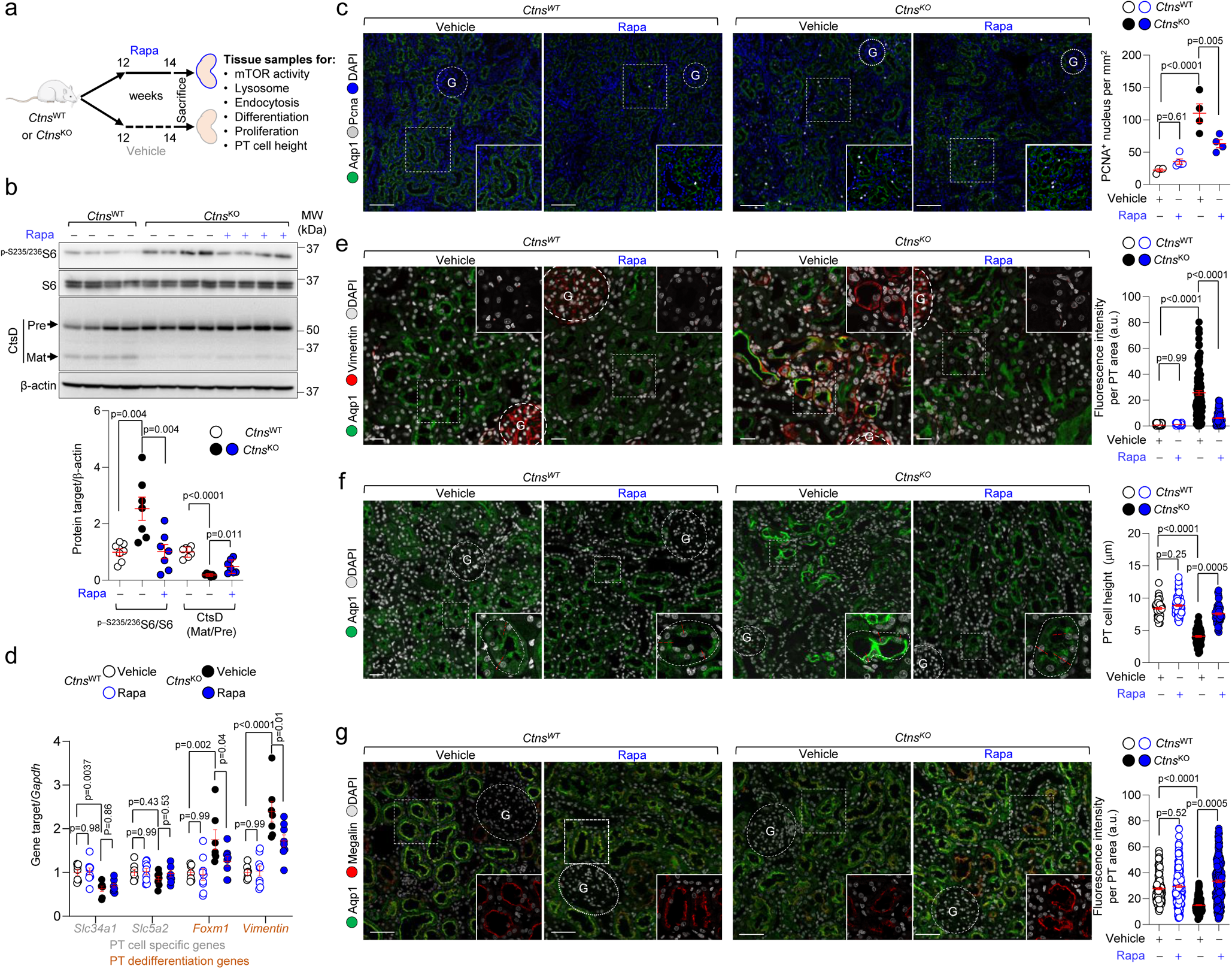
Rapamycin rescues lysosome and differentiation defects in the kidney PT of *Ctns*-deficient rats. (**a**) *Ctns* rats at 12 weeks of age were subcutaneously implanted with a pellet providing a long-term release of rapamycin (Rapa, 1.5 mg/kg/day). After 2 weeks of treatment, the kidneys were harvested and analysed. (**b**) Immunoblotting and quantification of the indicated proteins; n=7 rats per each condition. (**c**) Confocal microscopy and quantification of numbers of Pcna^+^ nuclei (white) in Aqp1-positive PT segments of *Ctns* rat kidneys; n=4 animals per each condition. (**d**) mRNA levels for the indicated genes; n>6 rats per condition. (**e**) Confocal microscopy and quantification of Vimentin (red) fluorescence intensity in Aqp1-positive PT segments of *Ctns* rat kidneys (green); n≥90 PTs pooled from 3 rats per each condition. (**f**) The whole-kidney sections stained for PT apical marker (Aqp1, green) and quantification of PT cell height in *Ctns* kidneys; n≥60 PTs pooled from 3 rats per each condition. (**g**) Confocal microscopy and quantification of Lrp2 (red) MFI in Aqp1 (green)-positive PT segments of *Ctns* rat kidneys; n≥164 PTs pooled from 4 rats per each condition. Plots represent mean ± SEM. Statistics calculate by one‒way ANOVA followed by Tukey’s or Sidak’s multiple comparisons test in **b**, **c, d**, **e**, **f**, and **g.** Scale bars are 50μm in **c** and 20μm in **e**, **f** and **g**. Unprocessed scans of original blots shown in Supplementary Fig. 14. Source data are provided as a Source Data file.

### mTORC1 inhibition improves PT dysfunction in a zebrafish model of cystinosis

Finally, to test whether mTORC1 inhibition rescues the PT dysfunction/LMW proteinuria in vivo, we treated *ctns* zebrafish larvae expressing a bona fide LMW biosensor for PT function (*lfabp10a::½* vdbp-mCherry; Fig. 8a; Ref. 33) with low, non-toxic doses of rapamycin (200 nM for 9 days). The short-term treatment with Rapamycin, which efficiently blunted mTORC1 activity (Fig. 8b), restored the lysosome-directed processing of the ultrafiltered LMW proteins (as testified by decreases in mCherry fluorescence intensity and vesicle size; Fig. 8c) and improved the PT function in the *ctns*^KO^ zebrafish pronephros, as judged by substantial decreases in LMWP/mCherry levels in the urine of treated *ctns*^KO^ zebrafish compared to those receiving the vehicle (Fig. 8d). The phenotype rescues occurred without changes in cystine levels in *ctns*-deficient zebrafish (<u>Supplementary Fig. 13e</u>). Taken together, these proof-of-concept studies indicate that modulation of mTORC1 signaling may serve as an attractive therapeutic strategy for treating PT dysfunction in cystinosis.

**Figure 8.**
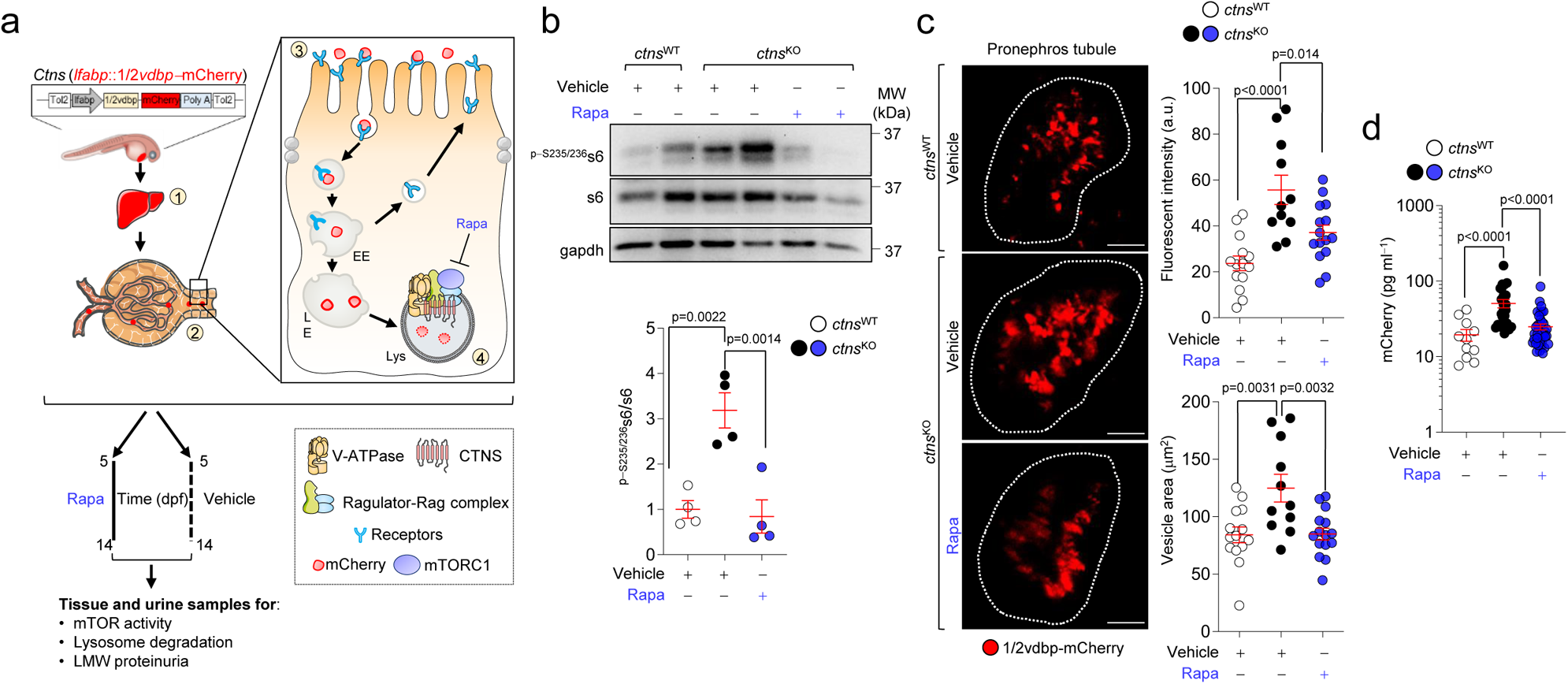
mTORC1 inhibition rescues PT dysfunction in a zebrafish model of cystinosis. (**a**) Diagram showing the generation of transgenic zebrafish line expressing a bona fide biosensor for LMW proteinuria. Fate of ½vdbp-mCherry under physiological conditions: ½vdbp-mCherry (~50 kDa) is (1) produced in liver, (2) secreted into the bloodstream and filtered by the glomerulus, (3) reabsorbed via receptor-mediated endocytosis by proximal tubule (PT) cells, and (4) processed by lysosomes. EE, early endosome; LE, late endosome; Lys, lysosome. (**b**-**d**) 5dpf-*ctns* zebrafish larvae stably expressing ½vdbp-mCherry were treated with vehicle or rapamycin (200 nM for 9 days). (**b**) Immunoblotting and quantification of the indicated proteins, with each lane representing a pool of 6 zebrafish; n=4 biologically independent experiments (**c**) Multiphoton microscopy and quantification of ½vdbp-mCherry fluorescence intensity and vesicles area per zebrafish pronephros. (**d**) After 9 days of treatment, urine samples were harvested and analysed. Quantification of urinary mCherry levels by ELISA; n= 11 vehicle-treated *ctns*^WT^ zebrafish; n=26 vehicle-treated *ctns*^KO^ zebrafish; and n=39 Rapa-treated *ctns*^KO^ zebrafish. Plots represent mean ± SEM. Statistics calculate by one‒way ANOVA followed by Sidak’s and Tukey’s multiple comparisons test, respectively, in **b** and in **c** and **d**. Scale bars, 10μm. Unprocessed scans of original blots shown in Supplementary Fig. 14. Source data are provided as a Source Data file.

## Discussion

Here, we demonstrate that cystine mobilization from lysosomes is crucial for maintaining the differentiation and endocytic function of PT cells. Lysosomal cystine storage, typically caused by the functional loss of CTNS, diverts the differentiating trajectories of PT cells towards growth/proliferation, disrupting their homeostasis and reabsorptive properties - with relevance across the spectrum of health and disease. Mechanistically, cystine storage stimulates Ragulator-Rag GTPase-dependent recruitment of mTORC1 and its constitutive activation. In cells and preclinical models of CTNS loss, the modulation of mTORC1 pathway restores lysosome function and catabolic autophagy, ultimately steering the proliferative/growing states of the cells towards differentiation and physiological homeostasis (Fig. 9).

**Figure 9.**
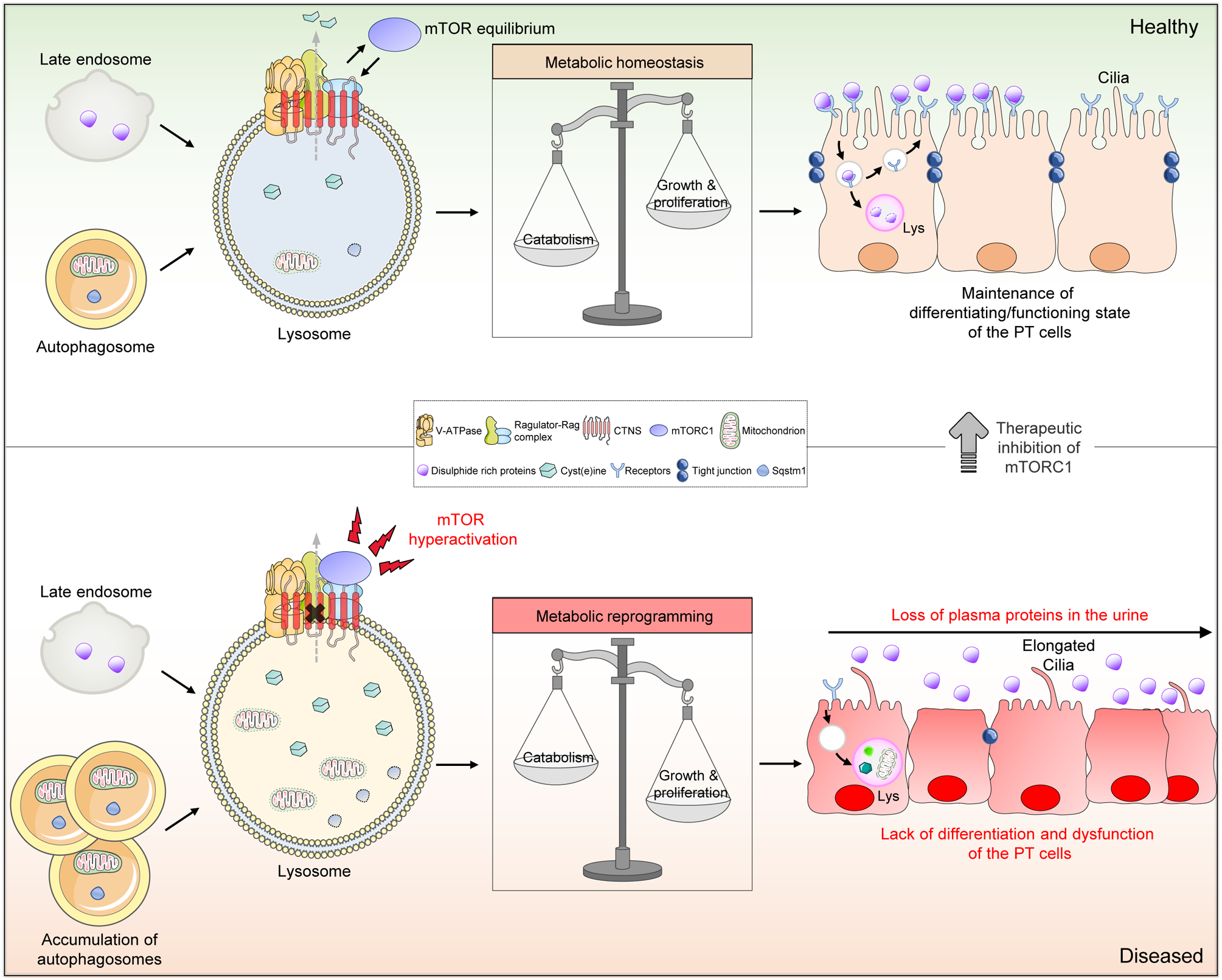
Role of CTNS-cystine-mTORC1 axis as a signaling node that instructs fate specification in the epithelial cells of the kidney tubule. The lysosomal cystine mobilization through CTNS serves as lysosomal signal that shapes the response of mTORC1 to direct metabolism and hence instruct cell fate decision during the differentiation of the kidney PT epithelium. In PT cells lacking CTNS, cystine storage triggers the Ragulator-Rag-dependent translocation of mTORC1 and its constitutive activation at the surface of the lysosome. This diverts the catabolic and differentiation trajectories of PT cells towards growth and proliferation, ultimately disrupting their homeostasis and functions. Therapeutic inhibition of hyperactive mTORC1 pathway rescues the lysosome proteolysis and rewires the proliferating trajectories of CTNS deficient PT cells towards differentiation and physiological homeostasis.

Our work indicates that lysosomes accumulating cystine due to CTNS loss display profound defects in their catabolic properties, with marked accumulation of intracellular constituents that are normally dismantled through autophagy (34). The proteolytic impairment of CTNS-deficient lysosomes occurs through mechanisms that are independent of lysosomal acidification. In lysosomes, the maintenance of cysteine levels appears to promote the catabolic activities of the lysosomal proteases (35). Thus, an abnormal storage of cystine (the oxidized dimer of cysteine) may fuel the oxidation of lysosomal thiols that preserve the folding of the catalytic sites of lysosomal proteases, impairing their degradative functions and organelle homeostasis. These catabolic anomalies are paralleled by the upregulation of programs for growth/proliferation while suppressing the differentiating states of the PT cells, hence causing defective receptor-mediated endocytosis and tubular proteinuria, i.e., the earliest manifestation of the epithelial cell disease (5). Thus, beyond cystine transport, CTNS might act as a sensor of the lysosome function − a gatekeeper of metabolism and homeostasis in the epithelial cells lining the kidney tubule.

How do CTNS-deficient/cystine accumulating lysosomes affect cell fate specialization and differentiation programs? Our integrative multi-omics analyses and AI/ML-target discovery framework, combined with cross-species validation, indicate that lysosomal cystine storage constitutively activates mTORC1 signaling in CTNS-deficient PT cells. These findings are in line with previous studies showing hyperactive mTORC1 pathway in other lysosomal storage diseases characterized by the abnormal accumulation of intracellular metabolites, e.g., cholesterol in Niemann Pick type C (NPC) and glycosaminoglycans in Mucopolysaccharidosis type VII (9,36–38). Conversely, mTORC1 activity seems to be reduced/unchanged in transformed, proliferative tubular cells derived from cystinosis patients and in patient-derived iPSCs (12,39,40). Such discrepancies may be related to immortalization protocols, which reduce per se the phosphorylation rates of canonical mTORC1 substrates (41), or to use of dedifferentiated cellular systems improper to recapitulate the characteristics of native kidney tubular cells.

The catabolic activities of the lysosome and the levels of mTORC1 activity have a crucial role in the maintenance of autophagy. Our time-course studies in *Ctns* mice suggest that defects in lysosome and abnormal (mTORC1-driven) nutrient signaling pathways, which appear early in the course of the disease, may disrupt the autophagy-mediated turnover of dysfunctional and/or damaged mitochondria, in turn destroying metabolism and inactivating the actions of fate-determining transcriptional programs that sustain differentiation and homeostasis (42). Because CTNS loss and the resulting cystine storage induce concomitant changes in the expression of genes associated with PT cell function and differentiation, as well as in the primary cilium, additional layers of (dys)regulation are likely involved.

In addition to the regulation of glutathione synthesis (43), exporting lysosomal cystine through CTNS may antagonize the reactivation of mTORC1 through the incorporation of carbons into the tricarboxylic acid cycle (TCA) cycle and limiting amino acid synthesis (13). In line, our integrated multi-omics analyses have revealed an enrichment of intermediates of TCA in the tubular cells of *Ctns*^KO^ mouse kidneys, with accumulation of dysfunctional and/or damaged mitochondria and excessive oxidative stress. The tight integration of lysosome and mitochondrial quality control systems suggests that, in diseases driven by lysosome dysfunction, their aberrant crosstalk may synergize to disrupt (mTORC1-driven) regulation of metabolic signaling and the functions of specialized cell types.

Despite the metabolic changes and signaling alterations, the *Ctns*^KO^ mice present a late onset and high variability in the extent of tubular dysfunction, mild structural changes, and no kidney failure (4,15,16), in contrast with the *Ctns*^KO^ rat model that closely recapitulates the human situation in terms of timing, severity, and multisystem involvement (23). The molecular basis of this inter-species difference may involve distinct levels of cystine storage in the lysosomes, reflecting specific transport or metabolic activities.

The mTORC1 hyperactivation evidenced in CTNS-deficient mPTCs and in the three model organisms, together with time-course analysis, suggests that CTNS tailors the activation of mTORC1 signaling in response to lysosomal cystine levels. Treatment of CTNS-defective mPTCs with cysteamine, which facilitates the export of cystine from lysosomes through the cationic amino acid PQ-loop repeat-containing protein 2 (PQLC2; ref. 5), or the exposure of wild-type mPTCs to CDME, which accumulates within lysosomes independently of CTNS, reduces and stimulates, respectively, mTORC1 activation at the lysosomal surface. Re-introduction of CTNS^WT^, which reinstates the lysosomal cystine efflux compared to disease-causing mutant CTNS^G339R^, successfully restored the sensitivity of mTORC1 to changes in nutrient availability in CTNS-defective PT cells. Conversely, CDME-induced cystine storage within lysosomes of wild-type PT cells, rendered mTORC1 insensitive to nutrient withdrawal. Our co-immunoprecipitation studies in *Ctns*^WT^ cells exposed to CDME or in *Ctns*^KO^ cells expressing CTNS^G339^ mutant suggest that, in both contexts, increased levels of cystine enable the interaction of CTNS with components of V-ATPase and Ragulator-Rag GTPase scaffold complex that recruits mTORC1 at the surface of the lysosomes. This suggests the possibility of conformational changes (44) that allows CTNS to signal cystine sufficiency to mTORC1 via the Ragulator-Rag GTPase complex. Thus, CTNS may function as a transceptor (e.g., transporter with a receptor-like function) involved not only in the mere transport of cystine but also in which the amino acid engagement is used for allosteric signal transduction, as recently described for other nutrient transporter systems (45), including the human solute transporter SLC38A9 (46, 47).

Our study offers therapeutic targets to rewire homeostasis in cystinosis, a life-threatening disease for which cysteamine has remained the standard of care despite major side-effects (5, 29). The inhibition of hyperactive mTORC1 signaling and disturbed homeostasis in diseased *Ctns*^KO^ kidneys may enhance Tfe3-mediated re-activation of cellular programs boosting lysosomes and catabolic autophagy, removing damaged (ROS-overproducing) mitochondria, stabilizing tight junctions, and repressing the Ybx3 signaling cascade that promotes dedifferentiation and growth/proliferation. However, the translatability of mTORC1 inhibitors is limited by poor availability, lack of specificity, and toxicity (6). The rescue of cellular and functional defects observed with the use of rapamycin (48) suggests therapeutic value for modulators of mTORC1 signaling in cystinosis. Given that dietary restriction of nutrients (e.g., low protein diet and/or reductions of essential and/or branched amino acids, including cystine) promote homeostasis by switching mTORC1 signaling off (49), combinations of nutritional interventions with existing and/or emerging drugs might have translatable potential in cystinosis and other currently intractable diseases.

In summary, our work describes a fundamental role of CTNS-cystine-mTORC1 signaling node as a guardian of cellular homeostasis and identifies an evolutionarily conserved lysosome-based signal that instructs fate specification in the epithelial cells of the kidney tubule. Modulating this pathway may yield therapeutic strategies for cystinosis and other lysosomal disorders, while creating opportunities to regulate homeostasis in specialized cell types.

## METHODS

### Antibodies and reagents

The following antibodies were used in this study: anti-LAMP1 (Santa Cruz Biotechnology, sc-19992, 1:1000); anti-LC3 (PM036, MBL, 1:200); anti-p62/SQSTM1 (PM045, MBL, 1:200); anti-phospho-S6 Ribosomal Protein (Ser235/236; Cell signaling technology, 4858, 1:500), anti-cathepsin D (Santa Cruz Biotechnology, sc-6486, 1:500), anti-S6 Ribosomal Protein (Cell Signaling Technology, 2217, 1:500), anti-phospho‒4E-BP1 (Ser65; Cell signaling technology, 9451, 1:500), anti-4E-BP1 (Cell Signaling Technology, 9644, 1:500), anti-phospho-p70 S6 kinase 1 (Thr389; Cell Signaling Technology, 9234, 1:500); anti-p70 S6 kinase 1 (Cell Signaling Technology, 2708, 1:500), anti-acetyl‒CoA carboxylase (Cell signaling, 3662, 1:500); anti-phospho‒acetyl‒CoA carboxylase (Ser79; Cell signaling, 3661, 1:500); anti-AMPKα (Cell signaling, 2793, 1:500); anti-phospho‒AMPKα (Thr172; Cell signaling, 2535, 1:500), anti-TFE3 (Sigma, HPA023881, 1:500); anti-mTOR (Cell Signaling Technology, 7C10, 1:400 for immunofluorescence studies), anti-Raptor (2280, Cell Signaling Technology, 2280, 1:400); anti−RagC (Cell Signaling Technology, 5466, 1:500); rabbit anti-phospho-AKT (Ser473; Cell Signaling Technology, 13038, 1:500), anti-AQP1(Aviva Systems Biology, OASA00210, 1:400), anti-PCNA (Dako, M0879, 1:500), anti-α-tubulin (Sigma Aldrich, T5168; 1:1000), anti-Cyclin A2 (Abcam, Ab181591; 1:400), anti Cdc20 (Santa Cruz, sc-13162, 1:400), anti-Golgin-97 (Cell Signaling Technology, 13192; 1:400); anti-VDAC (Cell Signaling Technology; 4866, 1:400), anti-PMP70 (Sigma, SAB4200181, 1:500), anti-calreticulin (Cell Signaling Technology, 12238, 1:400), anti-β-actin (Sigma-Aldrich, A5441, 1:1000), anti-γ-tubulin (Sigma Aldrich, clone GTU-88,1:10000), and anti-HA (Roche, 11867423001; 1:500), anti-acetylated-tubulin (Sigma,T7451; 1:400), Anti-Laminin gamma 1 (clone A5; Thermo Fisher Scientific, MA106100, 1:400), Atp6v0d1 (Thermo Fisher Scientific, PA5-103179, 1:200) ATP6V1B2 (D2F9R, Cell signaling,14617, 1:200), anti-vimentin (Abcam, Ab92547, 1:400), anti-Lipoclain2/NGAL (R&D systems, AF1857, 1:400), anti-cleaved caspase 3 (Cell signaling), 9662, 1:400), anti-caspase 3 (Cell signaling, 9661, 1:400). Compounds included Bafilomycin A1 (BfnA1; Enzo Life Sciences, ALX-380-030, 250 nM), ATP-competitive mTOR inhibitor Torin1 (TOCRIS Bioscience, CAS 1222998-36-8, 250 nM), allosteric mTOR inhibitor rapamycin (Sigma Aldrich, R0395-1MG, 250 nM in cell treatment; and MedChemExpress HY-10219-1G), Cysteamine (Sigma Aldrich, M6500-25G, 100μM), alcohol ester derivates of cystine (CDME, Sigma Aldrich 857327-5G, 0.05 mM), L-amino acid cystine (Sigma Aldrich, C7602−25G, 0.05 mM), L-cysteine (Sigma, C7352-25G, 1mM), N-Acetyl-L-cysteine (NAC, Sigma, A9165-5GNAC, 1mM), LysoTracker (Thermo Fisher Scientific, L12492), and the biotinylated Lotus Tetragonolobus Lectin (LTL; Vector Laboratories, B−1325). The pCAG R2pH-LAMP1-3xFLAG plasmid was a gift from Massimiliano Stagi (Addgene plasmid # 157940; http://n2t.net/addgene:157940; RRID: Addgene_157940). The cells used in this study were negatively tested for mycoplasma contamination using MycoAlert™ Mycoplasma Detection Kit (LT07-118, Lonza, Switzerland).

### Generation and maintenance of ctns zebrafish

The *ctns*-specific left and right TALENs (*ctns*-TALENs) were constructed in according to Golden Gate TALEN assembly protocol and using the Golden Gate TALEN and TAL Effector Kit 2.0 (Addgene, Kit #1000000024). CIscript-GoldyTALEN was a gift from Daniel Carlson & Stephen Ekker (Addgene, plasmid # 38142). TALENs were designed with the TAL Effector Nucleotide Targeter 2.0 software on the Website of Cornell University. The TALENs target the exon 3 of *ctns* zebrafish gene: left TALEN-F: TCTTTTAATCCTTTGTGTTCACA and right TALEN-R: CATCTGTAACGGTTTATTTCAAT. The spacer between two TALEN target sites is ~15 nucleotides and has an AciI restriction site in the middle, which is used for mutant screening. The TALEN expression plasmids were linearized with BamHI and then used for *in vitro* transcription (mMESSAGE mMACHINE T3 kit, Ambion). Approximately 1 nL of TALEN messenger ribonucleic acid (mRNAs, 400 ng/µL) was injected into one-cell stage zebrafish (*Danio rerio*) embryos. After 24 h, genomic DNA was extracted from injected embryos with normal appearance. Targeted genomic loci were amplified by using primers designed to anneal approximately 240 base pairs, and mutant allele was detected by AciI digestion of PCR product. The TALEN injected embryos were raised to adulthood (F0) and outcrossed with wild-type zebrafish. The embryos were then raised to adulthood (F1) for screening of heterozygous carriers. We identified a heterozygous carrier harboring *ctns^+/del8^* mutation and (F1) generations were crossed for obtaining homozygous mutant carrying *ctns^del8/del8^*. Zebrafish were kept at day/night cycle of 14/10 h at 28°C. Stable zebrafish line expressing mCherry-tagged half vitamin D binding protein (½vdbp-mCherry) in the liver was established and outcrossed with *ctns*^+/del8^ zebrafish to generate transgenic mutant line, which was crossed with *ctns*^+/del8^ zebrafish to produce homozygous larvae, and the measurement of ½vdbp-mCherry was assessed by ELISA. Where indicated, zebrafish larvae were treated at 5 dpf with zebrafish facility system water containing DMSO or Rapamycin (200 nM) for 9 days. The experimental protocols were approved by the appropriate licensing committee (Kanton Zürich Gesundheitsdirektion Veterinäramt; protocol ZH230/2018) at the University of Zurich.

### Zebrafish urine collection and ELISA mCherry measurement

Zebrafish larvae (AB backgorund) were placed in a 48-well microplate with one larva/500 μl facility water/well and kept at 28°C for 16 hours, followed by urine collection for ELISA assay (33). ½vdbp-mCherry was assayed according to manufacturer’s protocol. Briefly, 50 μL of fish pool water containing urine was distributed to a 96-well microplate pre-coated with anti-mCherry antibody. A 50 mL mixture of capture antibody and detector antibody was added to each well and incubated at room temperature for 1 hour. The wells were rinsed three times with washing buffer and incubated with 100ml of TMB development solution at room temperature for 10 minutes. The reaction was stopped by adding 100ml of stop solution, followed by reading the absorbance of each well at 450 nm.

### Rodent models

Experiments were conducted on age- and gender-matched *Ctns* knockout mouse (C57BL/6 background) (4) or rat (Sprague-Dawley background) (23) lines, and their corresponding control littermates. Rats and mice were maintained under temperature-and humidity-controlled conditions with 12 h light/12 h dark cycle with free access to appropriate standard diet in accordance with the institutional guidelines of National Institutes of Health Guide for the Care and Use of Laboratory Animals. Kidneys were collected for analyses at the time of sacrifice. The experimental protocols were approved by the appropriate licensing committee (Kanton Zürich Gesundheitsdirektion Veterinäramt; protocol ZH230/2019, ZH195/2020) at the University of Zurich. The rats aged 12 weeks were subcutaneously implanted with a slow-release pellet (Innovative Research of America, FL, USA). The pellets were designed to allow a constant release of rapamycin (1.5mg/kg B.W./day, R-5000, LC Laboratories, Woburn, MA). Control animals received a placebo pellet equivalent to their respective rapamycin-treated group. After 2 weeks of treatment, the rats were sacrificed, and kidney tissues were collected for analyses.

### Kidney function

The mice were placed overnight in metabolic cages with ad libitum access to food and drinking water; urine was collected on ice, body weight, water intake and diuresis were measured. Blood (from sublingual vein) was obtained after anesthesia with ketamine/xylazine or isoflurane. Urine and blood parameters were measured using UniCel DxC 800 pro Synchron (Beckman Coulter, Fullerton, CA, USA). Urinary levels of low-molecular weight Clara cell protein (CC16) were measured by using an enzyme-linked immunosorbent assay in according to the manufacturer’s instructions (BIOMATIK EKU03200, Thermo Fischer Scientific, Waltham, MA).

### Primary cultures of mouse proximal tubular cells

The kidneys were harvested from *Ctns* knockout mice and from their corresponding control littermates: one kidney was split transversally, and one half was fixed and processed for immunostaining while the other half was flash-frozen, homogenized by Dounce homogenizer in 1 mL of RIPA buffer that contains protease and phosphatase inhibitors and processed for western blot analysis. The contralateral kidney was taken to generate primary cultures of mPTCs (4). Freshly microdissected PT segments were seeded onto collagen-coated chamber slides (C7182, Sigma-Aldrich) and/or collagen coated 6- or 24-well plates (145380 or 142475, Thermo Fisher Scientific), and cultured at 37°C and 5% CO_2_ in DMEM/F12 (21041-025, Thermo Fisher Scientific) with 0.5% dialyzed fetal bovine serum (FBS), 15mM HEPES (H0887, Sigma-Aldrich), 0.55mM sodium pyruvate (P2256, Sigma Aldrich), 0.1ml L^−1^ non-essential amino acids (M7145, Sigma Aldrich), hydrocortisone, human EGF, epinephrine, insulin, triiodothyronine, TF, and gentamicin/amphotericin (Single Quots® kit, CC‒4127, Lonza), pH 7.40, 325mOsm kg^−1^. The medium was replaced every 48 h. Confluent monolayers of mPTCs were expanded from the tubular fragments after 6–7 days, characterized by a high endocytic uptake capacity. These cells were negatively tested for mycoplasma contamination.

### Cystine measurements

Mouse or rat kidney tissue, or zebrafish embryos or primary cultured cells were homogenized and lysed with N-ethylmaleimide (NEM) solution containing 5.2 mmol l−1N-ethylmaleide in 10mmol L^−1^potassium phosphate buffer adjusted to pH 7.4. The lysates were collected and precipitated with sulfosalicylic acid (12% w/v) and centrifuged at 10,000 rpm. for 10 min at 10 °C. The resulting supernatant was dissolved in citrate loading buffer (Biochrom Ltd, Cambridge, UK) and 50μl of this solution was analysed by Biochrom 30 Plus Amino Acid Analyzer (Biochrom Ltd). The protein pellet was dissolved in 0.1 mol L^−1^NaOH solution and the protein concentration was determined by Biuret method. The concentration of amino acids was measured by using a lithium high performance physiological column followed by post-column derivatization with ninhydrin. The amino acids were identified according to the retention time and the ratio of the area between the two wavelengths (570nm and 440 nm) and quantified by using EZChrom Elite software (Agilent Technologies Inc., Pleasanton, California, USA). Cystine concentration was normalized to the protein concentration and reported in nmol per mg protein.

### Nutrient starvation protocols and cell treatments

The withdrawal of serum, glucose, and amino acid was performed by washing mPTCs with Hank’s balanced salt solution (55021 C, Sigma-Aldrich) and placing them in normal growth and nutrient−deprived medium (amino acid-free RPMI, starvation medium) for the indicated times. For amino acid starvation and stimulation experiments, the primary cells were rinsed twice with and incubated for 2h starvation medium or incubated in starvation medium and then stimulated with acid-free RPMI supplemented for 60 minutes with a standard amino acid mixture composed of MEM nonessential amino acid solution, MEM essential amino acid solution, and L−glutamine (Invitrogen, Thermo Fisher Scientific) and with 0.5 % fetal calf serum that had been dialyzed against phosphate-buffered saline (PBS) in dialysis cassettes (Thermo Scientific) having an 3500 molecular weight cut-off. Where indicated, lysosomal proteolysis was inhibited by addition of BfnA1 (250 nM in cell culture medium for 2h and/or 4h). Where indicated, the cells were treated with either the allosteric mTORC1 inhibitor Rapamycin (250 nM in cell culture medium for 16h) or the ATP-competitive mTORC1 inhibitor Torin1 (250 nM in cell culture medium for 16h) or cysteamine (100 μM in cell culture medium for 16h). Where indicated, mPTCs were exposed to alcohol ester derivates of cystine (CDME, 0.1 mM) and L−amino acid cystine (0.1 mM) at the indicated time point. Afterwards, the cells were processed and analysed as described below.

### Adenovirus transduction

For RNA interference studies, the adenovirus constructs include scrambled short hairpin (Scmb) or shRNAs targeting mouse *Raptor*. For expression studies, adenoviral constructs used include CMV (control vector, Ad-CMV-GFP, Vector Biolabs) or an individually carrying mouse hemagglutinin (HA) tagged-*Ctns or* carrying mouse HA tagged-mutant *Ctns* (Gly339Arg; CTNS^G339R^) or 3×HA-tagged TMEM192 or 3×HA-tagged R2pH-LAMP1 or expressing mouse green fluorescent protein (GFP)-tagged-*Map1lc3b* or expressing mouse red fluorescent protein (RFP)-tagged-*Lamp1*. The adenoviral constructs were purchased from Vector Biolabs (University City Science Center, Philadelphia, USA). The cells were plated onto collagen-coated chamber slides or 24-well or 6-well tissue culture plates. 24 h after plating the cells with approximately 70-80% confluence, adenovirus transduction was performed by incubating the cells for 16 h at 37°C with culture medium containing the virus. The cells were then challenged with fresh culture medium every 2 days, cultured for 2 days for the expression studies, 5 days for RNA interference studies, and collected for analyses.

### Microarray and bioinformatics analysis

The Affymetrix (GeneChip mouse genome 430A 2.0 array) hybridization experiments were performed in triplicate at the Coriell Genotyping and Microarray Center (Coriell Institute for Medical Research, Camden, New Jersey, USA) on total RNA extracted from microdissected proximal tubules of *Ctns* knockout mouse kidneys and controls. Normalized expression levels were generated from the raw intensities using the RMA method implemented in oligo Bioconductor package (51). Differentially expressed probes between conditions (*Ctns* knockout versus controls) were identified using the Bioconductor package limma (50). *P* values were corrected for multiple testing using the Benjamin–Hochberg method. The probes were annotated using the mouse430a2.db Bioconductor Annotation package. Pathway enrichment analysis was performed using the function enrichPathway from the Bioconductor package ReactomePA, run with the default parameters, selecting “mouse” as organism, and using all the genes on the microarray as universe. The analysis was performed separately for the up-regulated and the down-regulated genes. The modules associated with mouse cystinosin gene were determined using GeneBridge toolkit (www.systems-genetics.org; ref. 20).

### Bulk RNA sequencing

Total RNA was isolated from kidney cortex samples from *Ctns* mouse (n=4 animals per each group) and rat (n=8 animals per each group) models using the Rneasy Plus Mini kit (). RNA-seq libraries were prepared using the TruSeq Stranded mRNA-seq reagents (Illumina) using 100 ng of total RNA following the protocol provided by the supplier (Genetic Diagnostic and Sequencing Services, Tubingen, Germany). The quality of the total RNA and the RNA-seq libraries was assessed on Fragment Analyzer (Agilent). The libraries were sequenced on Illumina NovaSeq6000 using the 100-nucleotide paired-end-run configuration following the protocol provided by the supplier.

### Proteomics

#### Sample preparation

Cell pellets (~5×10^5^ cells; n=3 mice per each group) were solubilized in 100μl of lysis buffer, treated with High Intensity Focused Ultrasound (HIFU) for 1 minute at an ultrasonic amplitude of 85%, and boiled at 95°C for 10 minutes. Proteins were extracted using a tissue homogenizer (Tissue Lyser II, QUIAGEN) for 2 cycles of 2 minutes at 30 Hz, boiled at 95°C for another 10 minutes, and subjected to a last cycle of 1 min of HIFU. The samples were centrifuged at 20000*g* for 10 min, and the protein concentration was determined using a Lunatic (Unchained Labs) instrument. For each sample, 50 µg of protein were taken. The samples were digested by adding 10μl of the ‘Digest’ solution. After 60 min of incubation at 37°C the digestion was stopped with 100μl of Stop solution. The solution was transferred to the cartridge and were removed by centrifugation at 3800 × g, while the peptides were retained by the iST-filter. Finally, the peptides were washed, eluted, dried and re-solubilized in 40μl of 3% acetonitrile, 0.1% FA. 1 µl of iRT peptides (Biognosys) at 1:100 dilution was added to each sample.

#### Liquid chromatography-mass spectrometry analysis

Mass spectrometry analysis was performed on an Orbitrap Fusion Lumos (Thermo Scientific) equipped with a Digital PicoView source (New Objective) and coupled to a M−Class UPLC (Waters). Solvent composition at the two channels was 0.1% formic acid for channel A and 0.1% formic acid, 99.9% acetonitrile for channel B. For each sample 1μl of peptides were loaded on a commercial MZ Symmetry C18 Trap Column (100Å, 5 µm, 180 µm × 20 mm, Waters) followed by nanoEase MZ C18 HSS T3 Column (100Å, 1.8 µm, 75 µm × 250 mm, Waters). The peptides were eluted at a flow rate of 300 nl/min by a gradient from 5 to 22% B in 80 min and 32% B in 10 min after an initial hold at 5% B for 3 min. The column was washed with 95% B for 10 min and afterwards the column was re-equilibrated to starting conditions for additional 10 min. Samples were acquired in a randomized order. The mass spectrometer was operated in data-dependent mode (DDA) acquiring a full-scan MS spectra (300−1’500 m/z) at a resolution of 120’000 at 200 m/z after accumulation to a target value of 500’000. Data-dependent MS/MS were recorded in the linear ion trap using quadrupole isolation with a window of 0.8 Da and HCD fragmentation with 35% fragmentation energy. The ion trap was operated in rapid scan mode with a target value of 10’000 and a maximum injection time of 50ms. Only precursors with intensity above 5’000 were selected for MS/MS and the maximum cycle time was set to 3 s. Charge state screening was enabled. Singly, unassigned, and charge states higher than seven were rejected. Precursor masses previously selected for MS/MS measurement were excluded from further selection for 20 s, and the exclusion window was set at 10 ppm. The samples were acquired using internal lock mass calibration on m/z 371.1012 and 445.1200. The mass spectrometry proteomics data were handled using the local laboratory information management system (LIMS).

#### Protein identification and label free protein quantification

The acquired raw MS data were processed by MaxQuant (version 1.6.2.3), followed by protein identification using the integrated Andromeda search engine. Spectra were searched against a Swissprot canonical mouse proteome (version from 2019-07-09), concatenated to its reversed decoyed fasta database and common protein contaminants. Carbamidomethylation of cysteine was set as fixed modification, while methionine oxidation and N-terminal protein acetylation were set as variable. Enzyme specificity was set to trypsin/P allowing a minimal peptide length of 7 amino acids and a maximum of two missed cleavages. MaxQuant Orbitrap default search settings were used. The maximum false discovery rate (FDR) was set to 0.01 for peptides and 0.05 for proteins. Label free quantification was enabled and a 2-minute window for match between runs was applied. In the MaxQuant experimental design template, each file is kept separate in the experimental design to obtain individual quantitative values. Protein fold changes were computed based on Intensity values reported in the proteinGroups.txt file. A set of functions implemented in the R package SRM Service was used to filter for proteins with 2 or more peptides allowing for a maximum of 3 missing values, and to normalize the data with a modified robust z-score transformation and to compute p-values using the *t*-test with pooled variance. If all measurements of a protein are missing in one of the conditions, a pseudo fold change was computed replacing the missing group average by the mean of 10% smallest protein intensities in that condition.

### Metabolomics

#### Sample preparation

Cell pellets (~5×10^5^ cells; n=3 mice per each group) were lysed and extracted with 500μL lysis solution (methanol: water 4:1, v/v) by vortexing for 30 min at 4°C. Precipitated proteins were pelleted by centrifugation (16,000*g*, for 15 min at 4 °C) and 50uL of the supernatants (50μL) were transferred to a clean Eppendorf vial. The transferred aliquot of the supernatants was kept at 35°C and dried down under a gentle flow of nitrogen. The dried extracts were reconstituted in 20uL water and 80uL injection buffer. 50uL of the reconstituted extract was transferred to a glass vial with narrowed bottom (Total Recovery Vials, Waters) for LC-MS injection. In addition, method blanks, QC standards, and pooled samples were prepared in the same way to serve as quality controls for the measurement. Injection buffer was composed of 90 parts of acetonitrile, 9 parts of methanol and 1 part of 5M ammonium acetate.

#### Liquid chromatography/mass spectrometry analysis

Metabolites were separated on a nanoAcquity UPLC (Waters) equipped with a BEH Amide capillary column (150 μm x50mm, 1.7μm particle size, Waters), applying a gradient of 5mM ammonium acetate in water (A) and 5mM ammonium acetate in acetonitrile (B) from 5% A to 50% A for 12min. The injection volume was 1μL. The flow rate was adjusted over the gradient from 3 to 2 μl/min. The UPLC was coupled to Synapt G2Si mass spectrometer (Waters) by a nanoESI source. MS1 (molecular ion) and MS2 (fragment) data was acquired using negative polarization and MS^E^ over a mass range of 50 to 1200 m/z at MS1 and MS2 resolution of >20’000.

#### Untargeted metabolomics data analysis

Metabolomics data sets were evaluated in an untargeted fashion with Progenesis QI software (Nonlinear Dynamics, Waters), which aligns the ion intensity maps based on a reference data set, followed by a peak picking on an aggregated ion intensity map. Detected ions were identified based on accurate mass, detected adduct patterns and isotope patterns by comparing with entries in the Human Metabolome Data Base (HMDB). A mass accuracy tolerance of 5mDa was set for the searches. Fragmentation patterns were considered for the identifications of metabolites. All biological samples were analysed at in triplicate and quality controls were run on pooled samples and reference compound mixtures to determine technical accuracy and stability. List of ranked proteins and metabolites had crossed each other, and the enrichment pathway analysis was performed using the OmicsNet 2.0 software.

### PandaOmics TargetID platform for target identification

In silico-based PandaOmics target discovery/scoring approach was applied to identify novel molecular targets for cystinosis. This approach is based on the combination of multiple scores derived from text and omics data (27, 51). Text-based scores are derived from various sources including scientific publications, grants, patents, clinical trials, and the key opinion leaders, and thus represent how strongly a particular target is associated with a disease. Specifically, text-based scores contain Attention, Trend, Attention Spike, Evidence, Grant funding, Funding per Publication, Grant Size, Average Hirsch, Impact Factor and Credibility attention index scores. In contrast, omics scores are based on the differential expression, GWAS studies, somatic and germline mutations, interactome topology, signaling pathway perturbation analysis algorithms, knockout/overexpression experiments and omics-data sources, and thus represent the target-diseases association according to molecular connections between proposed target and disease of interest. Omics scores include thirteen models (Heterogeneous Graph Walk, Matrix Factorization, Interactome Community, Causal inference, Overexpression/Knockout, Mutated/Disease Submodules, Mutations, Pathway, Network Neighbors, Relevance and Expression) that can be subdivided into classic bioinformatics approaches and complex AI-based models. For example, the Expression score relies on the combination of each gene’s fold change difference in disease versus control samples, the statistical significance of this change, and basal expression in the disease-relevant tissue. On the other hand, AI-based omics model called Heterogeneous Graph Walk (HeroWalk) is a guided random walk-based approach that is applied to a heterogeneous graph. The model learns node representations and then identifies gene nodes, which are close to the reference disease node. The “walks” are sampled with a predefined metapath, i.e., fixed sequence of node types in a walk, e.g., “gene”–“disease”–“gene”. The node degree controls the probability of transition between the nodes while sampling, following by the SkipGram model that learns the representation of each node based on the resulting corpus of walks. The cosine similarity between the specific disease and all genes produces a ranked list of genes. The top genes from this list are predicted to be promising target hypotheses. Combination of described scores results in a ranked list of targets proposed for a given disease and can be filtered out based on their novelty, small molecules synthesis availability, availability of PDB structure and other useful filters. Description of all mentioned scores and filters is available in the User manual section of PandaOmics (https://insilico.com/pandaomics/help). Targets that are not druggable by the small molecules and do have red flags in terms of safety were excluded from the analysis. After filtering, a ranked list of proposed targets was extracted from PandaOmics and presented as a heatmap.

### Reverse transcription-quantitative PCR

Total RNA was extracted from mouse tissues using Aurum^TM^ Total RNA Fatty and Fibrous Tissue Kit (Bio−Rad, Hercules, CA). DNAse I treatment was performed to cut genomic DNA contamination. Total RNA was extracted from cell cultures with RNAqueous^R^ kit (Applied Biosystems, Life Technologies). One μg of RNA was used to perform the reverse transcriptase reaction with iScript **^TM^** cDNA Synthesis Kit (Bio-Rad). Changes in mRNA levels of the target genes were determined by relative RT-qPCR with a CFX96^TM^ Real‒Time PCR Detection System (Bio-Rad) using iQ **^TM^** SYBR Green Supermix (Bio-Rad). The analyses were performed in duplicate with 100nM of both sense and anti-sense primers in a final volume of 20 µL using iQ^TM^ SYBR Green Supermix (Bio-Rad). Specific primers were designed using Primer3 (Supplementary Table 1<u>-2</u>). PCR conditions were 95°C for 3 min followed by 40 cycles of 15 sec at 95°C, 30 sec at 60°C. The PCR products were sequenced with the BigDye terminator kit (Perkin Elmer Applied Biosystems) using ABI3100 capillary sequencer (Perkin Elmer Applied Biosystems). The efficiency of each set of primers was determined by dilution curves. The program geNorm version 3.4 was applied to characterize the expression stability of the candidate reference genes in kidneys and six reference genes were selected to calculate the normalization factor. The relative changes in targeted genes over *Gapdh* mRNAs were calculated using the 2^−ΔΔCt^ formula.

### Endocytosis assay in mouse kidneys and derived PT cells

*Mouse kidneys.* Receptor-mediated endocytosis was monitored in the proximal tubule of mouse kidneys PTs by measuring the uptake of β-lactoglobulin (L3908, Sigma). Briefly, β-lactoglobulin was tagged with Cy5 using TM2 Ab labelling kit (Amersham) following the manufacturer’s instructions. Fifteen minutes after tail-vein injection of Cy5-β-lactoglobulin (1 mg/kg B.W) mice were anesthetized and their kidneys were harvested and processed by confocal microscopy.

#### Primary PT cells

The endocytic uptake was monitored in mPTCs cells following incubation for 60 min at 4 °C with 50μg × ml^−1^bovine serum albumin (BSA)–Alexa−Fluor−647 (A34785, Thermo Fisher Scientific) in complete HEPES-buffered Dulbecco’s modified Eagle’s medium. The cells were given an acid wash and warmed to 37 °C in growth cell medium for 15 min before being fixed and processed for immunofluorescence analyses.

### Lysosomal degradation

The detection of lysosomal activity was performed in live in kidney cells by using Bodipy‒FL‒PepstatinA (P12271, Thermo Fischer Scientific) or MagicRed −(RR)_2_ substrate (MR−CtsB; 938, Immunochemistry Technologies) according to the manufacturer’s specifications. The cells were pulsed with 1μM Bodipy‒FL‒Pepstatin A or with 1μM MagicRed −(RR)_2_ in Live Cell Imaging medium for 1h at 37°C, fixed and subsequently analyzed by confocal microscopy. The number of PepstatinA‒or MagicRed positive structures per cell were quantified by using the open-source cell image analysis software CellProfiler^TM^ as described below.

### Luminal lysosomal pH

Cells expressing an empty or 3×HA-tagged R2pH-LAMP1 (24) were treated in the presence and in the absence of non-saturating concentrations of Bafilomycin A1 as previously described (5) and analysed by confocal microscopy in a chamber heated to 37 °C at 5% CO_2_. mCherry and pHluorin fluorophores were excited at 561 nm and 488 nm, respectively and both channels acquired simultaneously to minimize misalignment between channels. Where indicated, the *Ctns* cells were also stained with LysoTracker dye (1μM for 4h). After washing, the cells were subsequently analysed by confocal microscopy in a chamber heated to 37°C at 5% CO_2_. Imaging settings were maintained with the same parameters for comparison between different experimental conditions. Images were acquired using a Leica SP8 confocal laser scanning microscope (Center for Microscopy and Image Analysis, University of Zurich) and the fluorescence intensity was quantified by the open-source image processing software Fiji (ImageJ, NIH).

### Fluorescence microscopy

Fresh mouse and rat kidneys were fixed by perfusion with 50-60 mL of 4% paraformaldehyde in PBS (158127, Sigma-Aldrich), dehydrated and embedded in paraffin at 58C. Paraffin blocks were sectioned into consecutive five μm-thick slices with a Leica RM2255 rotary microtome (Thermo-Fisher Scientific) on Superfrost Plus glass slides (Thermo-Fisher Scientific). Before staining, slides were deparaffinized in Xylenes (534056, Sigma-Aldrich) and rehydrated. Antigen retrieval was carried out by heating the slides at 95°C for 10 min in 10 mM sodium citrate buffer (pH 6.0). The slides were quenched with 50 mM NH_4_Cl, blocked with 3% BSA in PBS Ca/Mg (D1283, Sigma-Aldrich) for 30 min and stained with primary antibodies diluted in blocking buffer overnight at 4°C. After two washes in 0.1% Tween 20 (v/v in PBS), the slides were incubated with the corresponding fluorophore-conjugated Alexa secondary antibodies (Invitrogen) diluted in blocking buffer at room temperature for 1h and counterstained with 1μg Biotinylated Lotus Tetragonolobus Lectin (LTL; B-1325 Vector Laboratories) and 1µM 4’,6-Diamino-2-phenylindole dihydrochloride (DAPI; D1306, Thermo Fischer Scientific). The slides were mounted in Prolong Gold Anti‒fade reagent (P36930, Thermo Fisher Scientific) and analyzed by confocal microscopy. The images were acquired using Leica SP8 confocal laser scanning microscope (Center for Microscopy and Image Analysis, University of Zurich) equipped with a Leica APO 63x NA 1.4 oil immersion objective at a definition of 1,024 x 1,024 pixels (average of eight or sixteen scans), adjusting the pinhole diameter to 1 Airy unit for each emission channel to have all the intensity values between 1 and 254 (linear range). The micrographs were processed with Adobe Photoshop (version CS5, Adobe System Inc., San Jose, USA) software. Quantitative image analysis was performed by selecting randomly ~5-10 visual fields per each slide that included at least 3-5 PTs (LTL-positive or AQP1-positive where indicated), using the same setting parameters (pinhole, laser power, and offset gain and detector amplification below pixel saturation). For Pcna staining on rat kidney tissue following mTOR inhibition, the kidney sections were analyzed using a fluorescence microscope. The images were acquired using Leica Dmi8 wide-field fluorescence microscope equipped with a Leica HC PL APO x20/0.80 objective, and a Leica DFC9000 GTC camera. The full kidney slides were scanned and PCNA signal in the cortex of each kidney was quantified using the LASX software.

The cells were fixed for 10 min with 4% PFA in PBS, quenched with 50 mM NH_4_Cl and permeabilized for 20 min in blocking buffer solution containing 0.1% Triton X-100 and 0.5% BSA dissolved in PBS. Subsequently, cells were incubated overnight with the appropriate primary antibodies at 4°C. After repeated washing with PBS, the slides were incubated for 45 min with the suitable fluorophore-conjugated Alexa secondary antibodies (Invitrogen), counterstained with 1µM DAPI for 5 min, mounted with the Prolong Gold Anti-fade reagent and analyzed by a Leica SP8 confocal laser scanning microscope (Center for Microscopy and Image Analysis, University of Zurich) using the settings described above. Quantitative image analysis was performed by selecting randomly 5 visual fields pooled from biological triplicates, with each field including at least 10-15 cells, using the same setting parameters (e.g., pinhole, laser power, and offset gain and detector amplification below pixel saturation). The quantitative cell image analyses of subcellular structures were determined by using the open-source cell image analysis software CellProfiler^TM^ (52). In particular, the specific module “Measure-Object-Intensity-Distribution” was used to score the number of Map1Lc3b, or PepA or MR−positive structures. The pipeline “Cell/particle counting and scoring the percentage of stained objects” was used to score the fractions of Lamp1-positive structures that were also positive for Map1Lc3b. The “Cytoplasm-Nucleus Translocation Assay” was used to score the numbers of Ybx3 and Pcna-positive nuclei. For quantification of colocalization, non-overlapping images were acquired from each coverslip. Raw, unprocessed images were imported into FIJI v.2.0.0-rc-69/1.52i and converted to 8-bit images, and images of individual channels were thresholded independently to eliminate background and non-specific staining noise and converted to binary masks. Co-localization between Lamp1 and mTOR or RagC and Lamp1 was determined using the ‘‘AND’’ function of the image calculator. An open-source image processing software Fiji (ImageJ, NIH) was used for to measure the cilia length and the tubular height. The number of cells and/or fields of views used per each condition and the number of biologically independent experiments are indicated in the figure legends.

#### Zebrafish

Quantitative analysis of fluorescent signals was performed with one whole pronephric tubule for each larva using the same setting parameters. A multi-photon fluorescence microscope (Leica SP8 MP DIVE Falcon; Leica Microsystems, Heerbrugg, Switzerland) was used to acquire high-resolution images to analyse the fluorescent vesicles in the proximal tubules by using 25/1.0 NA water immersion objective (HC IRAPO L, Leica). The excitation wavelength was 1040 nm tunable laser for imaging of mCherry. The acquired data were processed by Huygens software (Scientific Volume Imaging, Hilversum, The Netherlands) for deconvolution, followed by segmentation using Ilastik software (EMBL, Heidelberg, Germany) and quantification of fluorescence intensity, vesicle number and vesicle area for each stack using an open-source image processing software Fiji (ImageJ, NIH).

### Immunoprecipitation

The cells (~5000000 cells) were lysed in a buffer containing 25mM Tris/HCl adjusted to pH 7.4, 150mM NaCl, 1% NP-40, and 1mM EDTA, 5% glycerol, protease, and phosphatase inhibitors for 10 min at 4 °C. Lysates were centrifuged at 12,000 r.p.m. for 10 min and the supernatants were incubated with 50 µl anti-HA magnetic beads (Thermo Fisher scientific, 88836) with end-over-end rotation for 16h at 4°C. The protein-bound beads were washed four times with fractionation buffer and proteins immunoprecipitated were reduced by the addition of 40μl of Laemmli sample buffer, heated at 95 °C for 5 min, and resolved on 12% SDS polyacrylamide gel electrophoresis (PAGE) gel, and analyzed by western blotting.

### Lysosome immunoprecipitation

Cells transiently expressing T192-3×HA were lysed, and intact lysosomes were immunoprecipitated using anti-HA-conjugated Dynabeads (Thermo Scientific, 88837). Briefly, the cells were seeded in a 6−well plate at a density appropriate for them to reach confluency after 24 h. The medium was removed, the cell monolayers were scraped into 10 ml KPBS (136 mM KCl and 10 mM KH2PO4, pH 7.25, containing protease (Roche, 1836153001) and phosphatase inhibitors (PhosSTOP Sigma, 04906845001) and collected by centrifugation at 1,500 r.p.m. for 5 min. The pelleted cells were resuspended in a total volume of 1 ml KPBS and ~4×10^6^ cells were mechanically broken by spraying 8−10 times through a 23G needle attached to a 1ml syringe, and then spun down at 2,000 r.p.m for 10min at 4°C, yielding a post nuclear supernatant (PNS). The post-nuclear supernatant was harvested, quantified using Bradford assay (Thermo-Fisher, 23246). Equal quantities of proteins from each condition were incubated with 50 µl anti-HA magnetic beads with end-over-end rotation for 1h at 4°C. The lysosome-bound beads were washed four times with KPBS. For immunoblotting analysis, anti-HA immunoprecipitates were reduced by the addition of 40μl of Laemmli sample buffer, heated at 95 °C for 5 min, and resolved on 12% SDS polyacrylamide gel electrophoresis (PAGE) gel, and analyzed by western blotting.

For cystine measurements on purified lysosomes, 50 μL of 80% MeOH was added to the beads bound to organelles and incubated for 5 mins at room temperature. The free amino acids were further derivatized before MS analysis. For whole cell lysate samples, 30 μL of supernatant were taken and mixed with 3 μL of 5 N NaOH in borate buffer to ensure the pH was below pH 9. For LysoIP, the samples were dissolved in 40 μL borate buffer from Waters. A mixture of 30 μL samples (both WCL and LysoIP), 5 μL borate buffer, 5 μL MSK-CAA-A2 internal standard mixture (1:5 dilution) and 10 μL AQC solution was added, giving a total volume of 50 μL and mixed thoroughly immediately after addition. After 10 mins incubation at 55°C, samples were loaded onto a Waters UPLC H-Class Plus system with an Acquity UPLC Qardupole Solvent Manager and Sample Manager. Derivatized amino acids were detected on a Waters QDa single quadrupole mass detector in positive mode. The Qda was operated by one MS, followed by selected ion monitoring (SIR) for each individual amino acid with defined retention window, shown in table below. The column temperature was maintained at 55 °C. The injection volume was 1 μL. Gradient elution was performed using 0.1% formic acid in water as eluent A and 0.1% formic acid in acetonitrile as eluent B. The flow rate was kept constant at 0.5 mL/ min with the following gradient (expressed as solvent B): Initial conditions: 1.0% B, 0.0-1min: 1% B, 1-4 min: 13.0% B, 4-8.5 min: 15.0% B, 8.5-9.5 min: 95.0% B, 9.5-11.5 min: 95% B, 12-15 min: 1% B. The data acquisition and data analysis were done by Masslynx 4.2 (Waters).

### Immunoblotting

Proteins were extracted from mouse tissues or cultured cells, lysed using a buffer containing protease (Roche) and phosphatase inhibitors (PhosSTOP Sigma), followed by sonication and centrifugation at 16,000*g* for 10 min at 4°C. The samples were thawed on ice, normalized for protein (20μg/lane), dissolved in Laemmli sample buffer and separated by SDS-PAGE under reducing conditions. After blotting onto PVDF and blocking with 5% non-fat milk (1706404, Bio-Rad Laboratories), the membranes were incubated overnight at 4°C with primary antibody, washed, incubated with peroxidase-labeled secondary antibody, and visualized with enhanced chemiluminescence (WBKLS0050, Millipore, Life technologies). Signal intensity was assessed by measuring the relative density of each band normalized to β-actin, GAPDH, γ-tubulin or α-tubulin with ImageJ software.

### Cell viability assay

The viability of mPTCs after drug treatments were assessed via MTT assay (ab211091, Abcam) in accordance with the manufacturer’s instructions. Briefly, the cells were washed three times with PBS and then incubated with 0.5 mg/ml of MTT diluted in cell media. After 4 h of incubation at 37°C and until appearance of intracellular purple formazan crystals, the remaining crystals were dissolved with dimethyl sulfoxide (276855, Sigma-Aldrich) or SDS, and the absorbance measured at 570nm.

### Cell growth and proliferation

To measure proliferation and growth, the cells were seeded in 24‒well plates at a density of 2.0 × 104 cells per well. The cells were cultured for 4 days, and cell medium was renewed daily. Where indicated, the cells were treated with Torin1 (16h at the indicated concentrations), then trypsinized every 24 h and quantified using the countess automated cell counter TC10 automated cell counter (BIO−RAD). The time–course experiments were repeated three times.

### Statistics and reproducibility

The plotted data were presented as mean ± standard error of the mean (SEM). Statistical comparisons between experimental groups were determined by using one-way analysis of variance (ANOVA) followed by Tukey’s or Sidak’s or Dunnett’s multiple comparison test, when appropriate. When only two groups were compared, two tailed unpaired or paired Student’s *t* tests were used as appropriate. The normality criteria (calculated by D’Agostino and Pearson *omnibus* normally test) were met. Non-parametric data were analysed using a Kruskal-Wallis test with Dunn’s multiple comparison correction. The levels of statistical significance are indicated by symbols, and the *P*-values are indicated in the figure legends along with the statistical tests. All experiments reported here were performed at least two to three times independently, unless otherwise indicated in the figure legends. The investigators were not blinded to allocation during the experiments and outcome assessment. Graph Pad Prism software v. 9.4.1 (GraphPad software) was used for generating all statistical analyses.

## Data availability

Source data and/or reagents supporting the findings of this study are available from corresponding authors, O.D. and A.L., upon reasonable request.

## ACKNOWLEDGEMENTS

We thank Nadine Nägele and Aleksandra Kokanovic for technical help, and Alessio Cremonesi and Benjamin Klormann for cystine measurements, and the Center for Microscopy and Image Analysis at the University of Zurich for providing equipment and confocal microscopy assistance, as well as the Functional Genomics Center at the University of Zurich for providing equipment and assistance with sample preparation and mass spectrometry-based analyses. We thank V. Berno, M. Stagi, R. Zoncu, C.W. Li and P. Nanni, S. Streb, and A. Othman for their advice and constructive suggestions, and A. Hall and I. Sakhi for providing the LysoTracker dye. We also acknowledge Euro-BioImaging (www.eurobioimaging.eu) for providing access to imaging technologies and services via the Italian Node (ALEMBIC, Milan, Italy). We are grateful to the Cystinosis Research Foundation (Irvine, CA, USA; project grants CRFS-2017-007 and CRFS-2020-005), the Swiss National Science Foundation (project grant 310030_189044), and the University Research Priority Program of the University of Zurich (URPP) ITINERARE - Innovative Therapies in Rare Diseases (OD and AL).

## AUTHOR CONTRIBUTIONS

A.L. and O.D. conceived and supervised the study. A.L., M.B., Z.C., B.P.F., and O.D. planned the experimental design. M.B. carried out lysosome immunopurification, RNA interference, gene expression experiments, cell imaging and molecular biology studies and analyzed the data. Z.C. generated the *ctns* knockout zebrafish model and other transgenic zebrafish reporter lines, and together with S.K. performed pharmacological rescue experiments in *ctns* zebrafish larvae and analyzed the data. B.P.F., M.B. and Z.C. carried out immunoblotting analyses of mTORC1 activity in mouse cells and in preclinical models (rat and zebrafish) of cystinosis, respectively. P.K. performed pharmacological rescue experiments in *Ctns* rats and analyzed the data. S.P. and E.D. analyzed the microarray data from *Ctns* mice. A.G and M.K. carried out drug target discovery using artificial intelligence and machine learning tools. E.L. carried out the multi-omics integration, analyzed the data, and performed multi-omics network building. A.L. and O.D. drafted the paper with inputs and comments from all the authors.

