## Supplemental Material for "Lysosomal cystine export regulates mTORC1 signaling to guide kidney epithelial cell fate specialization"

Marine Berquez et al.

### Supplementary information

Supplementary Figures 1-14

Supplementary Tables 1-2

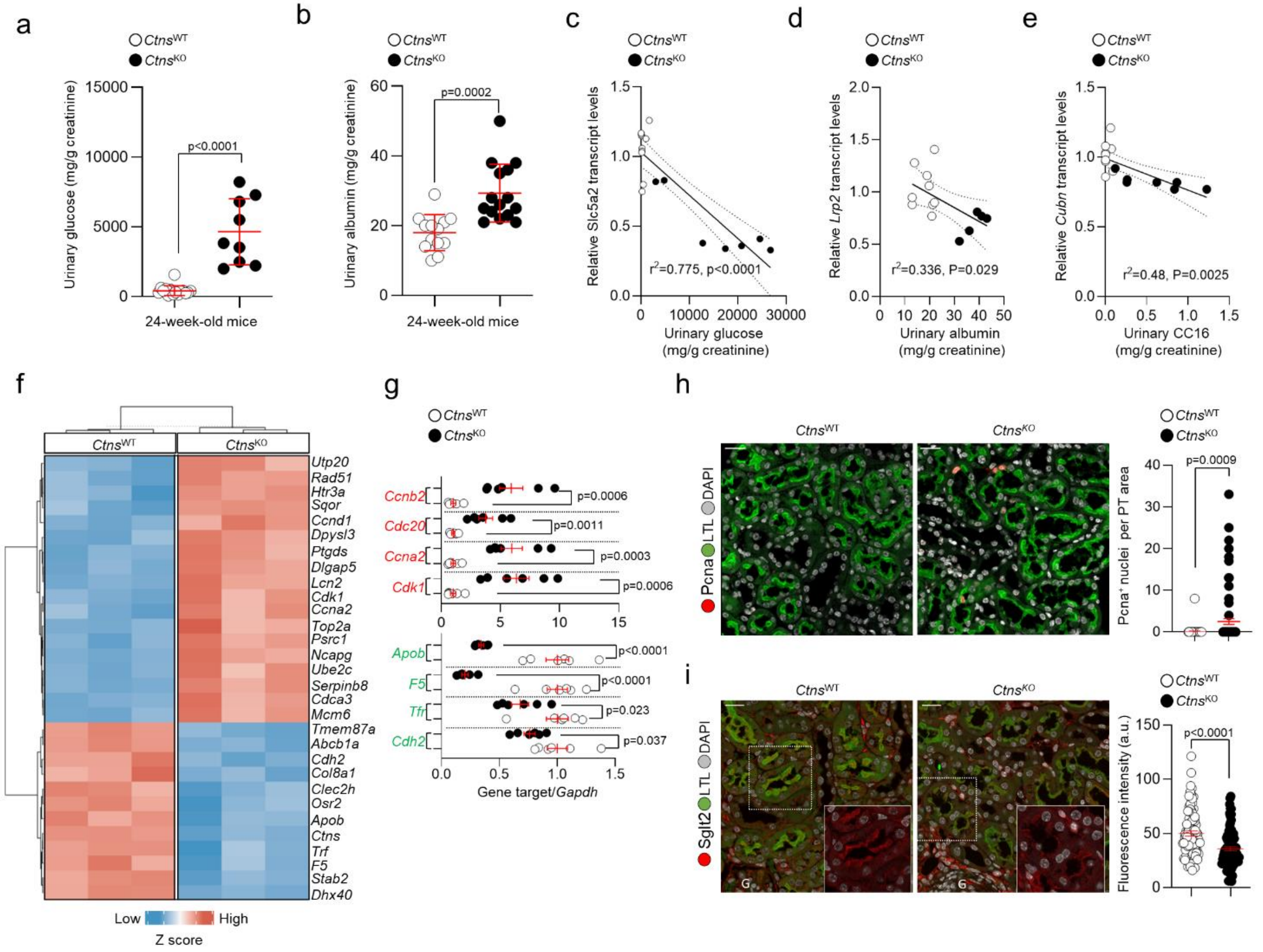


**Supplementary Figure 1. Differentiation defects and abnormal programs for growth and proliferation in the proximal tubule (PT) segments of CTNS-deficient mouse kidneys.** (**a**) Glucose and (**b**) albumin levels in the urine samples from *Ctns* mice; n>9 mice per genotype. (**c-e**) Scatter plots of transcript levels of (**c**) *Slc5a2*, (**d**) *Lrp2*, and (**e**) *Cubn* versus disease phenotype variables, including glucosuria, albuminuria, and LMW/CC16 proteinuria, with linear regression measured by Pearson correlation coefficient in the *Ctns*-deficient and control group. (**f**) Heatmap showing the expression of the top 30 differentially expressed genes (DEGs) in the PT segments obtained from 24-week-old Ctns-deficient mouse and (age-matched) control littermates; n=3 animals per each group. (**g**) Quantification of mRNA levels of the indicated genes; n=6 animals per group. (**h** and **i**) Immunofluorescence staining and quantification of (**h**) the number of Pcna-positive nuclei (red) or (**i**) mean fluorescence intensity of Sglt2 signal in the LTL (Lotus Tetragonolobus lectin)-positive PT segments (green) of mouse kidneys. Pcna: n=86 *Ctns*$^{WT}$ and n=90 *Ctns*$^{KO}$ PTs; Megalin: n=61 *Ctns*$^{WT}$ and *Ctns*$^{KO}$ PTs; data were pooled from 3 biologically independent replicates. Statistics calculated by unpaired two-tailed Student's *t* test. Plotted data represent mean ± SEM. Nuclei counterstained with DAPI (grey). Scale bars are 10 μm.

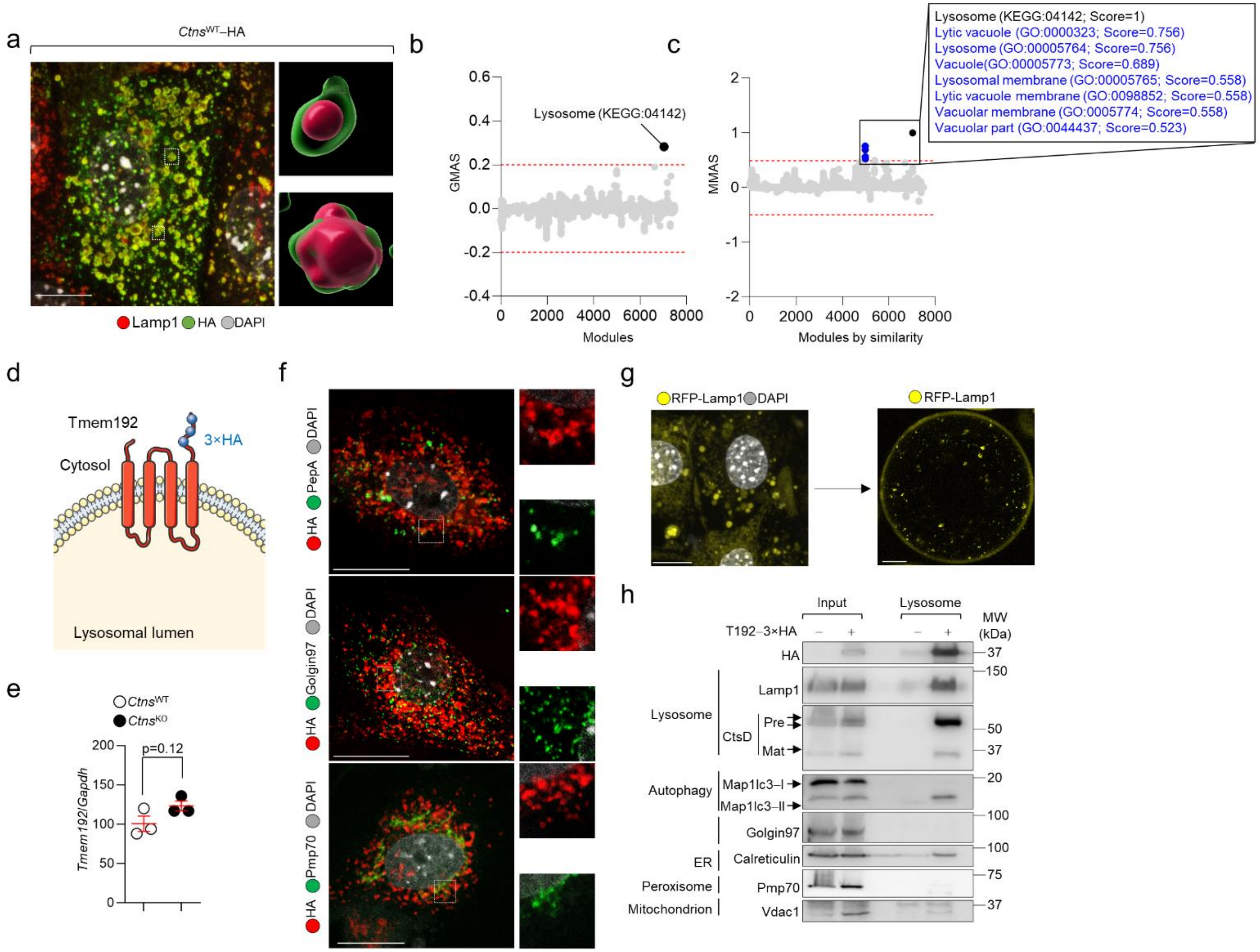


**Supplementary Figure 2. CTNS subcellularly localizes to the surface of the lysosome, and validation of the LysoIP workflow in the primary PT cells derived from *Ctns* mouse kidneys. (a)** mPTCs from *Ctns*$^{WT}$ mice were transduced with adenoviral particles transiently hemagglutinin-tagged CTNS$^{WT}$ (CTNS$^{WT}$-HA) for 2 days and immunostained for HA (green) and Lamp1 (red). Confocal microscopy and three-dimensional (3D) reconstruction of Z-stacked images of mPTCs confirmed the localization of CTNS at the surface of Lamp1-flagged lysosomes. (**b-c**) Systems biology-based tool (GeneBridge;

https://systems-genetics.org) was exploited to uncover additional functions of *Ctns* beyond cystine transport. This computational pipeline enables the gene's function through its (anti)correlation with other genes for which the Gene Ontology (GO) terms/ biological pathways are already known using large-scale mouse expression compendia. (**b**) Gene-module association analysis of Ctns and module-module association analysis of lysosome (KEGG:04142) module in mouse, using 252 and 269 mouse datasets. Red dashed line indicates the threshold of significant association in **b** and **c**. (**d**) Schematic showing the localization of 3×haemagglutinin (HA)-fused transmembrane protein 192 (Tmem192×3HA) at the surface of the lysosome. (**e**) mRNA levels of Tmem192 in *Ctns* mPTCs analyzed by RT-qPCR. (**f**) The cells expressing Tmem192×3HA were pulsed with Bodipy FL-PepstatinA (1μM at 37°C for 1h; green) and immunostained for HA (red), or for Golgin–97 (green) and HA (red), or for Pmp70 (green) and HA (red) and analyzed by confocal microcopy. (**g**) Confocal images of (right panel) mPTCs transiently expressing Lamp1-RFP-3×Flag and (left panel) bead-purified lysosomes following immunoisolation method. (**h**) Immunoblotting for protein markers of various subcellular compartments in whole-cell lysates and purified lysosomes. Lysates were prepared from cells expressing Tmem192×3HA; n=2 biologically independent experiments. Nuclei counterstained with DAPI (grey). Statistics calculated by unpaired two-tailed Student's t test. Plotted data represent mean ± SEM. Scale bars are 10mm in **a**, **f**, and **g**.

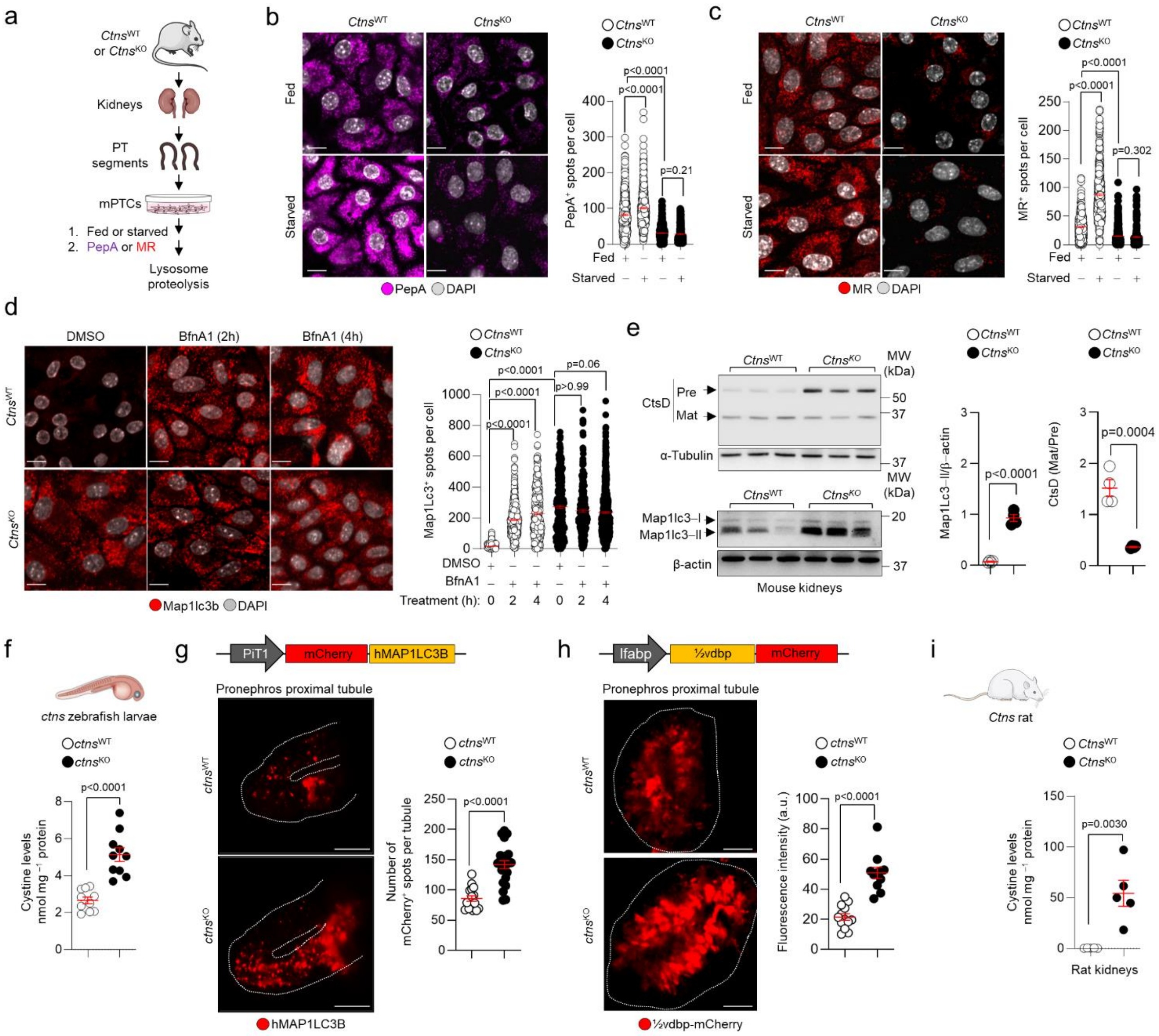
a
CtnsWT or CtnsKO
Kidneys
PT segments
mPTCs
1. Fed or starved
2. PepA or MR
Lysosome proteolysis
b
CtnsWT
CtnsKO
Fed
Starved
PepA DAPI
PepA+ spots per cell
p<0.0001
p<0.0001
p=0.21
Fed
Starved
c
MR DAPI
MR+ spots per cell
p<0.0001
p<0.0001
p=0.302
d
DMSO
BfnA1 (2h)
BfnA1 (4h)
Map1lc3b DAPI
Map1Lc3+ spots per cell
p<0.0001
p<0.0001
p<0.0001
p=0.06
p>0.99
DMSO
BfnA1
Treatment (h):
e
CtsD
Pre
Mat
MW (kDa)
α-Tubulin
Map1lc3–I
Map1lc3–II
β-actin
Mouse kidneys
Map1Lc3–II/β–actin
CtsD (Mat/Pre)
p<0.0001
p=0.0004
f
ctns zebrafish larvae
ctnsWT
ctnsKO
Cystine levels nmol mg−1 protein
p<0.0001
g
PiT1
mCherry
hMAP1LC3B
Pronephros proximal tubule
Number of mCherry+ spots per tubule
p<0.0001
hMAP1LC3B
h
lfabp
½vdbp
mCherry
Fluorescence intensity (a.u.)
p<0.0001
½vdbp-mCherry
i
Ctns rat
p=0.0030
Rat kidneys

**Supplementary Figure 3. CTNS loss and the resulting cystine storage compromise the catabolic activities of lysosomes in mPTCs, rat, and zebrafish lacking CTNS**. (**a**) mPTCs were cultured under fed and starved conditions for 4h, pulsed for 1h with (**b**) Bodipy FL-PepA (PepA, 1μM at 37°C) or with (**c**) MagicRed Cathepsin B substrate (MR, 1μM at 37°C) and analysed by confocal microscopy. Representative confocal images and quantification of the number of PepA- and MR-positive structures per cell; n> 362 cells (MR) and n>359 cells (PepA) pooled from two biologically independent experiments. (**d**) mPTCs were treated with non-saturating concentrations of BafilomycinA1 (BfnA1; 250nM for 4h) at the indicated times. The cells were immunostained for Map1Lc3b (red) and analyzed by confocal microscopy. Representative confocal images and quantification of the number of Map1lc3b-positive structures per cell (n>289 cells per each condition pooled from two biologically independent experiments). (**e**) Immunoblot and quantification of the indicated proteins in whole-kidney samples from *Ctns*$^{KO}$ relative to control mice; n=4 mice per each genotype. (**f** and **i**) Quantification of cystine levels by HPLC (**f**) in 5-dpf-*ctns* zebrafish larvae (n=10 biologically independent samples per each genotype, with each dot representing a pool of 6 zebrafish) and (**i**) in 16-week-old Ctns rat kidneys (n=5 animals per each genotype). (**g**) The *ctns* zebrafish were outcrossed with zebrafish expressing *PiT1::mCherry-hMAP1LC3B* (autophagosome marker, red) in the pronephric proximal tubule, and analysed by multiphoton microscopy. Representative images and quantification of the mean fluorescence intensity and numbers of mCherry-positive structures per pronephros (n= 13 wild-type and n=11 knockout zebrafish). (**h**) The *ctns* zebrafish were outcrossed with zebrafish expressing the ½vdbp-mCherry in the liver and analysed by multiphoton microscopy. Representative images and quantification of the number of mCherry-positive structures per pronephric tubule (n=16 *ctns*$^{WT}$ and n=23 *ctns*$^{KO}$ zebrafish). Plots represent mean ± SEM. Statistical calculated by unpaired two-tailed Student's t test in **e**, **f**, **g**, **h,** and **i**. Statistics calculated by Kruskal-Wallis followed by Dunn's multiple comparisons test in **b**, **c** and **d**. Nuclei counterstained with DAPI (grey). Scale bars are10μm.

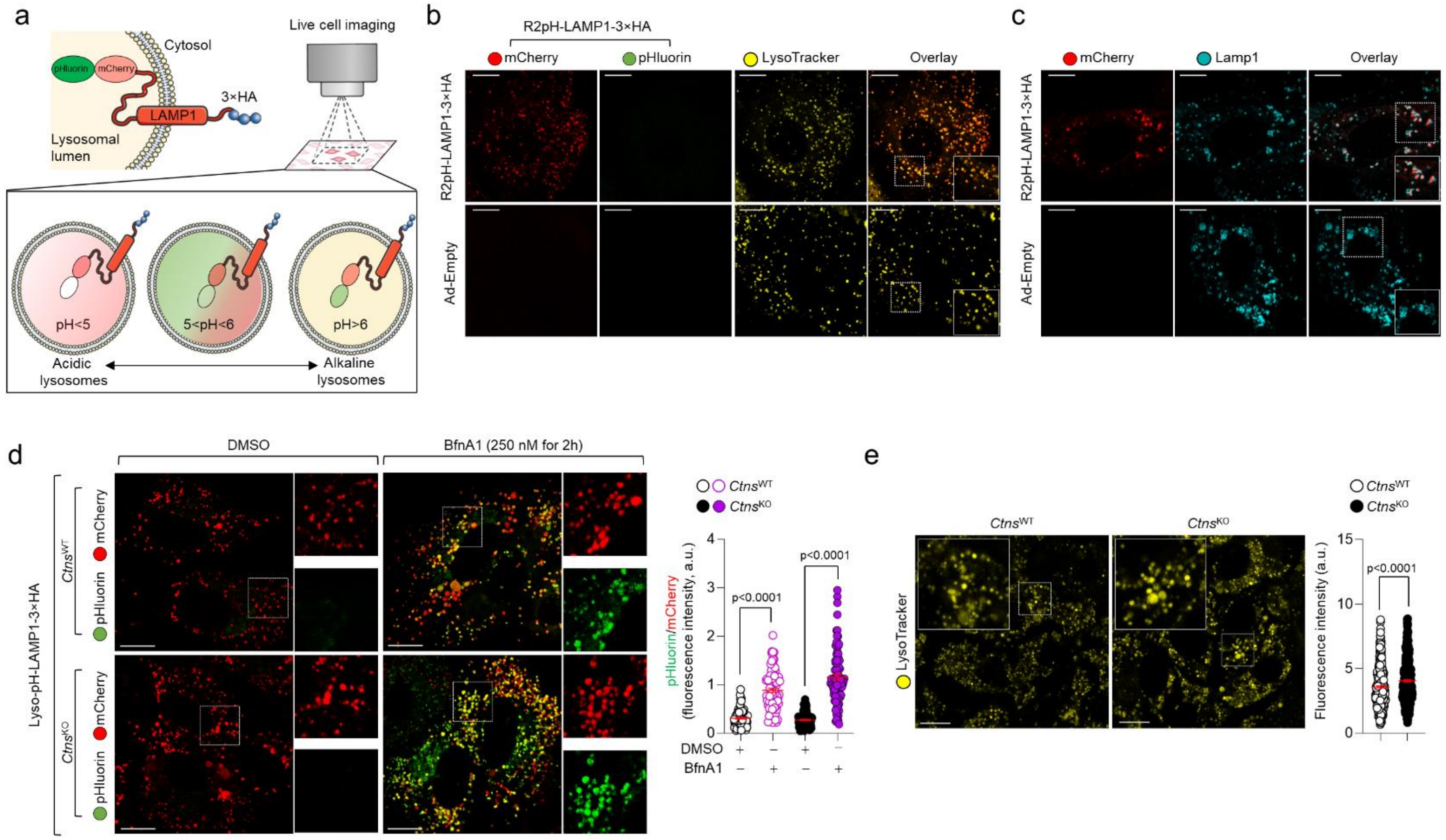


**Supplementary Figure 4. CTNS loss and cystine storage does not affect the lysosomal acidification in the tubular cells.** (**a**) Schematic showing the design and subcellular localization of a genetically encoded, bona fide biosensor that uses a pHluorin-mCherry linked to the luminal domain of the lysosome-localized protein, mouse Lamp1, with a cytosolic 3×HA tag [hereafter, called ratiometric R2pH-LAMP1-3×HA], to reliably measure the changes in pH homeostasis within the lumen of the lysosome. When the lysosomes are properly acidified, the acidic environment triggers the quenching of the pHluorin signal, leaving only (acidic insensitive) red fluorescence. In an acidic lysosome, the biosensor has a low ration value and is displayed in red, while it has a high ratio value in alkaline lysosomes and is displayed in yellow. (**b**) The *Ctns* mPTCs were transduced with an adenovirus that transiently expresses R2pH-LAMP1-3×HA for 24 h and loaded with Lysotracker dye (1μM for 4 hours; yellow) or (**c**) fixed and immunostained for Lamp1 (cyan) and analyzed by confocal microscopy. These functional assays showed the colocalization of the biosensor with LysoTracker- and Lamp1-positive lysosomes. (**d**) The Ctns mPTCs expressing R2pH-LAMP1-3×HA were treated with non-saturating concentrations of BafilomycinA1 (BfnA1; 250nM) for 4h and analyzed by confocal microscopy. Confocal images and quantification of green-to-red fluorescence intensity ratio per cell (n>88 cells per condition pooled from three biologically independent experiments). (**e**) The *Ctns* mPTCs were loaded with LysoTracker (1μM for 4 hours; yellow) and analysed by confocal microscopy. Representative confocal images and quantification of mean fluorescence intensity of LysoTracker signal in Ctns mPTCs (n>372 cells pooled from two biologically independent experiments). Plots represent mean ± SEM. Statistics calculate by one-way ANOVA followed by Sidak's multiple comparisons test in **d**, and by unpaired two-tailed Student's *t* test in **e**. Scale bars are 10μm.

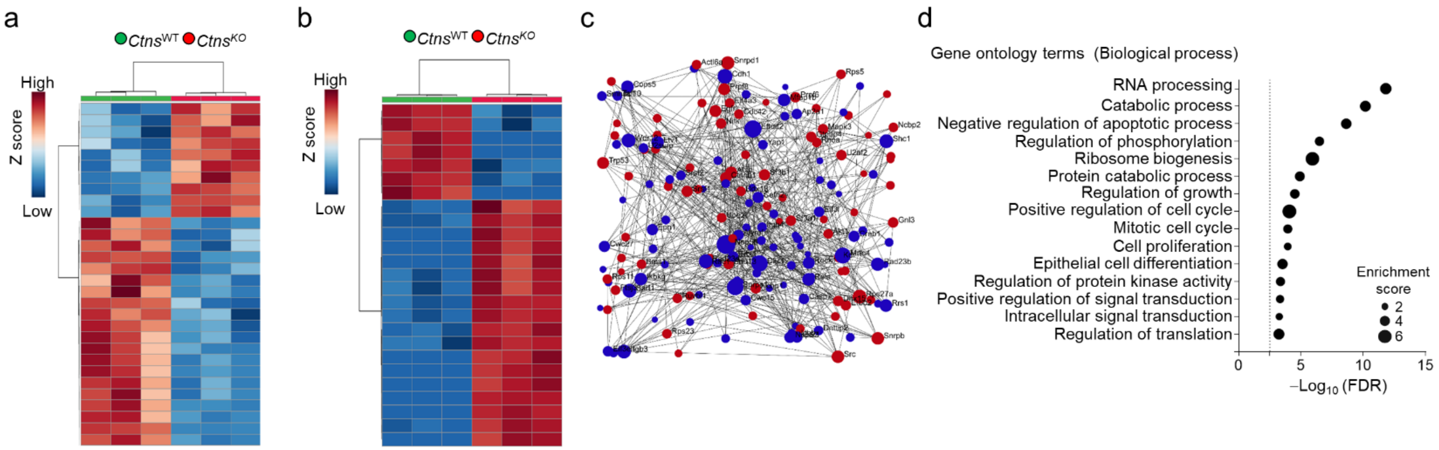


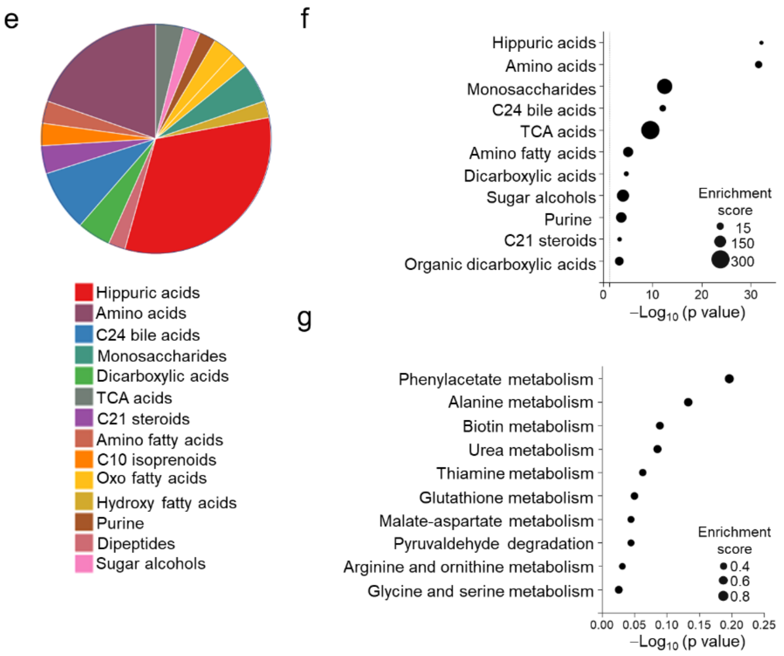


| Set name | Metabolites | References |
|---|---|---|
| Amino acids | Betaine; Arginosuccinic acid; Glycine; Phenylalanine; L-Alanine; L-threonine; L-Isoleucine; L-Lysine; L-Cystine; Ornithine; Sarcosine; L-Arginine; L-Leucine; L-Homoserine; Ureidosuccinic acid; L-Valine; D-Alanine; Nicotinuric acid; D-Ornithine; D-Arginine; β−Leucine; L-Allothreonine; 5-L-Glutammyl-taurine; Norvaline; γ−Glutamyl−β−aminoproprionitrile | RefMet |

| Set name | Metabolites | References |
|---|---|---|
| Glutathione metabolism | Cysteinlglycine; Glycine; Glutathione; L−Glutamic acid; L−Alanine; NADP; NADPH; Pyroglutamic acid; Adenosine triphosphate; L-cysteine; γ−Glutamylcysteine; FAD; Zinc (II) ion; ADP; Water; Phosphoric acid; Hydrogen peroxide; Oxidized glutathione; γ−Glutamylalanine; Zinc; Bromomethane | SMPDB KEGG |

**Supplementary Figure 5. Proteomics and metabolomics profiling in the tubular cells lacking CTNS and cystine storage.** (**a**-**b**) Heatmap showing the expression of the top 30 differentially expressed proteins (**a**) and metabolites (**b**) in *Ctns* mPTCs derived from the kidneys of 24-week-old mice; n=3 animals per each group. (**c**) Protein-protein interaction map and (**d**) overrepresentation analysis of top biological processes (Gene Ontology, GO terms; Biological process) associated with differentially expressed proteins [$log_2FC < -0.5$ and $Log_2FC > 1$, p-value < 0.05] using the Network Analyst toolkit (www.networkanalyst.ca). This web-based biological network analysis project and visualize gene/proteins within their biological networks to explore their statistical and functional relationships. The computational tool embeds genes/proteins within biological networks from 15 different databases. These networks are mined to extract the genes/proteins, miRNA, drugs, chemicals, or diseases that have the strongest relationships to genes/proteins in the uploaded list (seed genes). Black dashed line indicates the threshold of significant enrichment. Black circles indicate the size of the enrichment score. (**e**) Pie charts showing the distribution of sub-chemical metabolite sets and (**f**) metabolite set and (**g**) pathway enrichment analyses of differentially produced metabolites [$log_2FC < -0.5$ and $Log_2FC > 1$, p-value < 0.05]. Black circles indicate the size of the enrichment score.

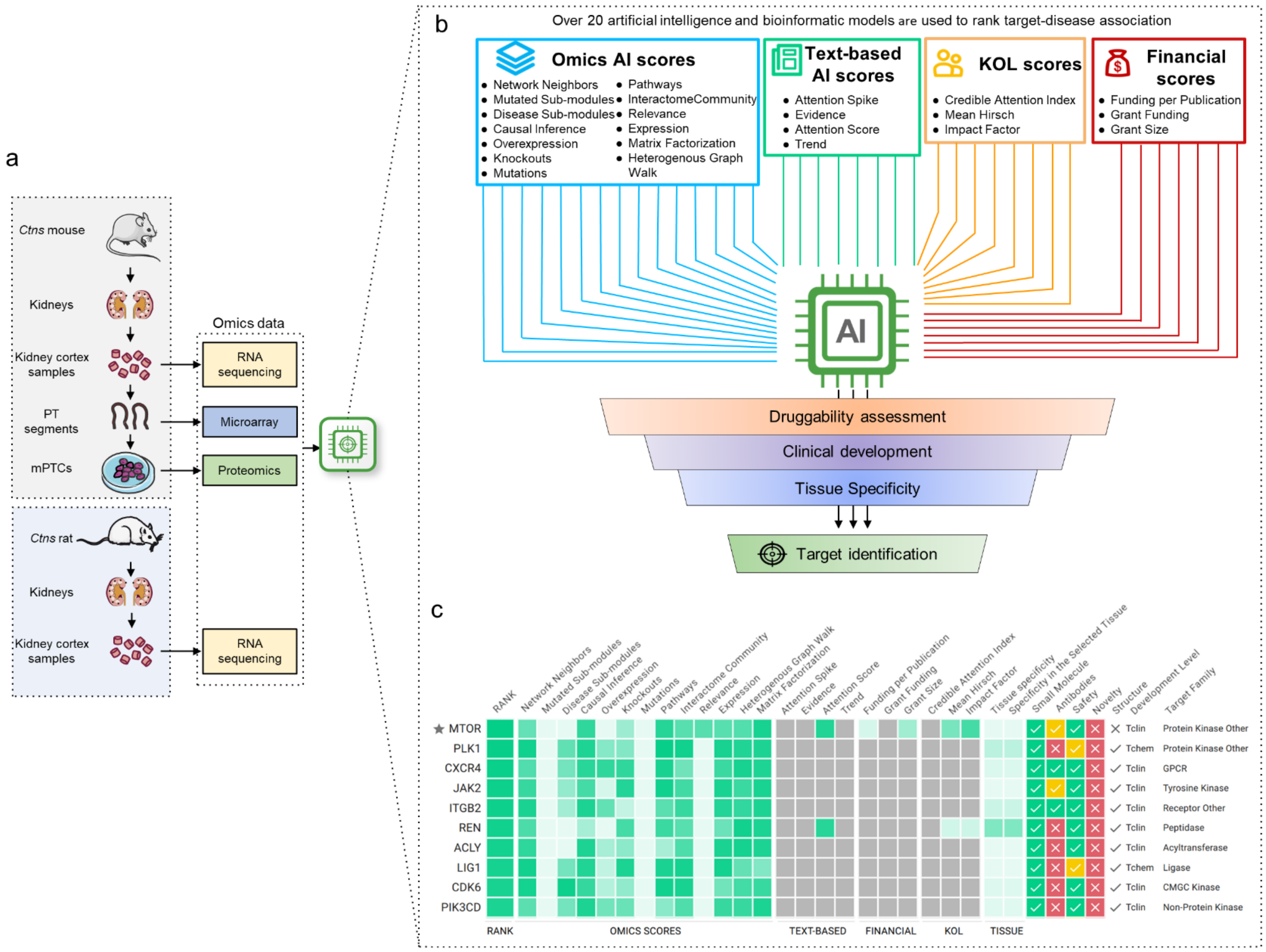
a
Ctns mouse
Kidneys
Kidney cortex samples
PT segments
mPTCs
Ctns rat
Kidneys
Kidney cortex samples
Omics data
RNA sequencing
Microarray
Proteomics
RNA sequencing
b
Over 20 artificial intelligence and bioinformatic models are used to rank target-disease association
Omics AI scores
Network Neighbors
Mutated Sub-modules
Disease Sub-modules
Causal Inference
Overexpression
Knockouts
Mutations
Pathways
InteractomeCommunity
Relevance
Expression
Matrix Factorization
Heterogenous Graph Walk
Text-based AI scores
Attention Spike
Evidence
Attention Score
Trend
KOL scores
Credible Attention Index
Mean Hirsch
Impact Factor
Financial scores
Funding per Publication
Grant Funding
Grant Size
AI
Druggability assessment
Clinical development
Tissue Specificity
Target identification
c
RANK
Network Neighbors
Mutated Sub-modules
Disease Sub-modules
Causal Inference
Overexpression
Knockouts
Mutations
Pathways
Interactome Community
Relevance
Expression
Heterogenous Graph Walk
Matrix Factorization
Attention Spike
Evidence
Attention Score
Trend
Funding per Publication
Grant Funding
Grant Size
Credible Attention Index
Mean Hirsch
Impact Factor
Tissue specificity
Specificity in the Selected Tissue
Small Molecule
Antibodies
Safety
Novelty
Structure
Development Level
Target Family
MTOR
PLK1
CXCR4
JAK2
ITGB2
REN
ACLY
LIG1
CDK6
PIK3CD
Tclin
Tchem
Protein Kinase Other
GPCR
Tyrosine Kinase
Receptor Other
Peptidase
Acyltransferase
Ligase
CMGC Kinase
Non-Protein Kinase
RANK
OMICS SCORES
TEXT-BASED
FINANCIAL
KOL
TISSUE

**Supplementary Figure 6. Artificial intelligence (AI)-powered pipeline to identify targets in CTNS-deficient/cystinosis-affected tubular cells.** An end-to-end artificial intelligence engine uses machine learning (ML) tools and statistical validation to rank disease-target associations and prioritize actionable (drug) targets in the tubular cells lacking CTNS and accumulating cystine. The AI-powered target discovery (PandaOmics) platform extracts disease knowledge from a variety of sources that include (**a**) multi-omics landscapes, here derived from transcriptomics signatures in kidney cortex samples obtained from *Ctns*$^{KO}$ mouse and rat or from proteomics profiling of primary PT cells derived from *Ctns* mouse kidneys, (**b**) text-curated data (patents, grants, publications, clinical trials, and key opinion leaders). The combination of described scores results in a ranked list of targets proposed for a given disease and can be filtered out through the assessment of druggability properties that include safety, novelty, accessibility by approved/investigational small molecules or biologics, and tissue specificity. (**c**) Heat map showing the top 10 most promising (drug) targets reversing dysregulated homeostasis in CTNS-deficient/cystinosis-affected tubular cells. Notably, mTOR has been identified as one of the top hits among the list of predicted targets.

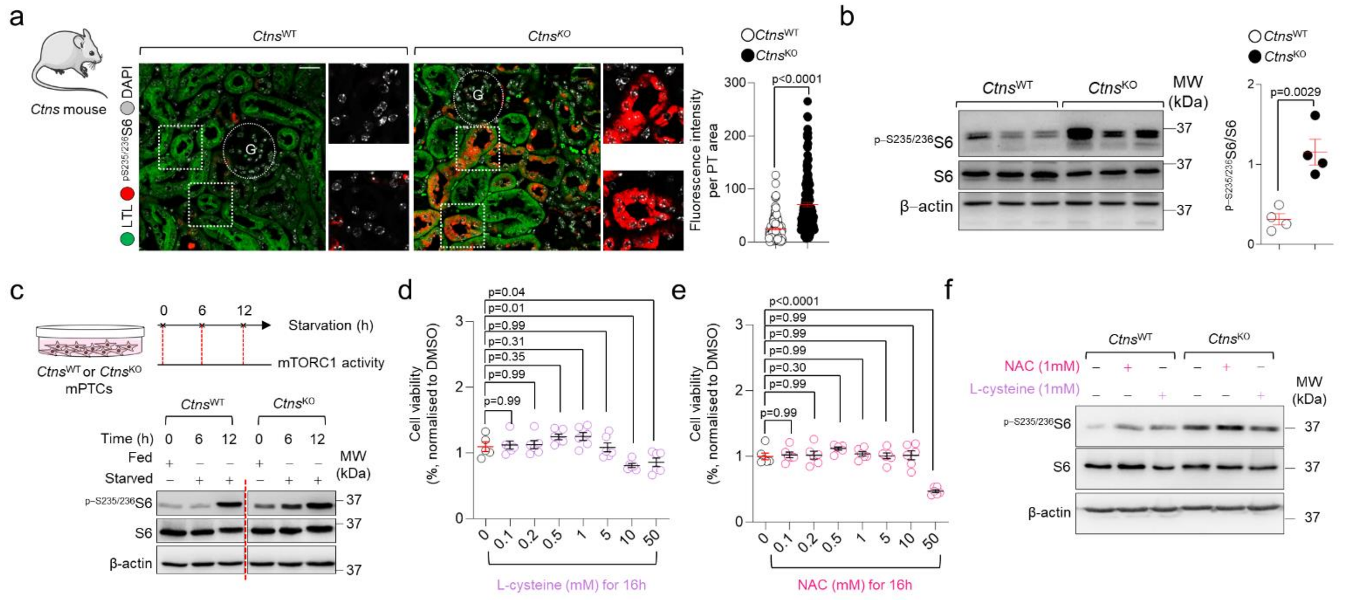


**Supplementary Figure 7. mTORC1 activity in *Ctns* mouse kidneys and derived PT cells.** (**a**) Representative confocal images and quantification of mean fluorescence intensity of phosphorylated $^{pS235/236}$S6 in LTL$^{+}$ (Lotus Tetragonolobus Lectin, green) PT segments of *Ctns* mouse kidneys (n>131 mouse PTs pooled from 3 animals per each group). (**b**) Immunoblot and quantification of the indicated proteins in whole-kidney samples from 24-week-old *Ctns* mice (n=4 mice per genotype). (**c**) The *Ctns* mPTCs were starved for the indicated times and cell lysates were immunoblotted for the indicated proteins and phosphoproteins; n=2 biologically independent experiments. (**d** and **e**) The *Ctns*$^{WT}$ mPTCs were treated with (**d**) L-cysteine or (**e**) (N-acetyl cysteine, NAC) at the indicated concentrations. After 16h treatment, the cell viability was assessed by using the MTT assay. The cell viability has been normalized over the mean of the corresponding control (n>5 wells pooled from two biologically independent experiments). (**f**) The *Ctns* mPTCs were treated with non-toxic concentrations of L-cysteine (1mM) and NAC (1mM) for 16h. Cell lysates were immunoblotted for the indicates proteins and phosphoproteins; n=2 biologically independent experiments. Plots represent mean ± SEM. Statistical calculated by unpaired two–tailed Student's *t* test in **a** and **b** and by one-way ANOVA followed by Dunnett's multiple comparisons test in **d** and **e**. Scale bars are 50μm.

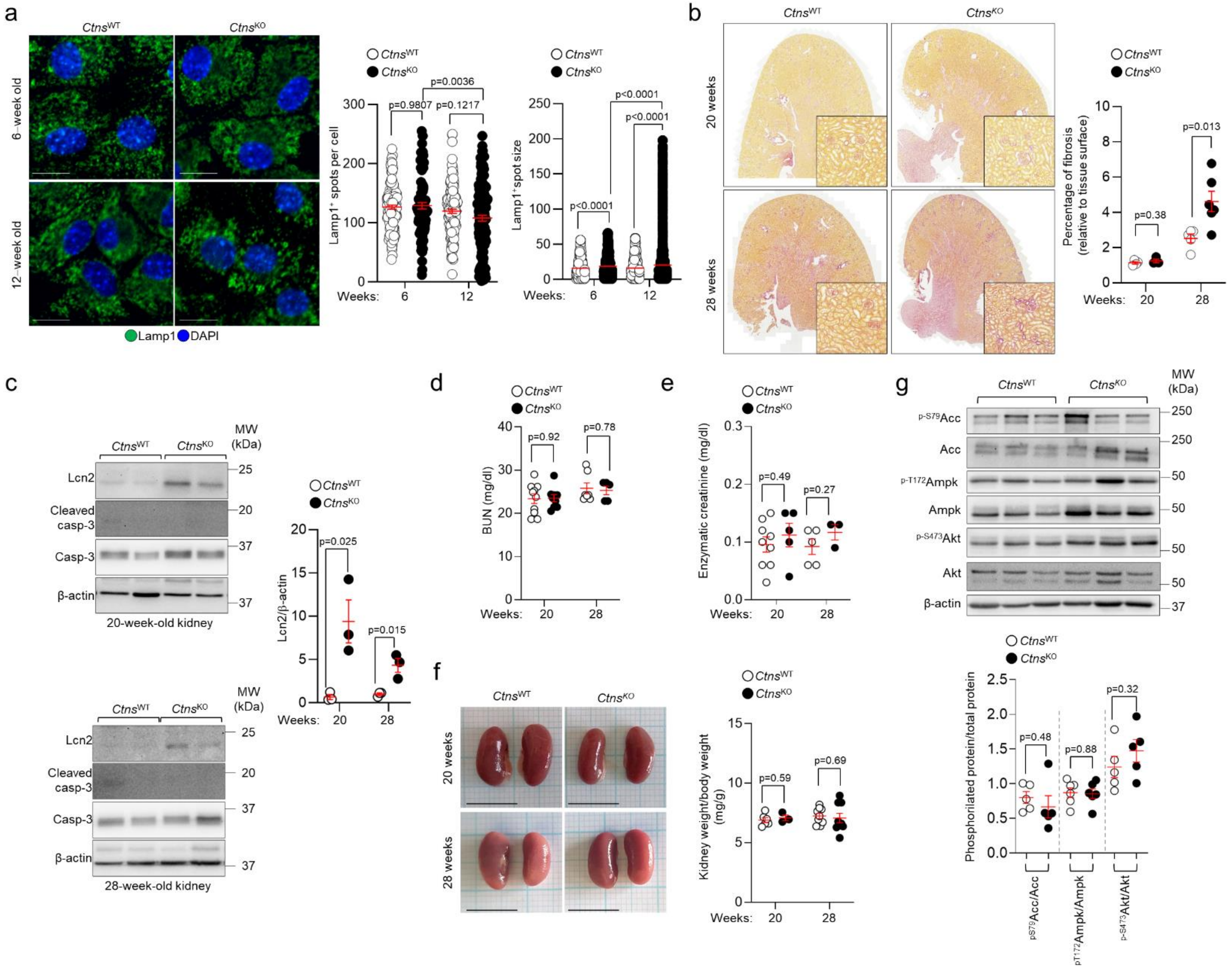
a
CtnsWT
CtnsKO
6-week old
12-week old
Lamp1 DAPI
Lamp1+ spots per cell
p=0.9807
p=0.1217
p=0.0036
Weeks: 6 12
Lamp1+spot size
p<0.0001
Weeks: 6 12
b
20 weeks
28 weeks
Percentage of fibrosis (relative to tissue surface)
p=0.38
p=0.013
Weeks: 20 28
c
Lcn2
Cleaved casp-3
Casp-3
β-actin
MW (kDa)
20-week-old kidney
28-week-old kidney
Lcn2/β-actin
p=0.025
p=0.015
Weeks: 20 28
d
BUN (mg/dl)
p=0.92
p=0.78
Weeks: 20 28
e
Enzymatic creatinine (mg/dl)
p=0.49
p=0.27
Weeks: 20 28
f
20 weeks
28 weeks
Kidney weight/body weight (mg/g)
p=0.59
p=0.69
Weeks: 20 28
g
p-S79Acc
Acc
p-T172Ampk
Ampk
p-S473Akt
Akt
β-actin
MW (kDa)
Phosphoriated protein/total protein
p=0.48
p=0.88
p=0.32
pS79Acc/Acc
pT172Ampk/Ampk
p-S473Akt/Akt

**Supplementary Figure 8. Cellular alterations and phenotypic changes in *Ctns* mouse kidneys. (a**) mPTCs derived from *Ctns* mice at 6 or 12 weeks of age were immunostained for the endogenous Lamp1 and analysed by confocal microscopy. Representative images and quantification of the number of Lamp1$^+$ structures and size. Number of Lamp1$^+$ spots: n>96 cells per each condition; Size of Lamp1$^+$ spots: n>13593 lysosomes per each condition, pooled from two biologically independent experiments. (**b**) Representative micrographs of Picrosirius Red staining and quantification of fibrosis in the kidneys of 20- and 28-week-*Ctns* mice (n=4 $Ctns^{WT}$ and n=3 $Ctns^{KO}$ at 20 weeks of age; n=5 $Ctns^{WT}$ and n=6 $Ctns^{KO}$ at 28 weeks of age). Insets: high magnification of the corresponding section. (**c**) Immunoblotting and quantification of the indicated proteins (n=3 mice per each genotype). (**d** and **e**) Quantification of (**d**) blood urea nitrogen (BUN) levels and (**e**) enzymatic creatinine levels measured from plasma samples of *Ctns* mice at the indicated ages (BUN: n=10 $Ctns^{WT}$ and n=9 $Ctns^{KO}$ at 20 weeks of age; n=7 $Ctns^{WT}$ and n=5 $Ctns^{KO}$ at 28 weeks of age; enzymatic creatinine: n=9 $Ctns^{WT}$ and n=5 $Ctns^{KO}$ at 20 weeks of age; n=5 $Ctns^{WT}$ and n=3 $Ctns^{KO}$ at 28 weeks of age). (**f**) Pictures of *Ctns* mouse kidneys and ratio of kidney weight to body weight at the indicated ages (n=6 $Ctns^{WT}$ and n=3 $Ctns^{KO}$ at 20 weeks of age; n=9 $Ctns^{WT}$ and $Ctns^{KO}$ at 28 weeks of age). Each dot of the graph represents the average of both kidneys derived from one mouse. (**g**) Immunoblotting and quantification of the indicated proteins and phosphoproteins (n>5 mice per genotype). Plotted data represent mean ±SEM. Statistical calculated by and by one-way ANOVA followed by Sidak's multiple comparisons test in **a** and unpaired two-tailed Student's *t* test in **b-g**. Scale bars are 10µm.

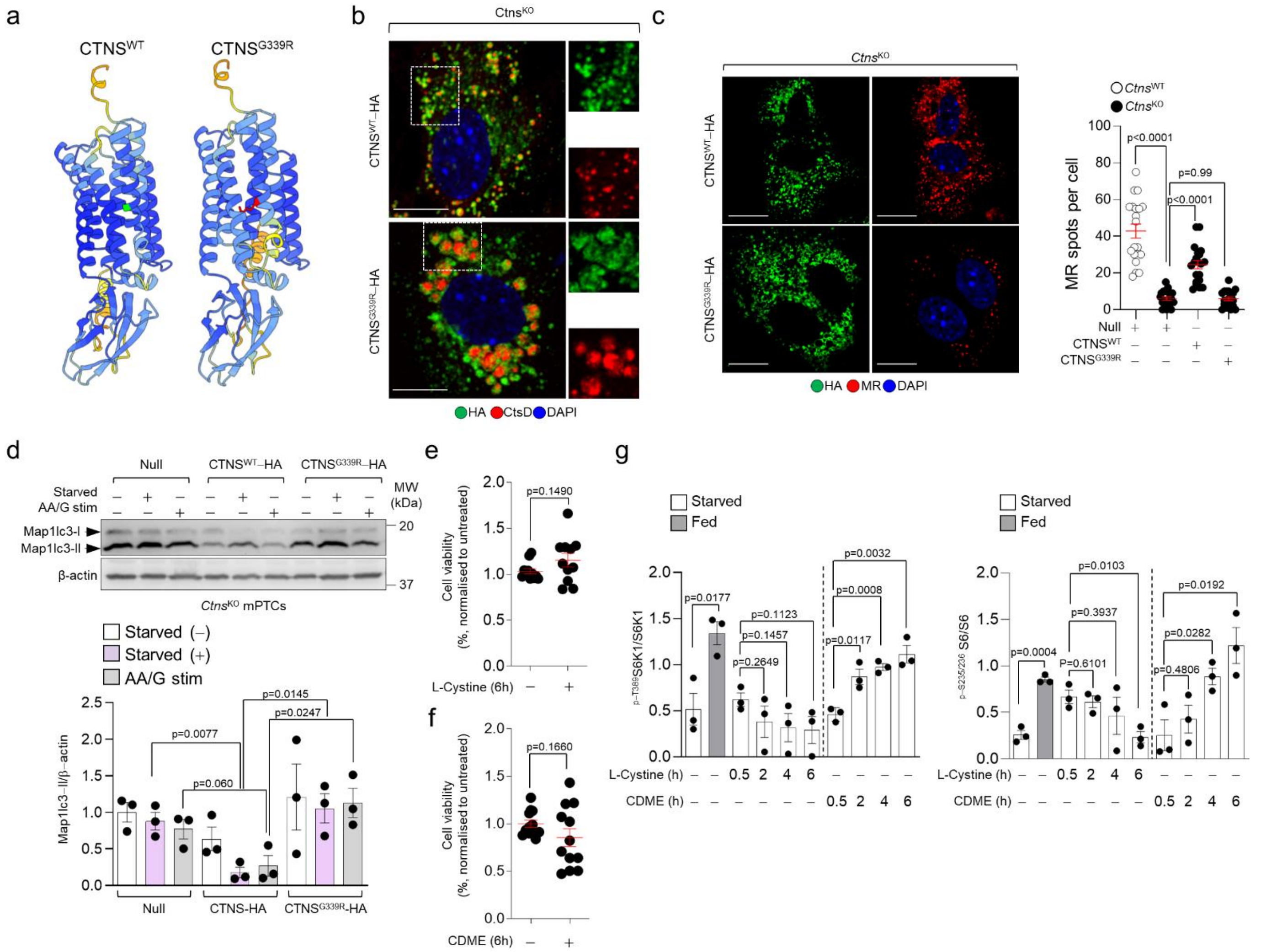


**Supplementary Figure 9. Changes in lysosomal cystine regulate mTORC1 in kidney tubular cells.** (**a**) Protein-structure prediction of CTNS$^{WT}$ and CTNS$^{G339R}$ by using the neural network AlphaFold. (**b-c**) mPTCs from *Ctns*$^{KO}$ mice were transduced with Null or HA-tagged wild-type CTNS (CTNS$^{WT}$-HA) or HA-tagged mutant CTNS (CTNS$^{G339R}$-HA) bearing adenoviral particles for 2 days. (**b**) Cells were immunostained for the epitope HA (green) and the endogenous cathepsin D (CtsD, red), and analyzed by confocal microscopy. Dotted white squares contain images at high magnification. (**c**) Cells were pulsed for 1h with MagicRed Cathepsin B (MR, 1μM at 37°C) substrate and immunostained for HA and analyzed by confocal microscopy. Representative images and quantification of MR-positive structures per cell (n=20 cells expressing HA per each condition pooled from two biologically independent experiments). (**d**) The *Ctns* mPTCs were cultured under fed or starved conditions or restimulated by adding back amino acids and glucose (AA/G stim). Immunoblotting and quantification of the indicated proteins in *Ctns*$^{KO}$ mPTCs; n=3 biologically independent experiments. (**e**-**g**) Starved *Ctns*$^{WT}$ mPTCs were stimulated with cystine dimethyl ester (CDME, 0.1 mM) or with unmodified L-cystine (0.1 mM) for the indicated times. (**e**-**f**) MTT assay monitoring cell viability in *Ctns* mPTCs exposed to L-cystine and CDME for the indicated times. Cell viability has been normalized over the mean of the corresponding control. n≥11 wells pooled from two biologically independent experiments. (**g**) Quantification of immunoblotting shown in Fig. 5h. Plotted data represent mean ± SEM. Statistical calculated by unpaired two–tailed Student's *t* test in **d**-**g** and by one-way ANOVA followed by Tukey's multiple comparisons test in **c**. Nuclei counterstained with DAPI (blue). Scale bars are 10μm.

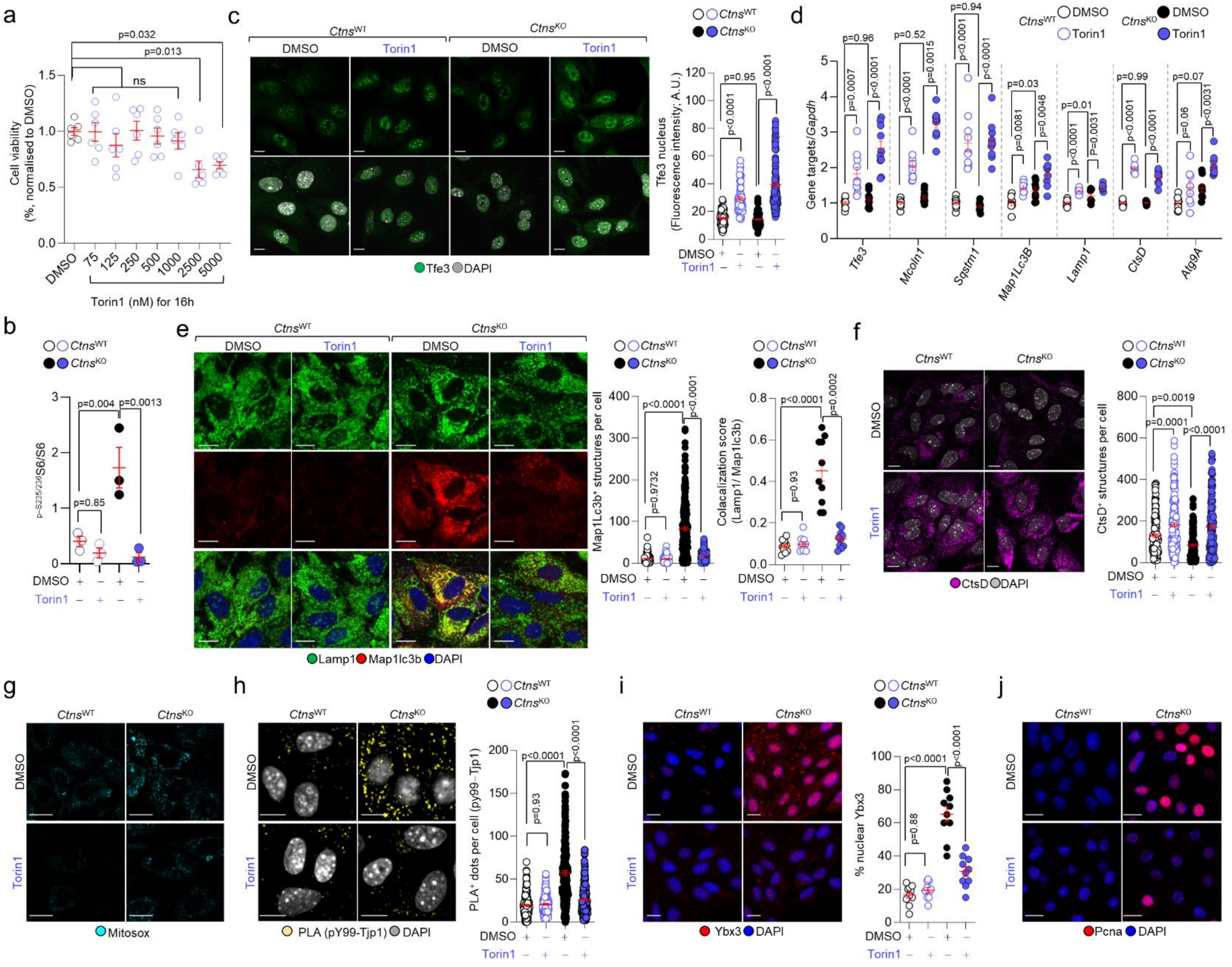
a
p=0.032
p=0.013
ns
Cell viability (%, normalised to DMSO)
DMSO
75
125
250
500
1000
2500
5000
Torin1 (nM) for 16h
c
CtnsWT
CtnsKO
DMSO
Torin1
Tfe3 DAPI
Tfe3 nucleus (Fluorescence intensity; A.U.)
p=0.95
p<0.0001
d
Gene targets/Gapdh
p=0.96
p=0.0007
p=0.52
p=0.0015
p=0.94
p=0.03
p=0.0081
p=0.0046
p=0.01
p=0.0031
p=0.99
p=0.07
p=0.06
Tfe3
Mcoln1
Sqstm1
Map1Lc3B
Lamp1
CtsD
Atg9A
b
p-S235/236S6/S6
p=0.004
p=0.0013
p=0.85
e
Lamp1 Map1lc3b DAPI
Map1Lc3b+ structures per cell
p=0.9732
Colacalization score (Lamp1/ Map1lc3b)
p=0.93
p=0.0002
f
CtsD DAPI
CtsD+ structures per cell
p=0.0019
g
Mitosox
h
PLA (pY99-Tjp1) DAPI
PLA+ dots per cell (py99-Tjp1)
i
Ybx3 DAPI
% nuclear Ybx3
p=0.88
j
Pcna DAPI

**Supplementary Figure 10. The suppression of mTORC1 signaling repairs lysosome, growth/proliferation, and dedifferentiation defects induced by CTNS loss/cystine storage.** (**a**) MTT assay measuring cell viability in Torin1-treated *Ctns*$^{WT}$ mPTCs. The cell viability has been normalized over the mean of the corresponding DMSO control (n=6 wells pooled from two biologically independent experiments). (**b**-**h**) mPTCs were treated with DMSO or Torin1 (250nm) for 16h. (**b**) Quantification of the immunoblotting shown in Fig. 6a. (**c**) Representative confocal images and quantification of mean fluorescence intensity of nuclear TFE3 signal (n> 69 cells per each condition). (**d**) mRNA levels of the indicated genes (n>7 samples per group, n=3 biologically independent experiments). (**e**) Representative confocal micrographs and quantification of the number of Lc3$^{+}$ puncta per cell and Map1lc3-Lamp1 colocalization (Map1Lc3b$^{+}$ structures: n>229 cells; colocalization score: n=10 randomly selected and non-overlapping fields of views per each condition). (**f**) Representative confocal images and quantification of the number of Cath D-positive structures per cell (n>137 cells per conditions). (**g**) The *Ctns* mPTCs were loaded with MitoSOX (mitochondrial ROS probe; 2.5μM for 10 min at 37 °C) and analysed by confocal microscopy. Representative confocal images of the quantification shown in fig 6e. (**h**) Confocal analysis of endogenous phosphorylation of Tjp1 by proximity ligation assay (PLA) and quantification of PLA$^{+}$ puncta per cell (n>154 cells per each condition). (**i**) Representative confocal images and quantification of the number Ybx3+ nuclei (n=10 non-overlapping fields of views per each condition). (**j**) Representative confocal images of the quantification shown in fig. 6g. Nuclei counterstained with DAPI (grey or blue). Plotted data represent mean ± SEM. Statistics calculated by one-way ANOVA followed Dunnett's multiple comparisons test in **a**, by one-way ANOVA followed Sidak's multiple comparisons test in **b-d** and **f**-**i,** by one-way ANOVA followed Tukey's multiple comparisons test in **e.** Scale bars are 10μm.

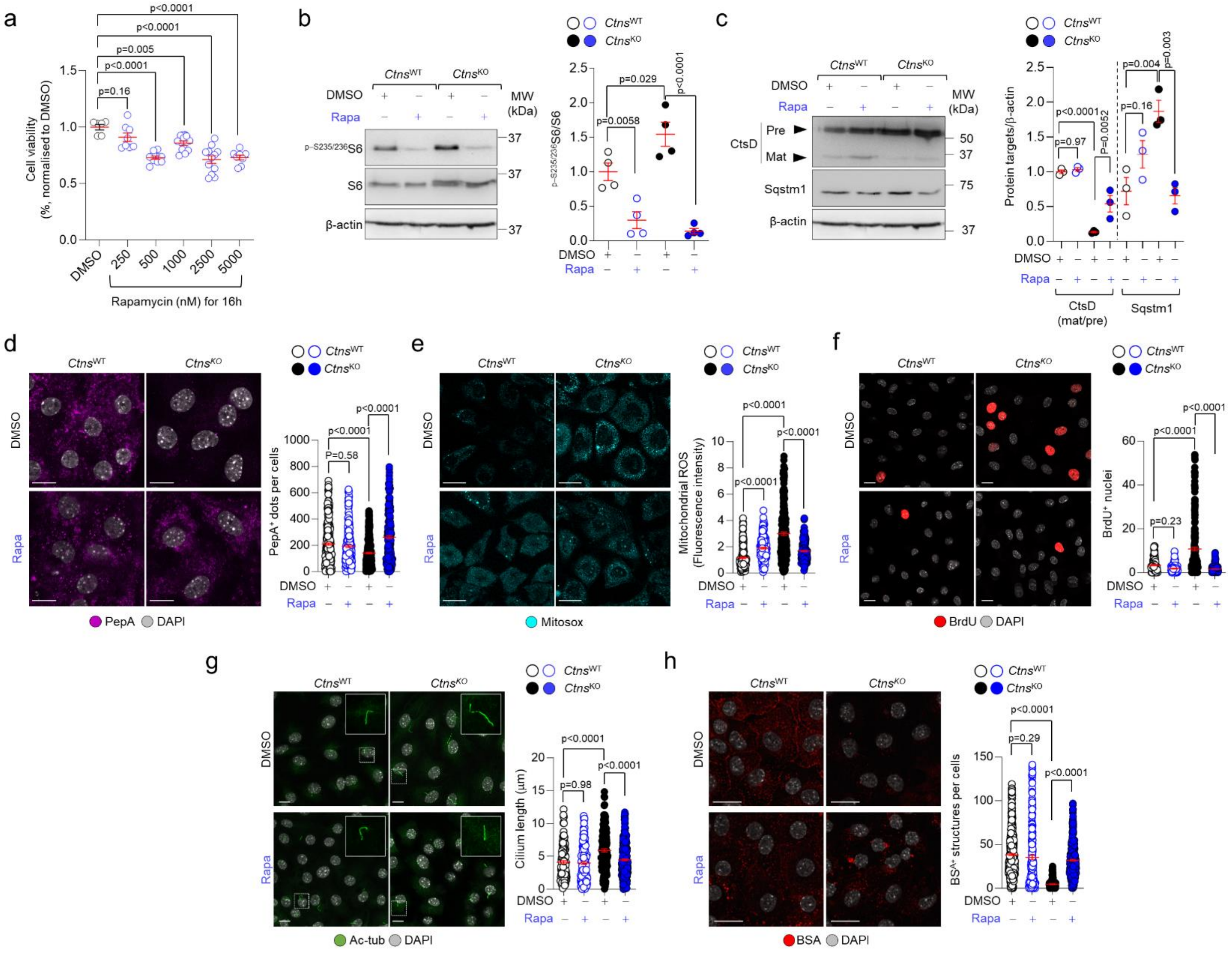
a
b
c
d
e
f
g
h
Cell viability (%, normalised to DMSO)
DMSO
250
500
1000
2500
5000
Rapamycin (nM) for 16h
p=0.16
p<0.0001
p=0.005
p<0.0001
p<0.0001
Ctns^WT
Ctns^KO
DMSO
Rapa
MW (kDa)
p-S235/236S6
S6
β-actin
p-S235/236S6/S6
p=0.0058
p=0.029
p<0.0001
CtsD
Pre
Mat
Sqstm1
Protein targets/β-actin
p<0.0001
p=0.97
p=0.0052
p=0.004
p=0.16
p=0.003
CtsD (mat/pre)
PepA⁺ dots per cells
P=0.58
PepA
DAPI
Mitochondrial ROS (Fluorescence intensity)
Mitosox
BrdU⁺ nuclei
p=0.23
BrdU
Cilium length (µm)
p=0.98
Ac-tub
BSA⁺ structures per cells
p=0.29
BSA

**Supplementary Figure 11. Rapamycin restores lysosome function and improves PT cell differentiation downstream of cystine storage.** (**a**) MTT assay measuring cell viability in Rapamycin-treated mPTCs. The cell viability has been normalized over the mean of the corresponding DMSO control (n>6 wells pooled from two biologically independent experiments). (**b**-**h**) The *Ctns* mPTCs were treated with low and non-toxic concentration of Rapamycin (250nm) for 16h.**(b**-**c**) Immunoblotting and quantification of the indicated proteins and phosphoproteins, n>3 biologically independent experiments. (**d**) The cells were pulsed for 1h with Pepstatin A (PepA;1µM at 37°C, pink) and analysed by confocal microscopy. Representative confocal images and quantification of the number of PepstatinA$^+$ structures per cell (n>221 cells per each condition). (**e**) Confocal microscopy and quantification of MitoSOX fluorescence intensity per cell (n>126 cells per each condition). (**f**) Confocal microscopy and quantification of BrdU$^+$ cells (expressed as percentage of total cells; n>112 randomly selected fields of views per each condition). (**g**) Maximum intensity projection of image stacks and quantification of the primary cilia length (Ac-tubulin, green) in *Ctns* mPTCs; n>132 cells pooled from 3 biologically independent experiments). (**h**) Cells were pulsed for 15min with Alexa633-labelled BSA and analyzed by confocal microscopy. Representative confocal images and quantification of the number of BSA-positive structures per cell; n>130 cells per each condition. Plots represent mean ± SEM. All values were pooled from three biologically independent experiments in **d**, **e**, **f**, **g** and **h**. Nuclei counterstained with DAPI (grey). Statistics calculated by one-way ANOVA followed Dunnett's multiple comparisons test in **a, b** and by one-way ANOVA followed by Sidak's or Tukey's comparisons multiple comparison test in **c**, **d**, e, **f**, **g,** and **h**. Scale bars are 10µm.

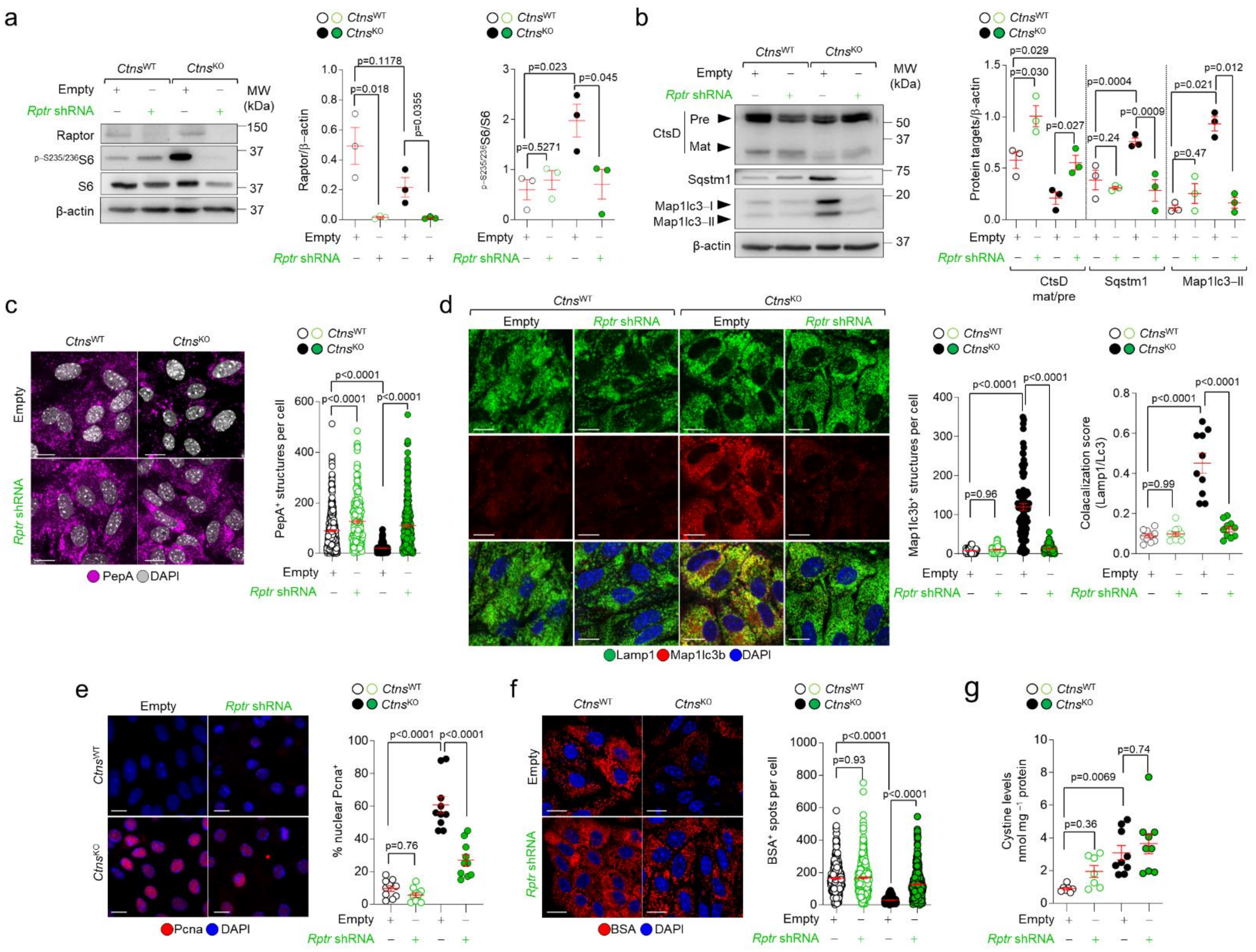

**Supplementary Figure 12.** ***Raptor*** **knockdown rescues lysosome, growth and differentiation defects in the tubular cells lacking CTNS and accumulating cystine.** (**a-g**) Cells were transduced with adenovirus particles bearing Empty or *Raptor*-shRNA (*Rptr* shRNA) for 5 days. (**a-b**) Immunoblotting and quantification of the indicated proteins and phosphoproteins, n=3 biologically independent experiments. (**c**) The transduced *Ctns* mPTCs were pulsed for 1h with Pepstatin A (PepA;1μM at 37°C, pink) and analysed by confocal microscopy. Representative confocal images and quantification of the number of PepstatinA-positive structures per cell; n>256 cells per each condition. (**d**) Representative confocal images and quantification of the number of Map1Lc3b-positive structures per cell and Map1Lc3b-Lamp1 colocalization (Map1Lc3b$^{+}$ structures: n>90 cells; colocalization score: n=10 randomly selected and non-overlapping fields of views per each condition). (**e**) Representative confocal images and quantification of the number of Pcna$^{+}$ nuclei (n=10 randomly selected and non-overlapping fields of views per each condition). (**f**) Cells were pulsed for 15min with Alexa633-labelled BSA and analyzed by confocal microscopy. Representative confocal images and quantification of the number of BSA-positive structures per cell; n>199 cells per each condition. (**g**) Quantification of cystine levels by HPLC (n>7 biologically independent samples per each condition). Plots represent mean ± SEM. All values were pooled from three biologically independent experiments in **c**, **d**, **e**, **f,** and **g**. Nuclei counterstained with DAPI (blue or grey). Statistics calculate by unpaired two-tailed Student's *t* test in **a** and **b,** and by one-way ANOVA followed by Sidak's or Tukey's comparisons in **c**, **d**, **e**, **f,** and **g**. Scale bars are 10μm.

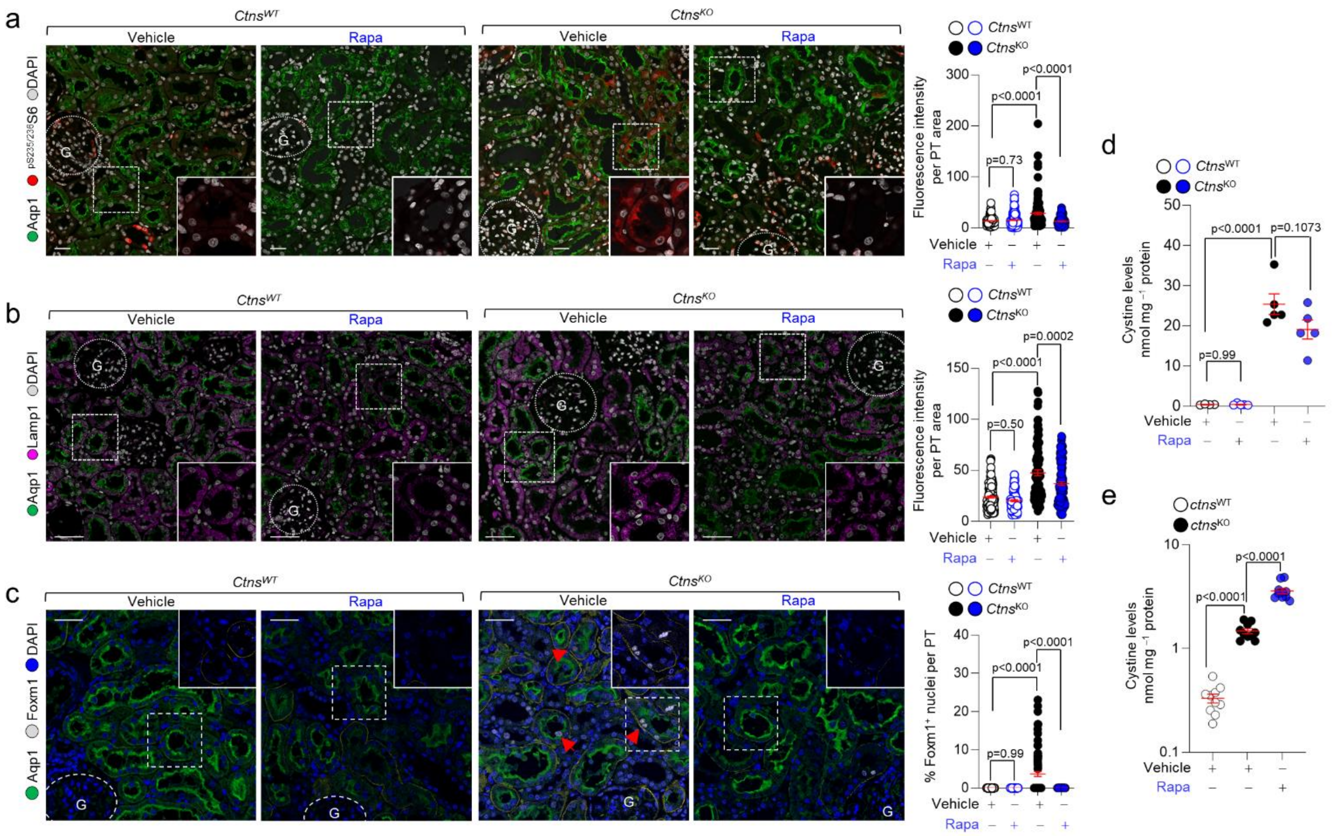

**Supplementary Figure 13. Effects of Rapamycin on storage and cellular alteration in the PT segments of *Ctns*-deficient rat kidneys and in zebrafish.** (**a**-**d**) *Ctns* rats at 12 weeks of age were subcutaneously implanted with a pellet providing a long-term release of rapamycin (Rapa, 1.5 mg/kg/day). After 2 weeks of treatment, the kidneys were harvested, immunostained for the indicated markers and analysed by confocal microscopy. (**a**-**c**) Representative confocal images and quantification of mean fluorescence intensity of (**a**) $^{pS235/236}$S6 (red) or (**b**) Lamp1 (pink) or (**c**) Foxm1 signals (gray) in Aqp1-positive PT segments (green) of *Ctns* rat kidneys; n>86 mouse PTs pooled from 4 animals per each group. (**f**) Quantification of cystine levels by HPLC; n=5 animals per each condition. (**e**) 5dpf-*ctns* zebrafish larvae were treated with vehicle or rapamycin (200nM) for 9 days. Quantification of cystine levels by HPLC, with each point representing a pool of seven zebrafish. Statistics calculate by one-way ANOVA followed by Tukey's multiple comparison test in **a, b**, cand **d**, and by ANOVA followed by Holm-Sidak's multiple comparisons test in **e.** Scale bars are 50μm.

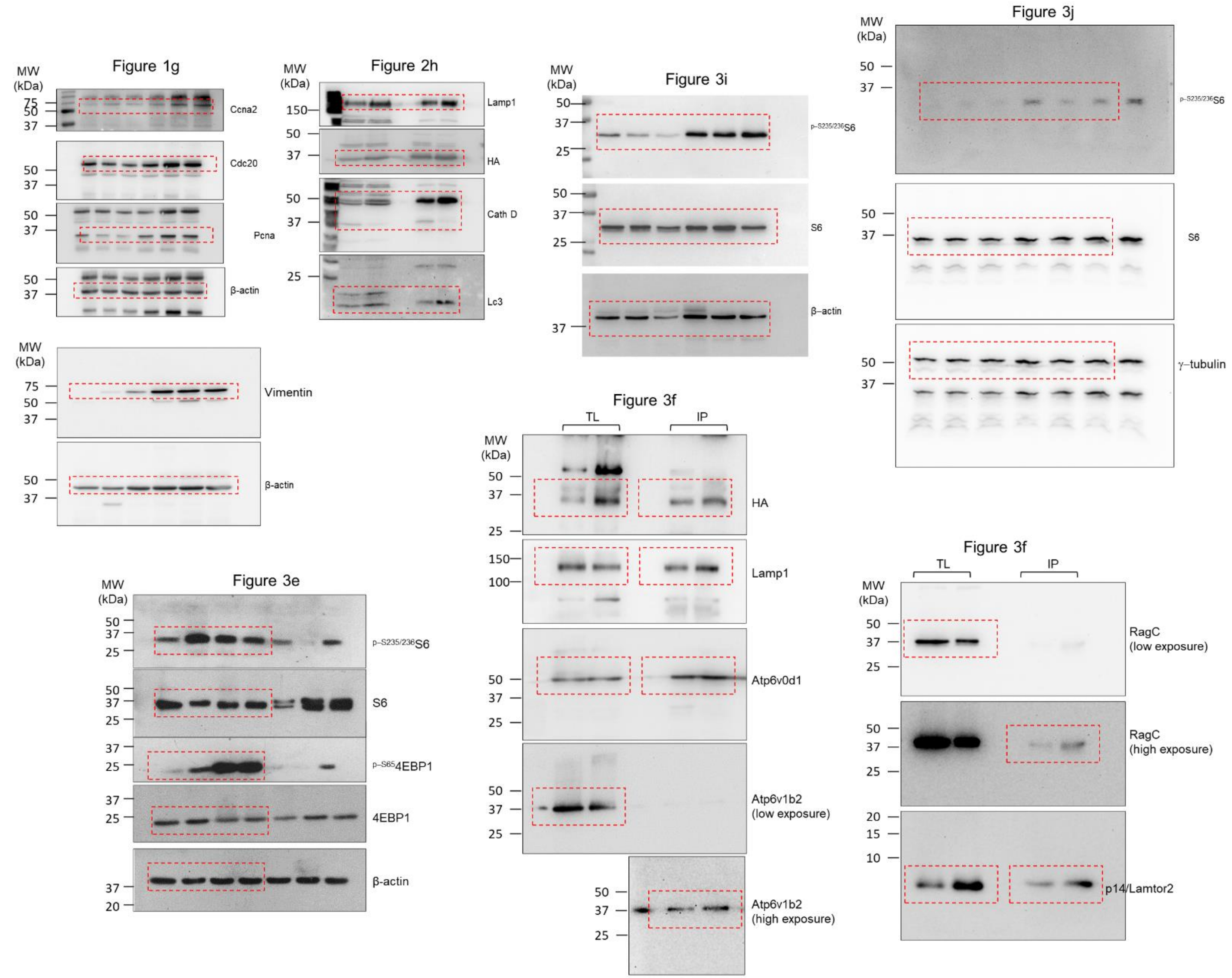


**Supplementary Figure 14.** Uncropped and unprocessed versions of western blots presented in the main figures

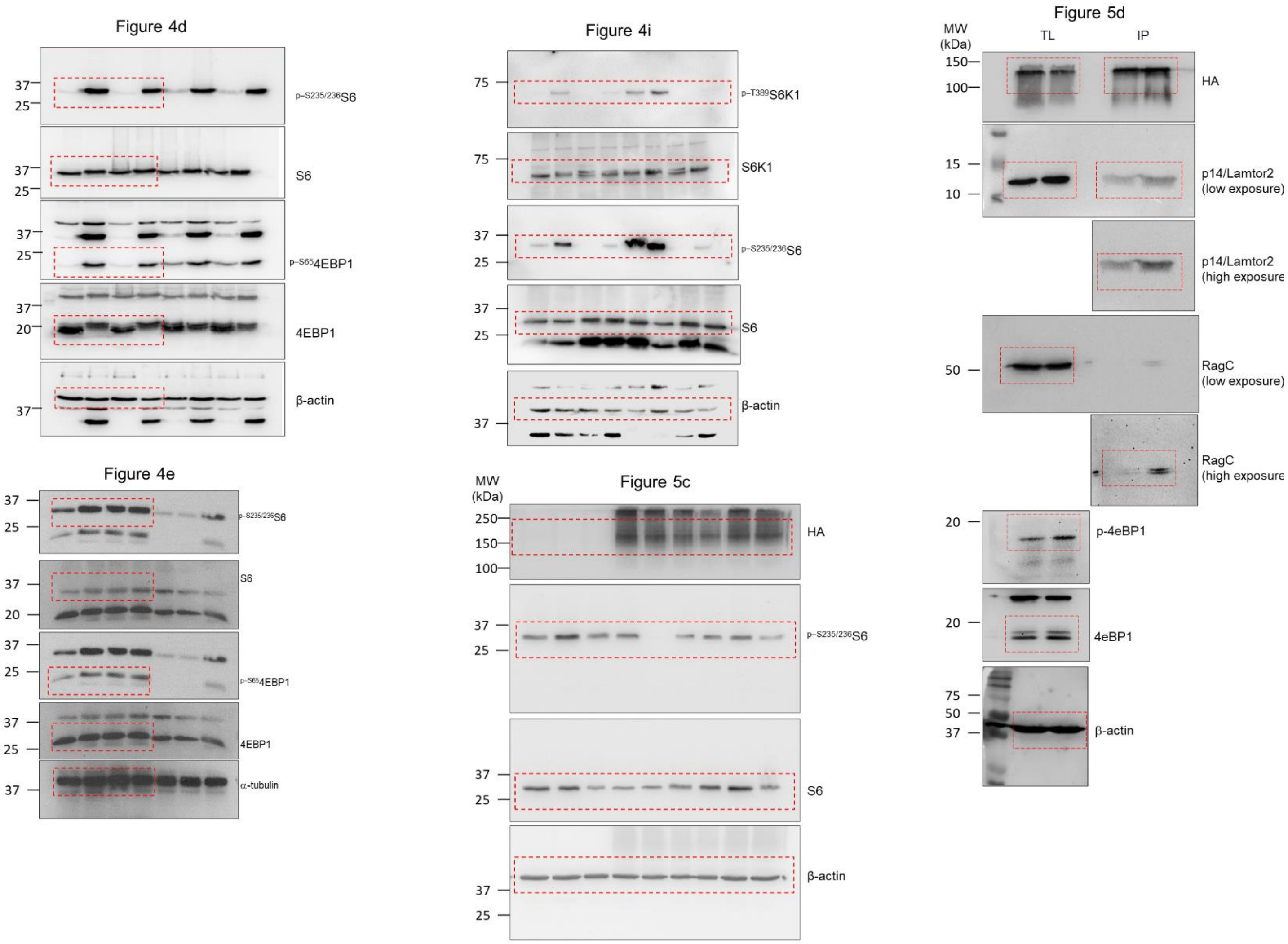


**Supplementary Figure 14 (continued)**. Uncropped and unprocessed versions of western blots presented in the main figures

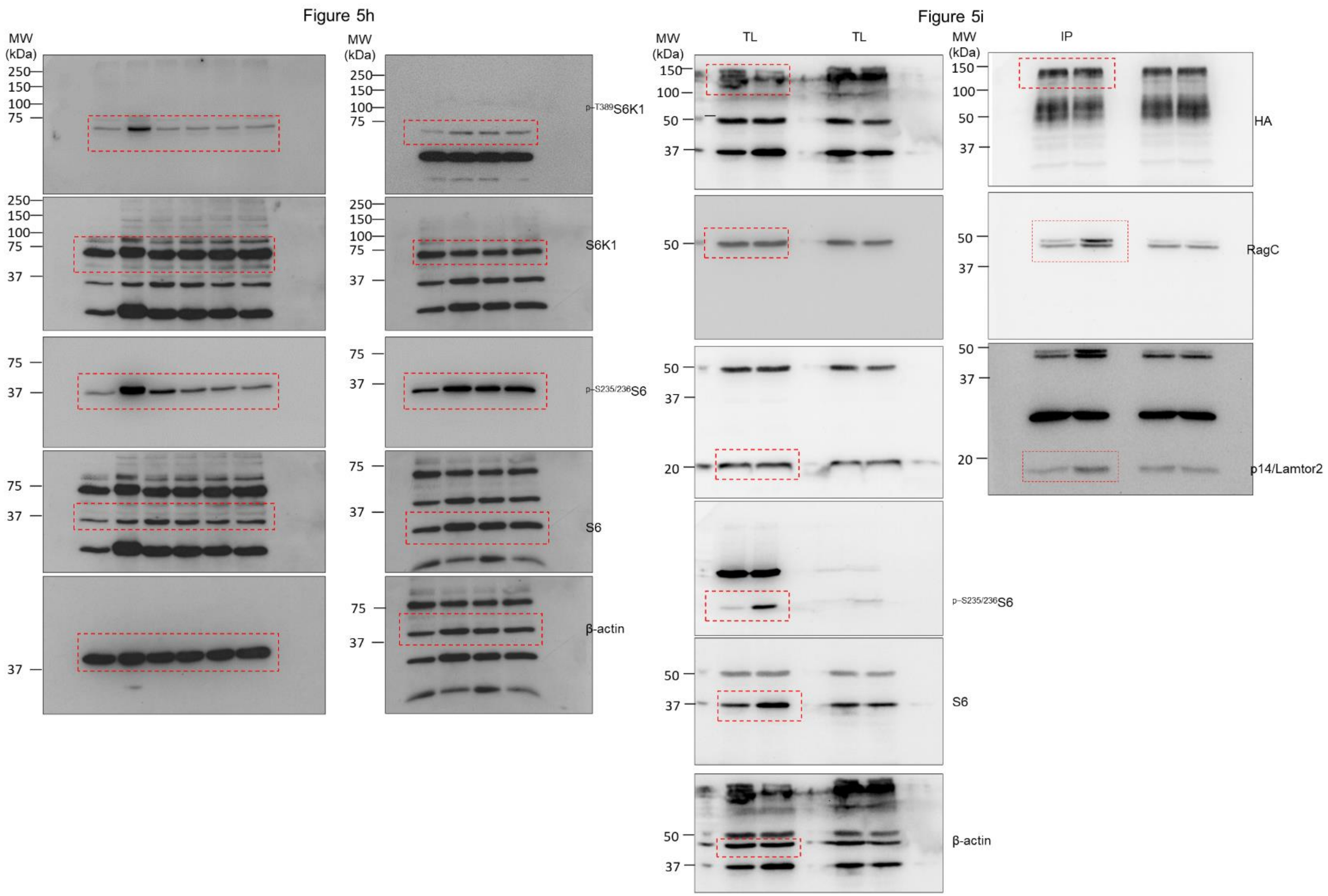


**Supplementary Figure 14 (continued).** Uncropped and unprocessed versions of western blots presented in the main figures

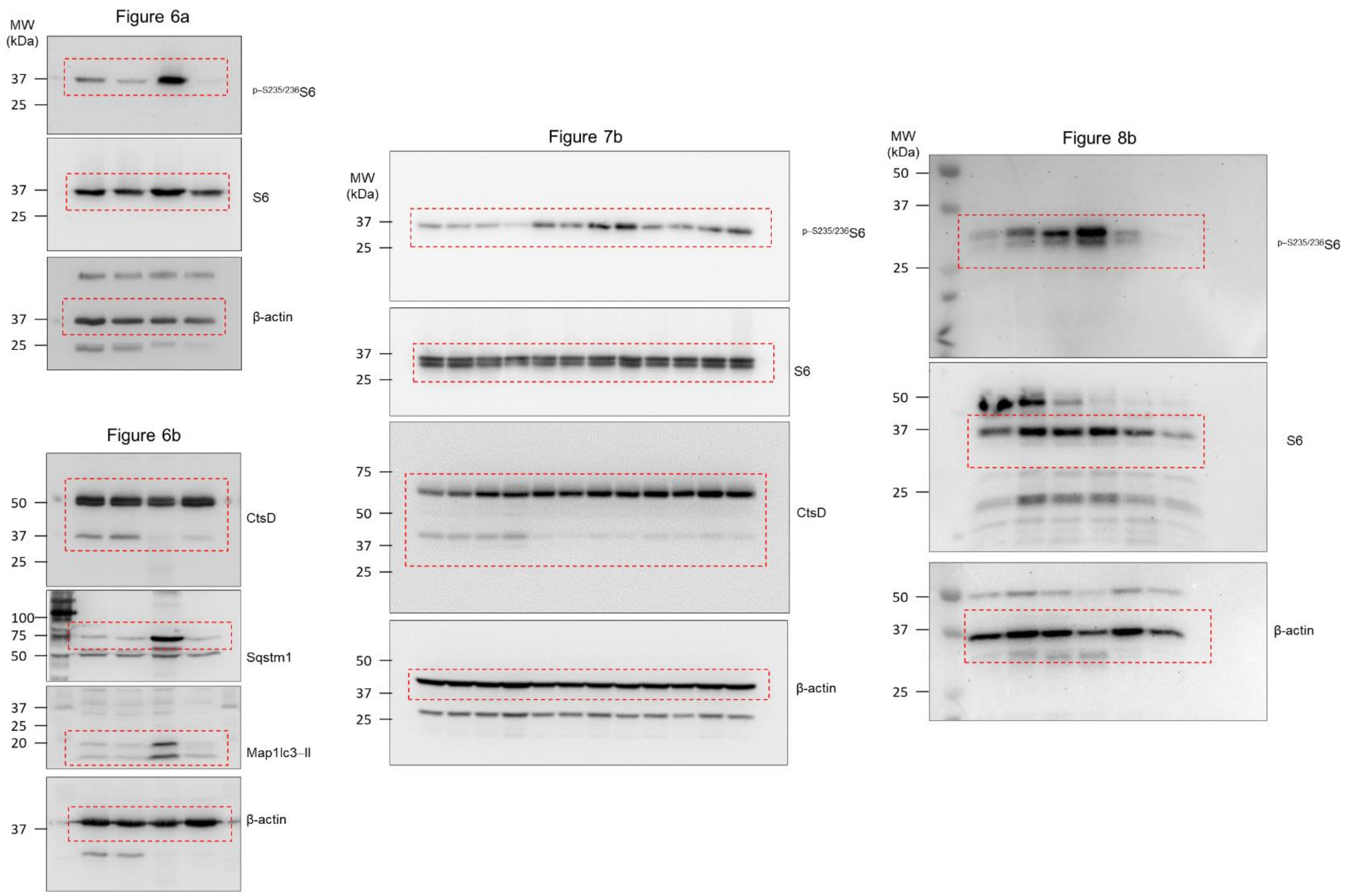


**Supplementary Figure 14 (continued).** Uncropped and unprocessed versions of western blots presented in the main figures

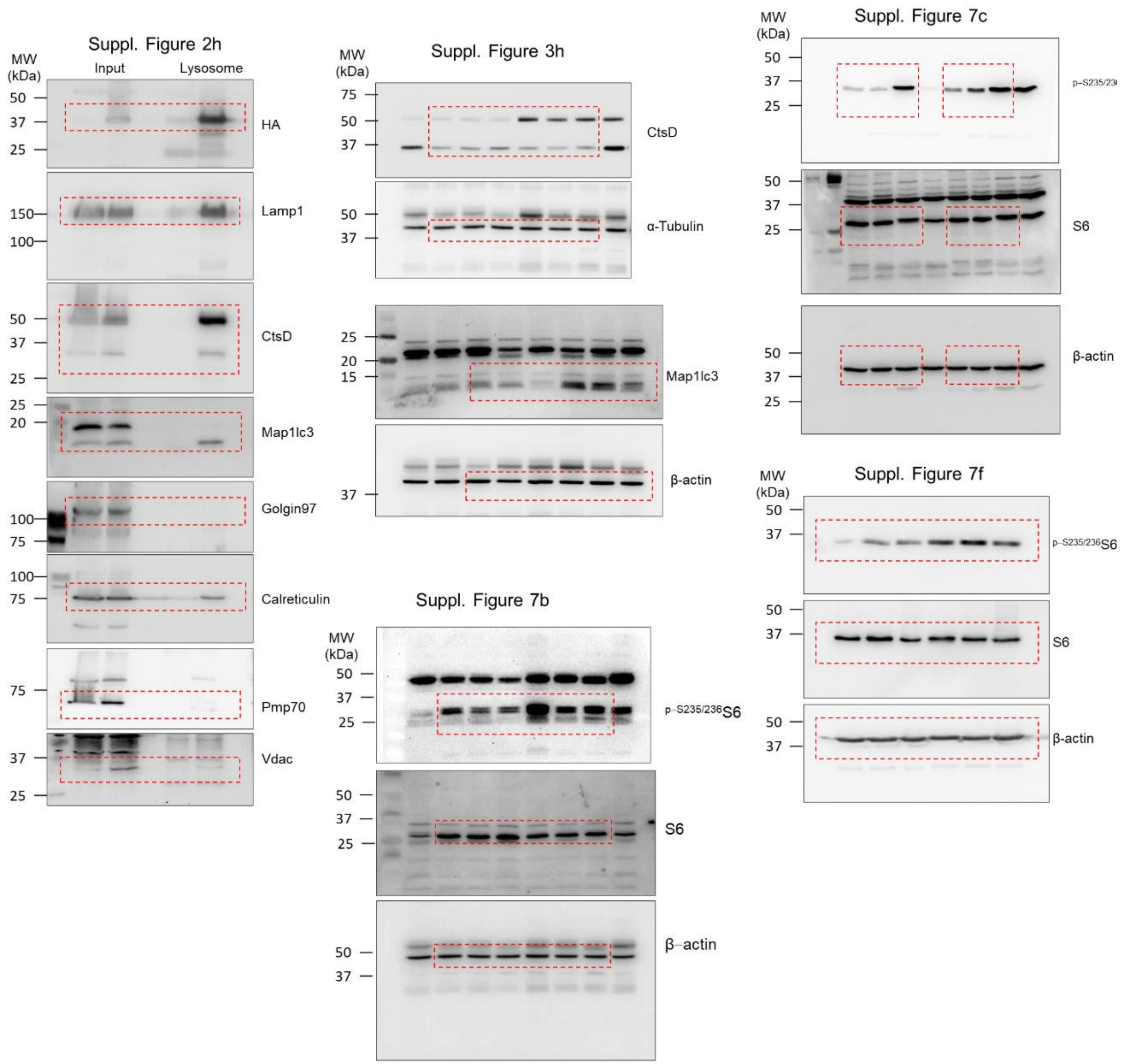


**Supplementary Figure 14 (continued).** Uncropped and unprocessed versions of western blots presented in the main and supplementary figures

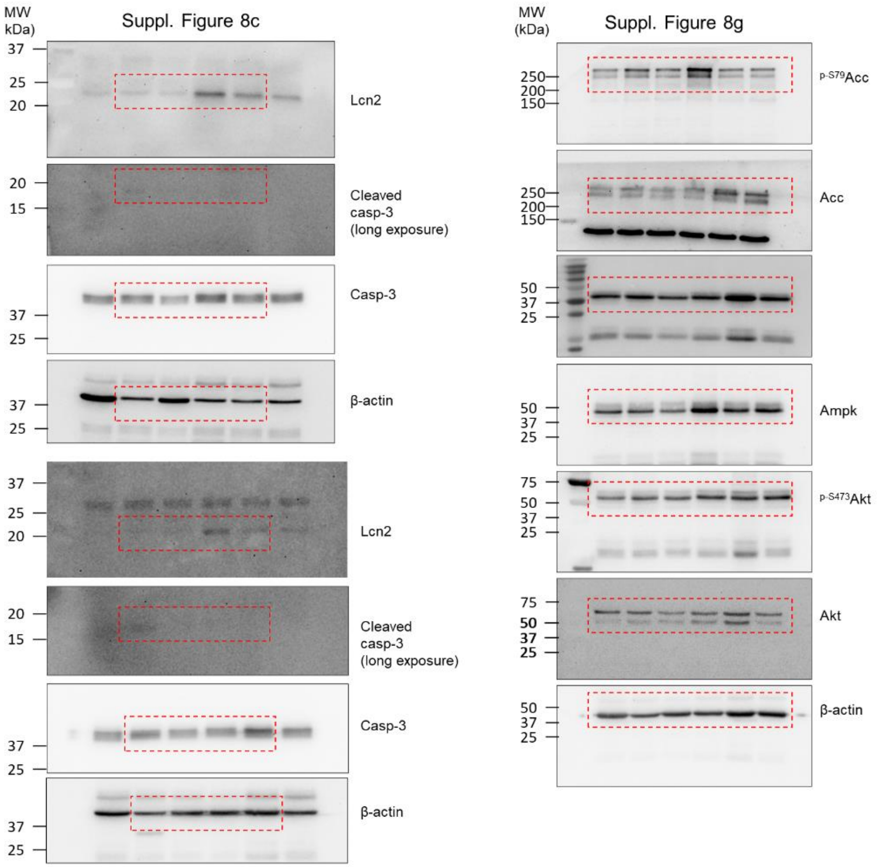


**Supplementary Figure 14 (continued).** Uncropped and unprocessed versions of western blots presented in the supplementary figures

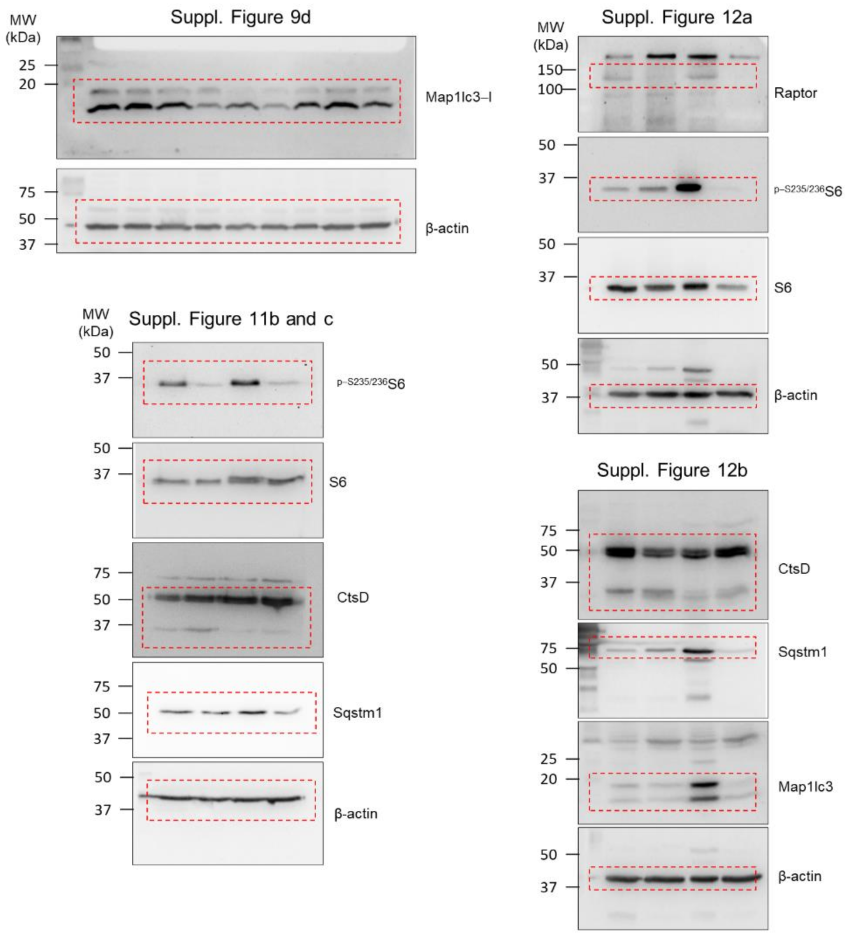


**Supplementary Figure 14 (continued).** Uncropped and unprocessed versions of western blots presented in the supplementary figures

**Supplementary Table 1.** Mouse primer pairs used for the gene expression analysis

| Gene name | Forward primer (5'-3') | Reverse primer (5'-3') |
|---|---|---|
| *Gapdh* | TGCACCACCAACTGCTTAGC | GGATGCAGGGATGATGTTCT |
| *Cdk1* | ACTCCACTCCGGTTGACATC | TCCACTTGGGAAAGGTGTTC |
| *Ccna2* | CTTGGCTGCACCAACAGTAA | AGCAATGAGTGAAGGCAGGT |
| *Cdc20* | ATGGAGCAGCCTGGAGACTA | GCTTACTCGAGCGGAGTGAC |
| *Ccnb2* | TGAAACCAGTGCAGATGGAG | CTGCAGAGCTGAGGGTTCTC |
| *Cdh2* | AGGGTGGACGTCATTGTAGC | TGTGACTAGCCCATCATTGC |
| *Tfr* | GCGCATTCAAGTGTCTGAAA | GAGCCACAACAGCATGAGAA |
| *F5* | TCCCGAGATATTCACGTGGT | TGCCAGCTACCTGGTTTTCT |
| *Apob* | CTCTTGCCACAGCTGATTGA | TGAGGGATTTGGGATCAGAG |
| *Lrp2* | CAGTGGATTGGGTAGCAG | GCTTGGGGTCAACAACGATA |
| *Tfe3* | TCTTCATCACGGGTCTTGCT | TTCTCGAGGTGGGTCTGAAC |
| *Mcoln1* | TCCACTTCCAGCTGAAGACA | CTGGATGTGGGTCTTGGTCT |
| *Sqstm1* | CCCCAATGTGATCTGTGATG | AAGGGGTTGGGAAAGATGAG |
| *Map1Lc3B* | CCGAGAAGACCTTCAAGCAG | CCAGGAACTTGGTCTTCTCC |
| *Lamp1* | TAGTGCCCACATTCAGCATCTCCA | TTCCACAGACCCAAACCTGTCACT |
| *CtsD* | CCAAGTTTGATGGCATCTTGGGCA | TGGAGTCAGTGCCACCAAGCATTA |
| *Atg9A* | CTCCGAGTGATTCTTGCACA | GACGATGGGACTCAGCAACT |
| *Foxm1* | AAGGCAAAGACAGGAGAGCT | AGGGCTCCTCAACCTTAACC |
| *Sox9* | CAAGAACAAGCCACACGTCA | GTGGTCTTTCTTGTGCTGCA |
| *Vimentin* | AATGCTTCTCTGGCACGTCT | AGTGAGGTCAGGCTTGGAAA |
| *Slc7a13* | GCCTTTGTTGTTTTGTGCCC | GGGCTCTTGGTACCTCAGTT |
| *Slc5a2* | TTGGGCATCACCATGATTTA | GCTCCCAGGTATTTGTCGAA |
| *Slc34a1* | GGGAGAAGCTATCCAGCTCA | ACAGCAAACCAGCGGTACTT |
| Slc3a1 | ACTCAGGTGGGAATGCATGA | CGGCTTCCTGGATAAATGGC |

**Supplementary Table 2.** Rat primer pairs used for the gene expression analysis

| Gene name | Forward primer (5'-3') | Reverse primer (5'-3') |
|---|---|---|
| *Slc34a1* | GCCCGAGATGTTGAAGAAGA | TCAGTATTGAGCGGGCCTAC |
| *Slc5a2* | TCTTTGTGCCCCGTGTTAAT | AGTGTACCCGGCAGAAGATG |
| *Foxm*1 | AAGGCAAAGACAGGAGAGCT | CTGCCAAGACACTGATGCTC |
| *Vimentin* | TGCCCTTGAAGCTGCTAACT | AATCCTGCTCTCCCCTT |
